# Sexual life cycle establishes the unicellular red algae Cyanidiophyceae as a genetically tractable model lineage for eukaryotic evolution

**DOI:** 10.1101/2025.10.15.681507

**Authors:** Shunsuke Hirooka, Takayuki Fujiwara, Mark Seger, Soichi Inagaki, Shota Yamashita, Dai Tsujino, Ryo Onuma, Yu Kanesaki, Satoru Watanabe, Yuu Hirose, Ryudo Ohbayashi, Mari Takusagawa, Baifeng Zhou, Reiko Tomita, Fumi Yagisawa, Peter Lammers, Atsuko H. Iwane, Shin-ya Miyagishima

## Abstract

The thermo-acidophilic unicellular algal class Cyanidiophyceae is the earliest-branching lineage in red algae, which diverged from Viridiplantae lineage (green algae and land plants) soon after chloroplast establishment in the common ancestor of Archaeplastida. Cyanidiophyceae possess extremely simple genomes (8.7–17.8 Mb; approximately 4,800– 7,800 genes), and the cell-wall-less, genetically tractable strain *Cyanidioschyzon merolae* 10D has served as a model organism. However, its unknown sexual life cycle has limited its utility in studies of evolution and genetics. Inspired by the recent discovery of sexual reproduction in the cyanidiophycean genus *Galdieria*, we identified similar life cycles in the other cyanidiophycean genera *Cyanidioschyzon*, *Cyanidiococcus*, and *Cyanidium*. In these genera, the cell-walled diploid form, exclusively observed in nature, produces a cell-wall-less haploid form when the culture pH is lowered, and both proliferate asexually. In addition, the cell-wall-less *Cyanidioschyzon merolae* 10D strain has been shown to be a haploid clone that forms a cell-walled diploid through mating with other haploid clones. Building on these findings, we generated high-quality genomic resources with phase-specific transcriptomes and developed genetic manipulation systems using the cell-wall-less haploids of these genera. We further uncovered phase-specific distribution of histone H3 lysine 27 trimethylation linked to haploid- and diploid-specific gene expression, including transcription factors involved in differentiation associated with sexual reproduction in plants. Additionally, biparental inheritance of organelle DNA occurs following isogamous mating of haploid cells but resolves into uniparental inheritance during diploid proliferation. These advances position Cyanidiophyceae as a powerful model lineage for studying early Archaeplastida evolution, the shared mechanisms of photosynthetic eukaryotes, and their environmental adaptation.

## Introduction

Eukaryotes have evolved and diversified over more than 2 billion years from a last eukaryotic common ancestor (LECA) that arose through a merger between an archaeon-derived host cell and an α-proteobacterial endosymbiont that ultimately became the mitochondrion (Richards et al., 2024). After LECA diverged into the ancestors of several eukaryotic supergroups, the common ancestor of the supergroup Archaeplastida acquired a chloroplast through the genetic integration of a cyanobacterial endosymbiont into the host nuclear genome, in a process known as primary endosymbiosis. This lineage subsequently diverged into Rhodophyta (red algae), Glaucophyta, and Viridiplantae (green algae and land plants) (Fig. 1A) (Bowles et al., 2023). In addition to Archaeplastida, many other eukaryotic lineages also acquired photosynthetic capabilities through secondary endosymbiosis, by genetically integrating unicellular red or green algal endosymbionts into their nuclear genomes. In particular, red algae contributed to the establishment of secondary chloroplasts (plastids) in various eukaryotic groups such as Stramenopila, Alveolata, Haptista, and Cryptista (Sibbald and Archibald, 2020; Strassert et al., 2021).

**Figure 1.**
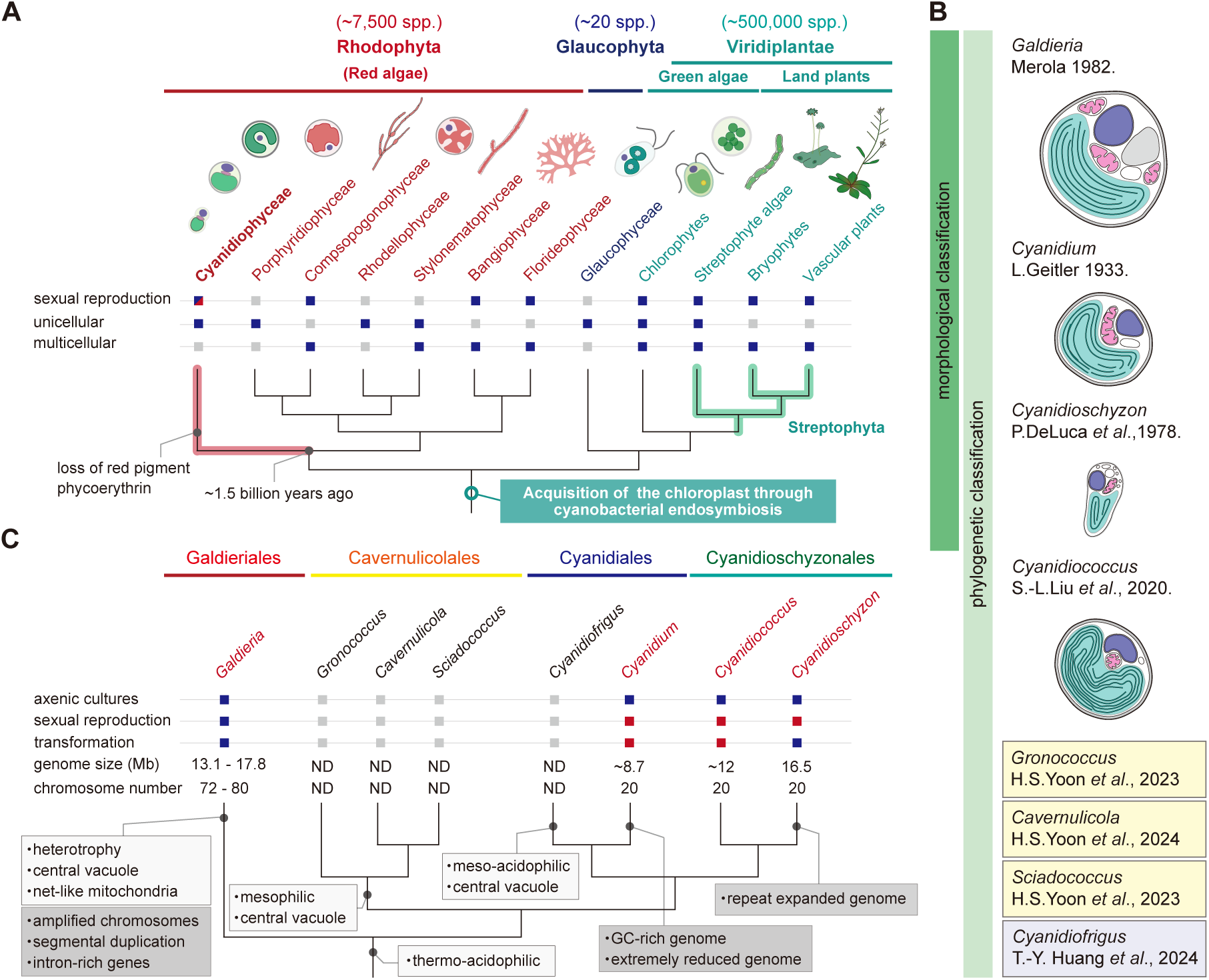
Taxonomy of Cyanidiophyceae. **A)** Evolutionary position of the unicellular red algae Cyanidiophyceae within Archaeplastida. The cladogram, based on previous studies (Munoz-Gomez et al., 2017; One Thousand Plant Transcriptomes, 2019), indicates whether sexual reproduction has been observed and whether the respective groups are unicellular or multicellular. Blue boxes indicate presence based on previous studies, gray boxes indicate absence (or no reports), and red boxes indicate presence based on both a previous study (Hirooka et al., 2022) and this study. **B)** Cyanidiophycean algae were originally classified into three genera—*Galdieria*, *Cyanidium*, and *Cyanidioschyzon*—based on morphological and physiological characteristics (Merola et al., 1981). Subsequently, a fourth genus, *Cyanidiococcus*, was described primarily on the basis of molecular phylogenetic analysis, along with morphological, physiological, and genomic traits (Liu et al., 2020). More recently, four additional genera— *Gronococcus*, *Cavernulicola*, *Sciadococcus*, and *Cyanidiofrigus*—have been described (Park et al., 2023; Huang et al., 2024). Illustrations of *Galdieria*, *Cyanidium*, *Cyanidioschyzon*, and *Cyanidiococcus*, created with reference to Merola et al. (1981) and Liu et al. (2020), are shown. **C)** The cladogram, based on a previous study (Park et al., 2023; Huang et al., 2024), shows the intergeneric phylogenetic relationships of Cyanidiophyceae, comprising four major clades: Galdieriales, Cavernulicolales, Cyanidiales, and Cyanidioschyzonales. Each clade is annotated with captions indicating traits (light gray) and genomic features (dark gray). Also shown are the availability of axenic cultures, the observation of sexual reproduction, genetic tractability, genome size, and chromosome number. Blue boxes indicate presence based on previous studies, gray boxes indicate absence (or no observations), and red boxes indicate presence based on this study.

Several lines of evidence indicate that, in addition to the mitochondrion, LECA already possessed complex eukaryotic features, including several functionally distinct membrane-bound organelles (such as the nuclear envelope, endoplasmic reticulum, peroxisomes, lysosomes, and Golgi apparatus), sophisticated cytoskeletal systems, and a sexual reproduction system (Richards et al., 2024). However, during the subsequent evolution and diversification of eukaryotes, each lineage independently developed a range of unique traits, including variations in cell morphology, organelle structures, motility, sex determination, and environmental adaptation (Grosberg and Strathmann, 2007; Beukeboom and Perrin, 2014; Adl et al., 2019; Fritz-Laylin and Titus, 2023). For example, among Archaeplastida, red algae, certain lineages of green algae, and land plants have each independently evolved multicellular sexual life cycles (Fig. 1A) (Bowles et al., 2023). To understand how such traits evolved, it is necessary to conduct not only phenotypic and genotypic comparisons across different lineages, but also a variety of comparative functional analyses, including genetic manipulation in model organisms (Fields and Johnston, 2005).

Nevertheless, when the distribution of available model organisms is considered in light of the phylogenetic diversity of eukaryotes, a substantial bias becomes evident (Hedges, 2002; Adl et al., 2019). As a result, our current understanding of eukaryotic evolution remains largely incomplete.

Regarding the situation in Archaeplastida, after the establishment of some angiosperms such as *Arabidopsis thaliana* as model organisms (Hedges, 2002) (vascular plants in Fig. 1A), the moss *Physcomitrium patens* (Rensing et al., 2020) and the liverwort *Marchantia polymorpha* (Kohchi et al., 2021) were later developed and have been used as model systems representing basal land plants to study the evolutionary processes underlying key traits of land plants, especially development and sexual reproduction (bryophytes in Fig. 1A). Tracing further back in the phylogeny, the unicellular green alga *Chlamydomonas reinhardtii* has been extensively studied as a model green alga (Goodenough, 2022) (chlorophytes in Fig. 1A). In addition, the green algal order Volvocales, which includes unicellular genus *Chlamydomonas* and multicellular genera with varying cell numbers—such as *Gonium* (16 cells; genetically tractable) (Hanschen et al., 2016) and *Volvox* (over 500 cells; genetically tractable) (Prochnik et al., 2010)—represents a lineage in which multicellularity evolved independently of land plants, and has served as a model lineage to study the evolution of multicellularity and its coordination with sexual reproduction (chlorophytes in Fig. 1A).

Recently, a procedure for genetic manipulation was developed in *Closterium* (Kawai et al., 2022), a member of the class Zygnematophyceae, which is expected to serve as a unicellular algal model that is evolutionarily closer to land plants than Volvocales or other genetically tractable green algae such as *Ostreococcus* (Lozano et al., 2014), thus filling the gap between core green algae and land plants (streptophyte algae in Fig. 1A). Tracing further back in the phylogeny, the unicellular red alga *Cyanidioschyzon merolae* (*Cz. merolae*), which belongs to the class Cyanidiophyceae (Fig. 1A), is equipped with advanced genetic manipulation systems, and research has begun using this organism to understand the early evolution of the Archaeplastida (Kuroiwa et al., 2018; Miyagishima and Tanaka, 2021), as described below.

However, because no sexual reproduction process has been identified in this alga, attempts to understand the early evolution of the sexual life cycle in Archaeplastida inevitably rely on comparisons with organisms outside Archaeplastida, such as model animals and yeasts belonging to the supergroup Opisthokonta (Adl et al., 2019).

The unicellular red algae Cyanidiophyceae primarily inhabit thermo-acidic environments (pH 0.05 to 5.0, <56°C) in sulfuric hot springs of volcanic areas worldwide and exhibit a blue-green color because they have lost the red pigment phycoerythrin during evolution, which is present in other red algae (Seckbach, 1994). This group is estimated to have branched off from a mesophilic and neutrophilic common ancestor of other unicellular and multicellular red algal lineages early in eukaryotic evolution, approximately 1.5 billion years ago (Yoon et al., 2004). The class Cyanidiophyceae includes a total of eight genera, with only four being axenically cultured: *Galdieria*, *Cyanidium*, *Cyanidiococcus*, and *Cyanidioschyzon*, based originally on morphological and physiological characteristics(Merola et al., 1981), and more recently expanded to also include genomic sequences and structures (Liu et al., 2020; Cho et al., 2023) (Fig. 1B). *Galdieria* cells are spherical, 3–11 μm in diameter, enclosed by a thick cell wall, and contain a single multi-lobed chloroplast, a net-like mitochondrion, and a central vacuole (Albertano et al., 2000; Yamashita et al., 2025). They proliferate by undergoing two to five rounds of successive cell divisions, generating 4 to 32 daughter cells before hatching from the mother cell (Albertano et al., 2000; Jong et al., 2021). Among Cyanidiophyceae, *Galdieria* is unique in its ability to grow photoautotrophically, mixotrophically, and heterotrophically, utilizing more than 50 different carbon sources, unlike other genera which are obligate photoautotrophs (Gross and Schnarrenberger, 1995; Barbier et al., 2005). Cells of *Cyanidium* and *Cyanidiococcus*, which are spherical (2–5 μm in diameter) and enclosed by a rigid cell wall, are morphologically indistinguishable by optical microscopy, although their genome sizes differ significantly (Fig. 1B and 1C) (Cho et al., 2023). These genera proliferate by forming four daughter cells through two successive divisions before being released from the mother cell (Merola et al., 1981; Jong et al., 2021).

In contrast, *Cyanidioschyzon* cells are oval-shaped (1.5–3.5 μm in length), lack a cell wall, and proliferate by binary fission (Fig. 1B) (Merola et al., 1981). Unlike *Galdieria*, these three genera contain a cup-shaped chloroplast, a single disc-shaped mitochondrion, and a few small vacuoles (Merola et al., 1981; Albertano et al., 2000). In addition to the above-mentioned thermo-acidophilic genera, the class Cyanidiophyceae contains four mesophilic genera that have not yet been axenically cultured: *Gronococcus*, *Cavernulicola*, and *Sciadococcus*, which form a monophyletic clade and thrive at moderate temperatures (18–25 °C), neutral pH, and low light in a few coastal caves (Fig. 1B and 1C) (Park et al., 2023); and another genus, *Cyanidiofrigus*, which is mainly found in non-aquatic microhabitats at moderate temperatures (20–30 °C) and low pH conditions (pH 0.5 to 2.5) (Fig. 1B and 1C) (Huang et al., 2024).

Cells of these mesophilic genera possess a cell wall and a central vacuole, proliferate by forming four daughter cells within a mother cell (Park et al., 2023; Huang et al., 2024). Furthermore, *G. phlegrea* Soos, a representative of the crypto-endolithic species belonging to one of the two major clades of *Galdieria* (Qiu et al., 2013), was isolated from soil samples in Soos National Park in the Czech Republic and exhibits a relatively low optimum growth temperature of ∼30 °C (Gross et al., 2002). These mesophilic genera are thought to have secondarily adapted to mesophilic environments from thermo-acidic environments.

In addition to the characteristics mentioned above—adaptation to thermo-acidic environments and simple intracellular architecture—Cyanidiophyceae are also notable for possessing one of the most compact nuclear genomes among eukaryotes (8.7 to 17.8 Mb; approximately 4,800 to 7,800 protein-coding genes) (Matsuzaki et al., 2004; Nozaki et al., 2007; Qiu et al., 2013; Schonknecht et al., 2013; Rossoni et al., 2019; Liu et al., 2020; Downing et al., 2022; Hirooka et al., 2022; Cho et al., 2023). Their compact genomes are attributed to two phases of significant gene loss: one that occurred in the common ancestor of Rhodophyta, and another in the common ancestor of Cyanidiophyceae (Qiu et al., 2015).

Although the specific causes of these reductions remain unclear, both events may have been driven by the need to reduce energy consumption—as an adaptation to oligotrophic environments in the former case, and in the latter, to reallocate the energy saved through genome reduction toward coping with environmental stress such as high temperature and low pH (Van Etten et al., 2023). Although the genomes of Cyanidiophyceae are streamlined, they have also acquired ∼1% of their gene inventory through horizontal gene transfer (HGT) from bacteria and archaea, and the majority of these HGT-derived genes are apparently related to adaptation to extreme environments (Schonknecht et al., 2013; Rossoni et al., 2019). Thus, Cyanidiophyceae, comprising eight genera (Fig. 1C), could serve as an excellent model lineage for understanding the evolution of Archaeplastida and, more broadly, eukaryotes.

Furthermore, comparisons among genera will not only help understand the traits and mechanisms possessed by the common ancestor over one billion years ago, but also provide information to distinguish and analyze those that each genus or species has independently evolved.

Among Cyanidiophyceae, a strain of the genus *Cyanidioschyzon*, *Cz. merolae* 10D (16.5 Mb genome; 4,803 protein-coding genes [Matsuzaki et al., 2004; Nozaki et al., 2007] — this number will be revised later in this study), which was isolated from a mixture of algae in sulfuric hot spring water in Italy (Toda et al., 1995), has emerged as a model organism (Kuroiwa et al., 2018; Miyagishima and Tanaka, 2021). Owing to the absence of a cell wall — the presence of which hampers the introduction of exogenous DNA —and its highly efficient homologous recombination activity, arbitrary chromosomal loci in this strain can be precisely modified via polyethylene glycol (PEG)-mediated transformation (Ohnuma et al., 2008). Based on this technique, several molecular genetic procedures have been developed, including gene knockouts, epitope tagging (Imamura et al., 2009; Fujiwara et al., 2013), selectable marker recycling (Takemura et al., 2018; Bora and Tanaka, 2025), inducible gene expression (via heat shock, ammonium-to-nitrate exchange, IPTG, or estradiol) (Sumiya et al., 2014; Fujiwara et al., 2015; Fujiwara et al., in press), inducible protein knockdown (using rapamycin, IPTG, or estradiol) (Fujiwara et al., 2024; Fujiwara et al., in press), and CRISPR/Cas9-based genome editing (Tanaka et al., 2021). Taking advantage of these techniques and the simple cellular and genomic content of *Cz. merolae* 10D, the strain has increasingly been used in a wide range of research fields, including organelle division (Miyagishima et al., 2003; Nishida et al., 2003; Yoshida et al., 2010; Imoto et al., 2013), the cell cycle (Miyagishima et al., 2014; Fujiwara et al., 2020), the coordination of these two mechanisms (Miyagishima et al., 2012; Sumiya et al., 2016), the photosynthetic apparatus (Krupnik et al., 2013; Nilsson et al., 2014; Nikolova et al., 2017; Pham et al., 2019), epigenetics (Mikulski et al., 2017; Hisanaga et al., 2023), RNA processing (Soma et al., 2007; Stark et al., 2015), responses to environmental changes (Imamura et al., 2009; Kobayashi et al., 2014; Pancha et al., 2019), and industrial applications (Rahman et al., 2017; Hirooka et al., 2020; Yoshida et al., 2021; Seger et al., 2023; Villegas-Valencia et al., 2023). Thus, if present, uncovering sexual reproduction in *Cz. merolae* would greatly expand the range of its applications for understanding eukaryotic evolution and for studies based on genetics.

Regarding this point, we recently identified a sexual reproduction process in *Galdieria* (Hirooka et al., 2022), the earliest-diverging lineage within Cyanidiophyceae (Fig. 1C), which originated approximately one billion years ago (Yoon et al., 2004). In *Galdieria*, the previously recognized natural, cell-walled form (Fig. 1B) is diploid and, upon a decrease in the pH of the culture, produces a cell-wall-less haploid that is likewise capable of asexual reproduction via mitotic cell division (Hirooka et al., 2022). In addition, we successfully transferred the PEG-mediated genetic manipulation method developed in *Cz. merolae* 10D to *G. partita*, a species of the genus *Galdieria*, by utilizing the cell-wall-less haploid (Hirooka et al., 2022). Inspired by these findings and developments, we have explored the possibility that *Cz. merolae* and other members of Cyanidiophyceae also undergo a sexual reproduction process, establishing them as a model lineage equipped with functional genomics techniques for understanding eukaryotic evolution.

Here, we report the sexual life cycles of *Cyanidioschyzon* and two other genera of Cyanidiophyceae, *Cyanidiococcus* and *Cyanidium*, in which diploid cells are cell-walled and proliferate by forming four daughter cells within a mother cell, as has been observed in *Cyanidiococcus* and *Cyanidium* (Fig. 1B), while haploid cells are cell-wall-less and proliferate by binary fission, as has been observed in *Cz. merolae* 10D (Fig. 1B). In line with these findings, it is also shown that *Cz. merolae* 10D (isolated from Italy) is a haploid clone, which produces a cell-walled diploid through isogamous mating with a haploid of the opposite mating type (generated from a cell-walled diploid clone isolated from the USA).

Based on these findings, we have also established high-quality genomic resources for strains belonging to each genus of Cyanidiophyceae, including complete genome sequences based on haploids without allelic polymorphisms, gene models, and transcriptome datasets encompassing haploid- and diploid-specific genes. Furthermore, we have developed genetic manipulation techniques by utilizing the cell-wall-less haploids of the respective genera, thereby establishing Cyanidiophyceae as a genetically tractable model lineage. Furthermore, we found changes in the genomic distribution of tri-methylation of histone H3 lysine 27 (H3K27me3), a well-known repressive epigenetic mark, between haploid and diploid cells, which correlate with haploid- and diploid-dominant gene expression. These phase-dominant genes include transcription factors belonging to families that, in plants, are involved in diverse developmental processes, including flower formation and transitions in the sexual life cycle. In addition, we also observed initial biparental inheritance of mitochondrial and chloroplast DNA upon diploid formation through the mating of haploids, followed by the resolution of heteroplasmy (i.e., eventual uniparental inheritance) during diploid proliferation. In this way, Cyanidiophyceae—based on the developed datasets, procedures, and observations—has now become a powerful model lineage for understanding the early evolution of Archaeplastida, as well as the fundamental and shared mechanisms and environmental adaptation in photosynthetic eukaryotes.

## Results

### All members of the Cyanidiophyceae, including the genera *Galdieria*, *Cyanidium*, *Cyanidiococcus*, and *Cyanidioschyzon*, undergo sexual reproduction

The genetically tractable *Cz. merolae* was initially identified as the only cell-wall-less member of the Cyanidiophyceae that proliferates via binary fission and was described in 1978 (De Luca et al., 1978; Merola et al., 1982). However, since the 2000s, several strains possessing a cell wall—morphologically indistinguishable from *Cyanidium* and *Cyanidiococcus* under optical microscopy—have been isolated and assigned to the *Cyanidioschyzon* clade based on molecular phylogenetic analyses (Fig. 1C) (Toplin et al., 2008; Lee et al., 2015). Conversely, *Cyanidiococcus* possesses a cell wall and releases four daughter cells from a mother cell, similar to *Cyanidium* (Fig. 1B), but was recently established as a distinct genus that forms a sister group to *Cyanidioschyzon* (Cyanidioschyzonales in Fig. 1C), rather than to *Cyanidium*, based on molecular phylogenetic analysis (Fig. 1C) (Liu et al., 2020). Subsequently, a strain lacking a cell wall and morphologically resembling the cell-wall-less *Cz. merolae* was also assigned to the *Cyanidiococcus* clade based on molecular phylogenetic analyses (Fig. 1C) (Cho et al., 2020). Thus, molecular phylogenetic classification and the original classification based on cellular morphological characteristics (i.e., presence or absence of a cell wall and proliferation by binary fission or by forming four daughter cells within a mother cell) have become inconsistent, leading to taxonomic confusion. Under these circumstances, while culturing *Cyanidiococcus yangmingshanensis* (*Cc. yangmingshanensis*) NIES-4659—isolated by us from a Japanese sulfuric hot spring—we occasionally observed ovoid-shaped, smaller cells appearing within the population of originally cell-walled, spherical forms. These cells were morphologically similar to *Cz. merolae* 10D (described later).

Based on this observation, the apparent discrepancy between cellular morphology and molecular phylogeny, and the recent discovery of a sexual life cycle in *Galdieria* (Hirooka et al., 2022), we hypothesized that previously unrecognized sexual reproduction may also occur in *Cyanidiococcus*, *Cyanidioschyzon*, and even *Cyanidium*. Specifically, we assumed that the cell-walled form, which produces four daughter cells within a mother cell, may represent the diploid phase, whereas the cell-wall-less form, which proliferates by binary fission, may correspond to the haploid phase in these three genera. Supporting this hypothesis, a genome survey of these genera revealed the presence of gamete fusion and meiosis-related genes encoded in their genomes (Fig. 2A).

**Figure 2.**
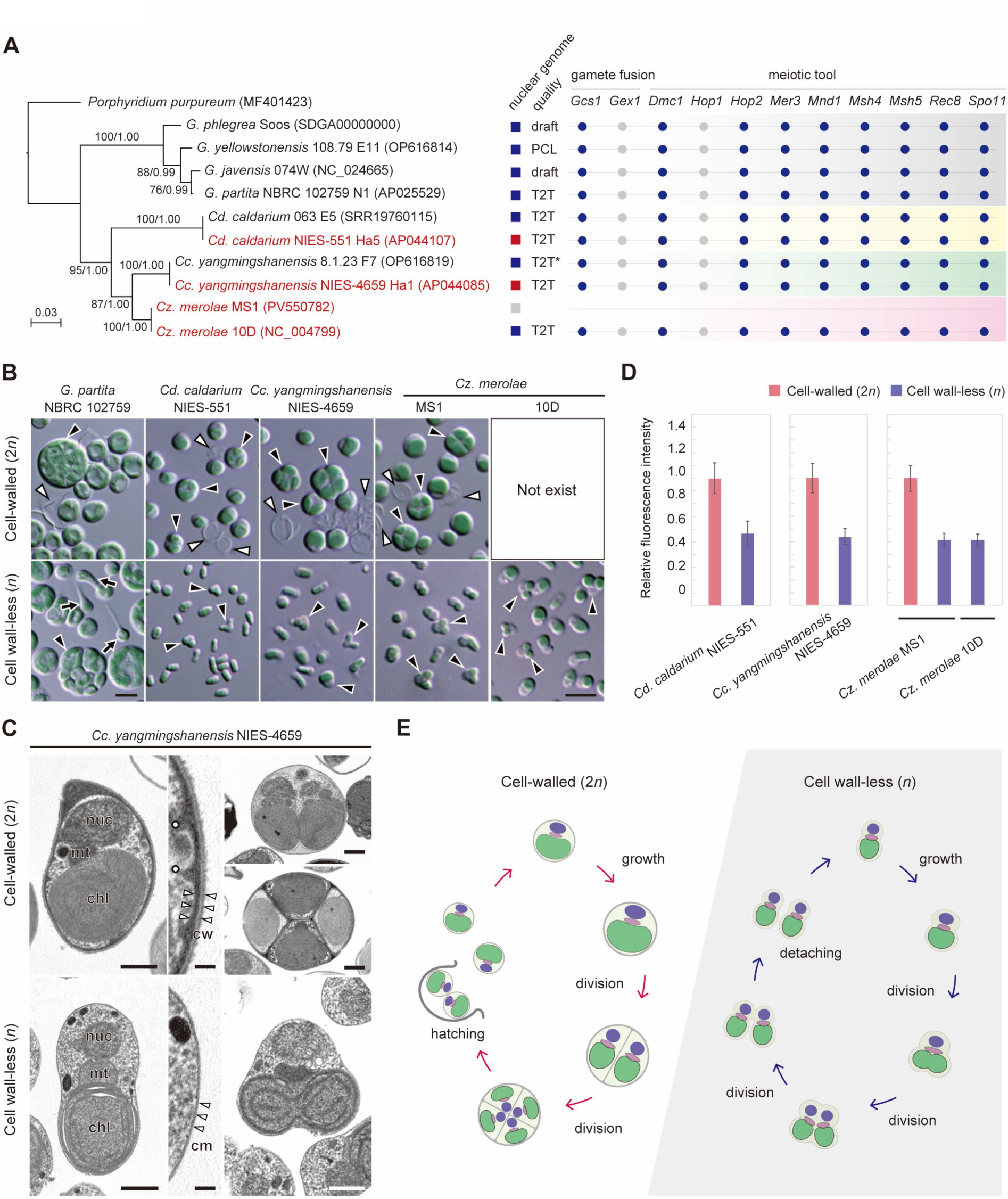
Sexual reproduction in Cyanidiophyceae. **A)** Phylogenetic relationships of several cyanidiophycean strains. The tree, based on amino acid sequences of the chloroplast-encoded *rbcL*, was generated by maximum likelihood analysis (RAxML-NG ver. 1.0.3)(Kozlov et al., 2019). *P. purpureum* was used as the outgroup. Bootstrap values >50% (left) obtained by ML analysis and posterior probabilities >0.95 (right) obtained by Bayesian analysis (MrBayes ver. 3.2.7)(Ronquist et al., 2012) are shown above the branches. Branch lengths represent evolutionary distances, as indicated by the scale bar. Strains used in this study are highlighted in red. The boxes on the right indicate strains for which genomes have been sequenced to date: blue, sequenced in previous studies; red, sequenced in this study; gray, not yet sequenced. The quality of assembled genomes was categorized into three classes: draft, PCL (pseudochromosome-level), and T2T (telomere-to-telomere). An asterisk (*) denotes a genome consisting of 19 T2T contigs and 1 single-end telomere contig (near-complete genome). Presence or absence of putative gamete fusion and meiotic tool genes in the genomes of cyanidiophycean strains was determined by BLASTp searches (gene IDs or GenBank accession numbers are listed in Supplementary Data Set 1). *Arabidopsis thaliana* genes were used as queries for gamete fusion–related genes (Mori et al., 2006; Alandete-Saez et al., 2011), and *Cz. merolae* 10D and *Galdieria javensis* 074W genes were used as queries for meiotic tool genes (Thangavel et al., 2023). Circles indicate presence (blue) or absence (gray). **B)** Differential interference contrast (DIC) micrographs of cell-walled cells (2*n*; original diploid clones) and cell wall-less cells (*n*; haploid clones generated in this study) of cyanidiophycean algae. Black arrowheads indicate dividing cells; the white arrowhead indicates the mother cell wall released upon hatching of daughter cells; the black arrow indicates tadpole-shaped cells of *G. partita*. Scale bar: 5 µm. **C)** Transmission electron micrographs of *Cc. yangmingshanensis* NIES-4659 2*n* and *n* cells. cm, cell membrane; *nuc*, nucleus; *chl*, chloroplast; *cw*, cell wall; *mt*, mitochondrion; white dots, eisosomes. Scale bars: 1 µm (left), 0.1 µm (center), 0.5 µm (right). **D**) Relative amounts of nuclear DNA in cell-walled and cell wall–less cells, based on fluorescence intensity of DAPI-stained nuclei. The mean fluorescence intensity of cell-walled cells was defined as 1.0. Data are means ± standard deviations (SDs) from 20 independent cells. **E**) Schematic illustration of proliferation modes in 2*n* and *n* cells of *Cc. yangmingshanensis*, *Cz. merolae*, and *Cd. caldarium*. The 2*n* cells possess a cell wall and reproduce asexually by forming four daughter cells within the mother cell wall, whereas *n* cells lack a cell wall and reproduce asexually by binary fission.

To test this hypothesis, we prepared three cell-walled cyanidiophycean strains *Cyanidium caldarium* (*Cd. caldarium*) NIES-551, *Cc. yangmingshanensis* NIES-4659, and *Cz. merolae* MS1 (Fig. 2B). Unlike *Cz. merolae* 10D, *Cz. merolae* MS1 is a cell-walled strain that was originally isolated from Yellowstone National Park, USA, and assigned to the *Cyanidioschyzon* clade based on phylogenetic analysis (Fig. 2A). Notably, *Cz. merolae* MS1 was a very rare contaminant in *Galdieria sulphuraria* (*G. sulphuraria*) CCMEE 5587.1 (originally described in Toplin et al., 2008; isolated from Yellowstone National Park). In photoautotrophic cultures of *G. sulphuraria* CCMEE 5587.1 aerated with 2% CO_2_, *Cz. merolae* MS1 grew faster than *G. sulphuraria*, became dominant in the culture, and was subsequently isolated as a clonal strain. The three cell-walled strains, maintained in MA medium (an inorganic medium) at pH 2.0, were spherical and proliferated by forming four daughter cells within a mother cell (Fig. 2, B and C; Supplementary Fig. S1, A and B). Based on previous findings that lowering the culture pH from 2.0 to 1.0 triggered a diploid-to-haploid transition in *Galdieria* (Hirooka et al., 2022), we transferred the cultures to lower-pH media—pH 0.75 for *Cd. caldarium* NIES-551 and pH 1.0 for the other strains—and incubated them for 2 to 3 weeks. As a result, ovoid, smaller cells resembling *Cz. merolae* appeared among the populations of originally cell-walled cells in all three cultures. When these *Cz. merolae*-like cells were isolated and cultured under the same low-pH conditions, they proliferated (Fig. 2B), and transmission electron microscopy confirmed that they lacked a cell wall, in contrast to the original spherical, cell-walled cells (Fig. 2C; Supplementary Fig. S1, A and B). When the nuclear DNA was stained with DAPI and quantified based on fluorescence intensity (Kuroiwa et al., 1986), the newly generated cell-wall-less forms in each strain showed, as expected, approximately half the DNA content of the original cell-walled forms (Fig. 2D). These results indicate that, as in *Galdieria*, the other genera of Cyanidiophyceae— *Cyanidium*, *Cyanidiococcus*, and *Cyanidioschyzon*—also possess a sexual life cycle, in which both the cell-walled diploid and the cell-wall-less haploid are capable of asexual reproduction (Fig. 2E). Thus, it became clear that *Cz. merolae* 10D, which has been used in research to date, should be recognized as being in the haploid phase of the *Cz. merolae* life cycle (further supporting evidence for this will be presented later).

The diploid cells of the three genera *Cyanidium*, *Cyanidiococcus*, and *Cyanidioschyzon* are morphologically indistinguishable by optical and transmission electron microscopy, and the same applies to their haploid forms (Fig. 2, B and C; Supplementary Fig. S1, A and B). As an exception, we found that the cell wall of *Cd. caldarium* NIES-551 diploid cells (∼50 nm thick) was thicker than those of *Cc. yangmingshanensis* NIES-4659 and *Cz. merolae* MS1 diploid cells (∼20 nm thick) (Fig. 2, C and E; Supplementary Fig. S1). This difference is likely a useful morphological marker to distinguish *Cyanidium* from *Cyanidiococcus* and *Cyanidioschyzon*. However, whether this feature represents a shared derived trait (synapomorphy) of the genus *Cyanidium* requires further investigation of additional strains in the future. Thus, at present, the distinction among *Cyanidium*, *Cyanidiococcus*, and *Cyanidioschyzon* relies primarily on differences in DNA sequences, chromosome numbers, and genome sizes (Cho et al., 2023; Park et al., 2023) (Fig. 1C; Table 1).

**Table 1.**
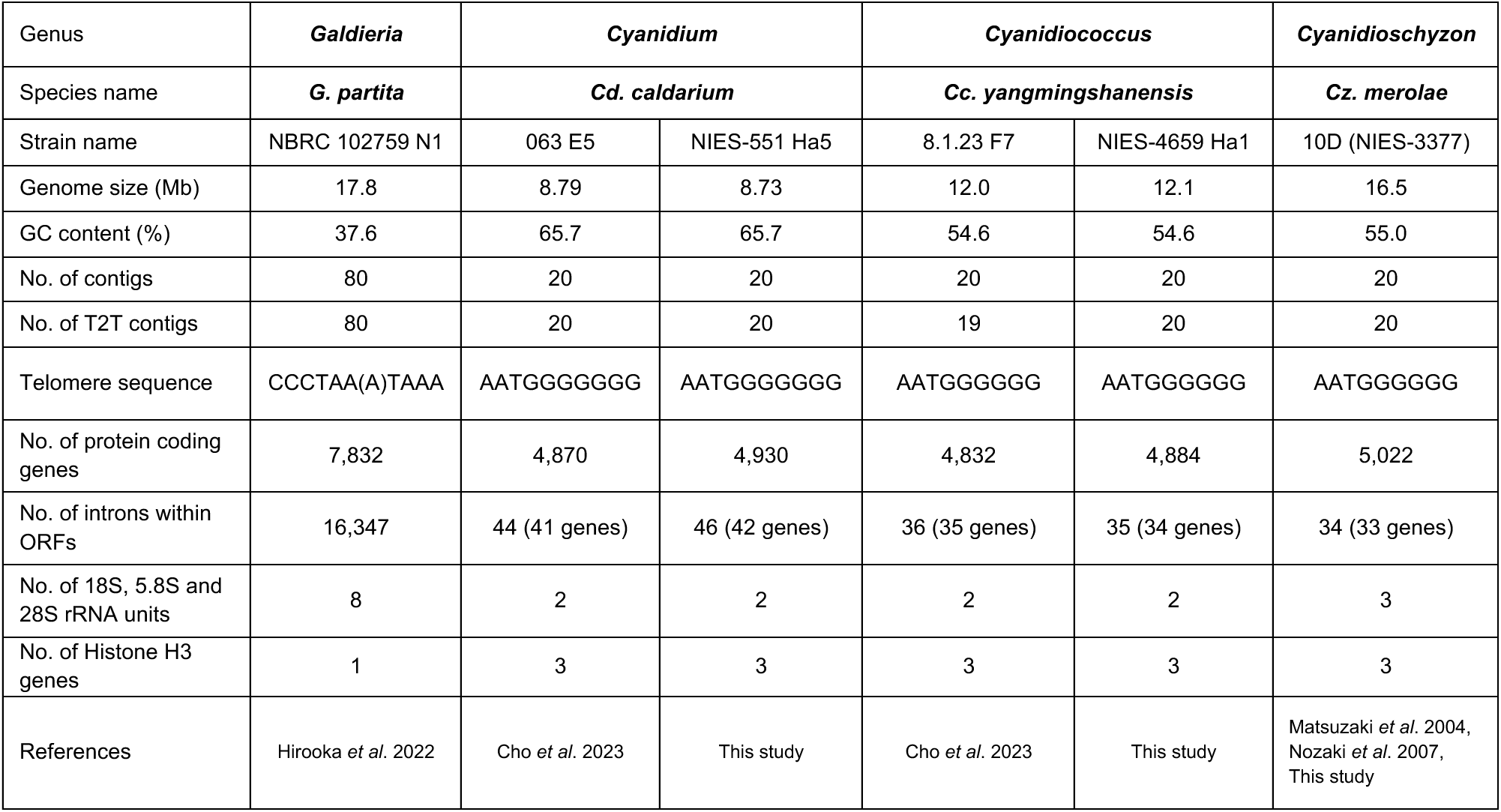
Comparison of genomic features among the Cyanidiophyceae genera *Galdieria*, *Cyanidium*, *Cyanidiococcus*, and *Cyanidioschyzon*.

In contrast, the haploid forms of *Cyanidium*, *Cyanidiococcus*, and *Cyanidioschyzon* are also clearly distinct from those of *Galdieria*, as with the known differences in the diploid forms described above: haploid cells of *Galdieria*, except during cell division, exhibit a tadpole-like shape with an extendable tail in which actin and microtubules are localized (Hirooka et al., 2022), whereas such structures are not observed in the haploid cells of the other three genera (Fig. 2B). Furthermore, haploid *Galdieria* cells form colonies of 4 to 32 daughter cells through successive cell divisions before the daughter cells detach from one another (Hirooka et al., 2022), while haploid cells of the other three genera proliferate by binary fission (Fig. 2, B and E).

### Development of genomic resources for Cyanidiophyceae through integration of haploid and diploid transcriptomes, intergeneric comparisons, and detection of meiotic recombination

To advance evolutionary and functional genomic analyses in Cyanidiophyceae, high-quality genome data with accurate gene annotation are essential. Although telomere-to-telomere (T2T) or near-complete genome assemblies have recently been established for representative strains of each genus except *Galdieria* (Fig. 2A) (Nozaki et al., 2007; Cho et al., 2023), gene prediction in these datasets was based solely on transcriptomic data from either the diploid or haploid phase. However, as shown below, certain genes are specifically expressed in either the haploid or diploid phase, and incorporating transcriptome data from both phases would enable more accurate genome annotation. For *Galdieria*, as described below, we recently generated a T2T genome assembly and integrated transcriptomic data from both haploid and diploid phases for *G. partita* (Hirooka et al., 2022). In this study, we newly constructed T2T genome assemblies with gene annotations for haploid clones of *Cd. caldarium* NIES-551 haploid clone 5 (Ha5) and *Cc. yangmingshanensis* NIES-4659 Ha1, and also reannotated the genome of *Cz. merolae* 10D.

The assembled nuclear genome sizes of *Cd. caldarium* NIES-551 Ha5 and *Cc. yangmingshanensis* NIES-4659 Ha1 are 8.73 Mb and 12.1 Mb, respectively (Table 1). These sizes are mostly consistent with those reported in previous studies on other strains of these species (*Cd. caldarium* 063 E5 and *Cc. yangmingshanensis* 8.1.23 F7) (Cho et al., 2023) (Table 1), as well as with the results of pulsed-field gel electrophoresis of chromosomes (Supplementary Fig. S2). The smaller genome sizes of these two genera compared to *Cz. merolae* 10D (16.5 Mb) are attributed to shorter length of intergenic regions rather than to differences in the number of genes as previously shown (Cho et al., 2023). We predicted gene models by incorporating RNA-seq data from both diploid and haploid phases, and identified 4,930 protein-coding genes in the nuclear genome of *Cd. caldarium* NIES-551 Ha5, of which 38 genes contained a single intron and 4 genes contained two introns within their ORFs, and 4,884 protein-coding genes in the nuclear genome of *Cc. yangmingshanensis* NIES-4659 Ha1, of which 34 genes contained a single intron and one gene contained two introns (Table 1). These gene numbers are greater than the previous predictions for other strains of these species, which identified 4,870 and 4,832 protein-coding genes, respectively (Table 1) (Cho et al., 2023).

In *Cz. merolae* 10D, previously published gene models were based on evidence from expressed sequence tags (ESTs) and amino acid sequence similarity to proteins from other organisms in the GenBank non-redundant (nr) database (Matsuzaki et al., 2004). This set of gene models, consisting of 4,803 protein-coding genes, has not been updated since 2007 (https://www.ncbi.nlm.nih.gov/datasets/genome/GCF_000091205.1/) and contains several annotation errors. Recently, 11 previously unannotated introns were identified in the *Cz. merolae* 10D genome through RNA-seq analysis (Wong et al., 2023), and the start codons of 161 genes were revised using ribosome profiling (Ribo-seq) (Mogi et al., 2025). Furthermore, in the present study, we identified an additional 219 protein-coding genes using RNA-seq data from *Cz. merolae* 10D (haploid) and from both the diploid and haploid phases of *Cz. merolae* MS1, as well as predicted protein data from *Cc. yangmingshanensis*. As a result, we obtained a re-annotated set of 5,022 protein-coding genes, of which 33 genes contained a single intron and one gene contained two introns (Table 1; Supplementary Data Set 3).

Regarding *Galdieria*, genome sequences of more than ten strains are currently available; however, none of them have been assembled at the chromosome level. Among them, recent advanced genomic studies have generated near-chromosome-level assemblies for diploid strains of *Galdieria* spp., including ACUF 138, ACUF 017, SAG 107.79, and 108.79 E11 (Downing et al., 2022; Cho et al., 2023). However, due to extensive segmental duplications across chromosomes, generating T2T genome assemblies from diploid strains remains challenging. To overcome this difficulty, we used a haploid clone to assemble a 17.8-Mb T2T genome of *G. partita* NBRC 102759 N1, which contains 7,832 protein-coding genes with an intron-rich gene structure (16,347 introns; Table 1) (Hirooka et al., 2022).

Then, we compared gene content across the four genera of Cyanidiophyceae. Because each genome contains groups of duplicated genes, such genes were counted as a single orthologous gene per species, and interspecies comparisons were performed using OrthoFinder (Emms and Kelly, 2019) (Fig. 3A). The comparative analysis revealed a total of 5,456 orthologous protein-coding genes in the Cyanidiophyceae pangenome, of which 3,206 orthologs (58.76%) were shared among all four genera. *Cd. caldarium*, *Cc. yangmingshanensis*, and *Cz. merolae* shared 4,339 orthologs (90.47%) out of the 4,796 orthologs identified in those three strains (Fig. 3A). Among the total 5,456 orthologous protein-coding genes in Cyanidiophyceae, 660 were uniquely present in *G. partita*, while 1,133 were shared exclusively by the other three strains (*Cd. caldarium*, *Cc. yangmingshanensis*, and *Cz. merolae*) and were absent in *G. partita* (Fig. 3A). These results indicate that the most pronounced divergence in genomic content exists between *Galdieria* and the other three genera—*Cyanidium*, *Cyanidiococcus*, and *Cyanidioschyzon*—which is consistent with their differences in cellular morphology (Fig. 2B) and phylogenetic relationships (Fig. 1C; Fig. 2A). Notably, although *Cd. caldarium* diverged from the common ancestor of *Cc. yangmingshanensis* and *Cz. merolae* approximately 0.3 billion years ago and their genome sizes and GC contents have diverged (Cho et al., 2023), most intron-containing genes are still shared among the three genera (Supplementary Data Set 4), suggesting that these introns play essential roles in their biological processes.

**Figure 3.**
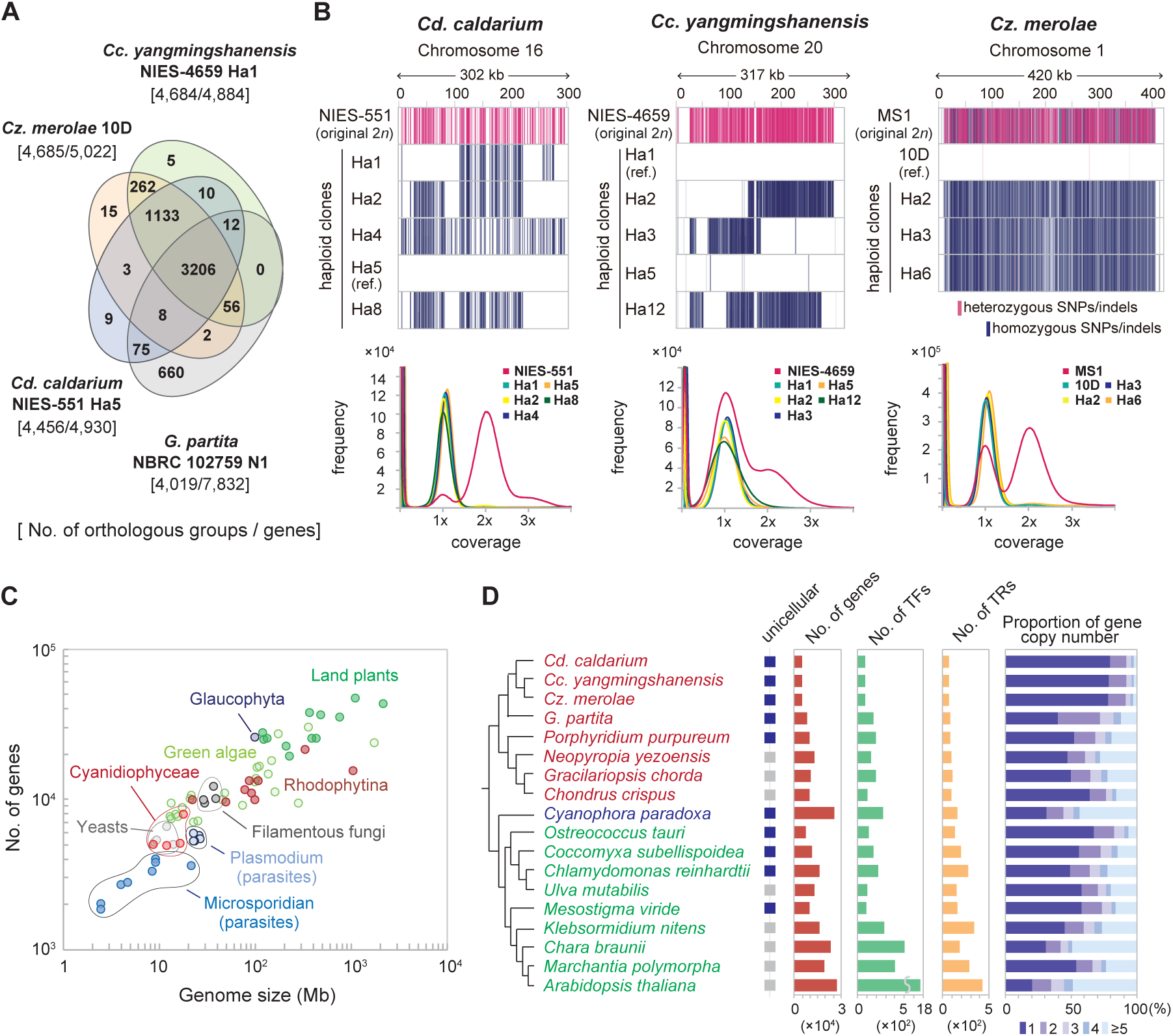
Comparison of genomic features of Cyanidiophyceae with other eukaryotes. **A)** Venn diagram showing the number of orthologs (identified by OrthoFinder; genes duplicated within respective genomes were counted as one) shared among the genomes of *G. partita* NBRC102759 N1, *Cd. caldarium* NIES-551 Ha5, *Cc. yangmingshanensis* NIES-4659 Ha1, and *Cz. merolae* 10D. **B)** Mapping of SNP/indel positions in the original 2*n* clones and their derived *n* clones of cyanidiophycean algae. Sites matching the haploid reference genome sequences are shown in white; heterozygous SNPs/indels in red; and homozygous SNPs/indels in blue. Histogram of *k*-mer counts (*k* = 41) was generated from whole-genome Illumina short reads using Jellyfish ver. 2.3.0 (Marcais and Kingsford, 2011). **C)** Comparison of the number of protein-coding genes and genome sizes of cyanidiophycean red algae with those of other eukaryotes. **D)** Comparison of the number of protein-coding genes, transcription factors (TFs), and transcriptional regulators (TRs) (TFs and TRs identified by iTAK (Zheng et al., 2016)), as well as the proportion of gene copy numbers in Archaeplastida. The dendrogram with species names on the left is based on previous studies (Munoz-Gomez et al., 2017; One Thousand Plant Transcriptomes, 2019). See also Supplementary Data Set 5 for details.

Using the genome assemblies generated above, we examined whether meiotic recombination occurs in *Cyanidium*, *Cyanidiococcus*, and *Cyanidioschyzon*, as previously observed in *Galdieria* (Hirooka et al., 2022), by analyzing single nucleotide polymorphisms (SNPs) and insertions/deletions (indels) in haploid clones derived from the original diploid clones. Numerous heterozygous SNPs and indels were detected when genomic reads from the original diploid clones (*Cd. caldarium* NIES-551, *Cc. yangmingshanensis* NIES-4659, and *Cz. merolae* MS1) were mapped to their respective reference genomes, indicating that each clone is a heterozygous diploid (Fig. 3B). Subsequently, when genomic reads from multiple distinct haploid clones derived from each original diploid clone were mapped to the corresponding reference genome, different SNP/indel patterns were observed depending on the haploid clone (Fig. 3B). Furthermore, *k*-mer analysis of genomic reads from the original diploid clones revealed two major peaks: one corresponding to heterozygous *k*-mers (with a peak at 1× coverage) and the other to homozygous *k*-mers (with a peak at 2× coverage) (Fig. 3B). In contrast, genomic reads from haploid clones exhibited a single peak (Fig. 3B). These results demonstrate, at the whole-genome level, that meiotic recombination occurs in these algae.

### Comparison of genome features with other eukaryotes

Based on the newly refined datasets, we compared the genomes of cyanidiophycean algae with previously sequenced genomes of other eukaryotes in terms of genome size and the number of protein-coding genes (Fig. 3C; Supplementary Data Set 5). The genome of *Cd. caldarium* (8.73 Mb) is among the smallest of all sequenced eukaryotic genomes, excluding parasitic microsporidia (2.5–21.6 Mb) (Fig. 3C), and is comparable to that of the methylotrophic yeast *Ogataea polymorpha* (8.97 Mb) (Riley et al., 2016). Specifically, the number of protein-coding genes (4,884–7,832) in cyanidiophycean algae is also comparable to those of yeasts such as *O. polymorpha* (5,177) (Riley et al., 2016), *Saccharomyces cerevisiae* (6,569) (Goffeau et al., 1996), and *Schizosaccharomyces pombe* (4,970) (Wood et al., 2002), although the nuclear genomes of cyanidiophycean algae additionally encode genes related to photosynthesis, as well as chloroplast biogenesis and regulation. Within Archaeplastida, several green algal lineages also possess streamlined nuclear genomes (12.9– 17.4 Mb); however, the number of protein-coding genes is relatively large, ranging from ∼7,300 to over 9,300 (Foflonker et al., 2018; Lemieux et al., 2019; Kato et al., 2023) (Supplementary Data Set 5). Comparative analyses of orthologous protein-coding genes within Archaeplastida further revealed that cyanidiophycean algae possess a remarkably streamlined gene repertoire compared with other photosynthetic eukaryotes (Supplementary Fig. S3). In addition, their nuclear genomes encode a relatively small number of regulatory genes, such as transcription factors (80–169) and transcriptional regulators (70–86) (Fig. 3D; Supplementary Data Set 5).

Gene duplication is widely observed among eukaryotic genomes and is considered one of the major evolutionary mechanisms, providing the genetic material necessary for the emergence of genes with new or modified functions (Ohno, 1970). In flowering plants (angiosperms), approximately 65% of genes have duplicated copies, most of which are derived from multiple ancient whole-genome duplication events (Panchy et al., 2016). In contrast, the genomes of cyanidiophycean algae (excluding *Galdieria*) have a high proportion of single-copy genes (∼80%; Fig. 3D). Taken together, Cyanidiophyceae possess one of the simplest genomic architectures among free-living eukaryotes examined to date, including photosynthetic lineages.

### Exclusive diploid dominance of Cyanidiophyceae in natural habitats

Most strains of *Galdieria*, *Cyanidiococcus*, and *Cyanidioschyzon* isolated from acidic hot springs worldwide possess a cell wall (Toplin et al., 2008; Huang et al., 2024). This observation suggests that Cyanidiophyceae predominantly exist in the diploid phase in natural environments. To investigate whether haploid cells are also present in these habitats, we conducted field sampling at two sulfuric hot springs in Japan: Kusatsu Hot Spring (Fig. 4A; Supplementary Fig. S4A–D) and Tsukahara Hot Spring (Supplementary Fig. S4E).

**Figure 4.**
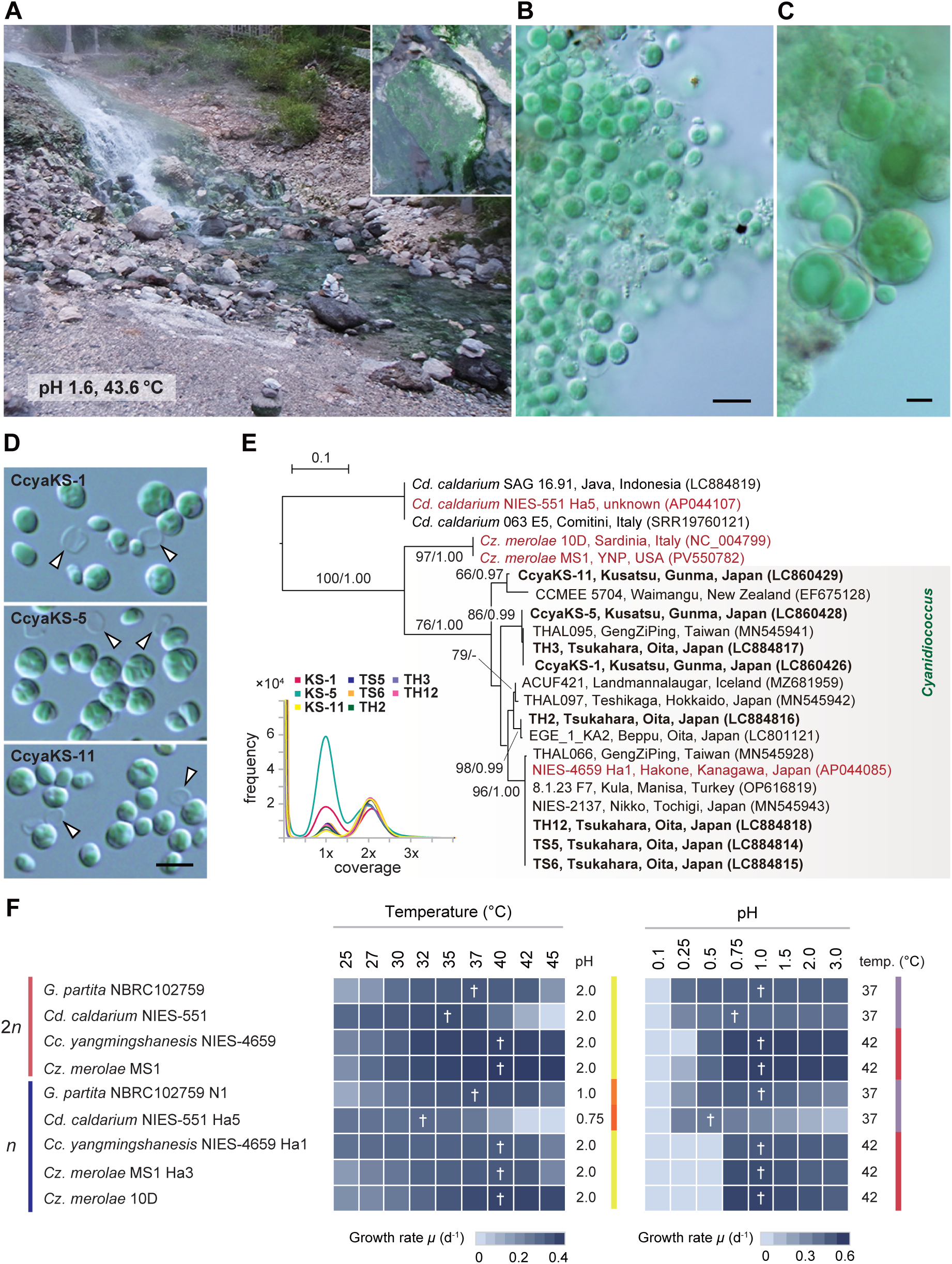
Natural habitat and optimal growth conditions of Cyanidiophyceae. **A)** Cyanidiophycean algae inhabiting an acidic hot spring in Kusatsu, Gunma Prefecture, Japan, were predominantly found in association with stones in the water. **B and C)** DIC micrographs of morphologically diploid *Cyanidiococcus* **(B)** and *Galdieria* **(C)** obtained from blue-green mats collected in the acidic hot spring. Scale bar: 5 µm. **D)** DIC micrograph of cultured cell-walled *Cyanidiococcus* strains isolated from the acidic hot spring in Kusatsu. The white arrowhead indicates the mother cell wall released upon hatching of daughter cells. Scale bar: 5 µm. **E)** Phylogenetic relationships of several cyanidiophycean strains, including strains collected from Kusatsu and Tsukahara (Oita Prefecture, Japan) hot springs (highlighted in bold). The tree, based on nucleotide sequences of the chloroplast-encoded *rbcL*, was generated by maximum likelihood analysis (RAxML-NG ver. 1.0.3) (Kozlov et al., 2019). *Cd. caldarium* was used as the outgroup. Bootstrap probabilities (BP) >50% (left) from ML analysis and Bayesian inference (BI) >0.95 (right) from Bayesian analysis (MrBayes ver. 3.2.7) (Ronquist et al., 2012) are shown above the branches. Branch lengths represent evolutionary distances, as indicated by the scale bar. Strains used in this study are highlighted in red. Histogram of *k*-mer counts (*k* = 41) generated from whole-genome Illumina short reads of *Cc. yangmingshanensis* strains collected from Kusatsu and Tsukahara hot springs, using Jellyfish ver. 2.3.0 (Marcais and Kingsford, 2011). **F)** Heatmaps showing the growth rate, based on increases in OD_750_, under the indicated temperature and pH conditions. Daggers (†) denote the highest growth rates for each alga. See also Supplementary Data Set 2 for details.

Cyanidiophycean algae were often found attached to stones on the riverbed, forming blue-green mats (Fig. 4A; Supplementary Fig. S4E). We collected these mats and examined them using microscopy, which revealed only cells with diploid morphology and no cells with haploid morphology (Fig. 4, B and C; Supplementary Fig. S4F). Among them, smaller cells morphologically similar to diploid *Cyanidium*, *Cyanidiococcus*, or *Cyanidioschyzon* were dominant (Fig. 4B; Supplementary Fig. S4F), although larger cells corresponding to *Galdieria* diploids were occasionally observed (Fig. 4C). To identify the taxonomic affiliation of the smaller cells, we isolated single cells and established clonal cultures. In all resulting cultures, remnants of the mother cell wall were observed following the hatching of daughter cells (Fig. 4D; Supplementary Fig. S4G). Phylogenetic analysis of the *rbcL* gene (encoded in the chloroplast genome) classified all clones into the genus *Cyanidiococcus* (Fig. 4E). Furthermore, *k*-mer analysis of Illumina short reads revealed two peaks (Fig. 4E), indicating that all clones are heterozygous diploids.

Members of the Cyanidiophyceae are also known to inhabit the surfaces of humid mud and the endolithic regions (a layer a few millimeters beneath the rock surface) above the waterline (Gross et al., 1998; Qiu et al., 2013; Van Etten et al., 2023), observations of these habitats likewise yielded only cell-walled diploid forms (Supplementary Fig. S4A–D).

Taken together, these results suggest that Cyanidiophyceae exist exclusively in the diploid phase in their natural habitats, and that haploid cells may arise only under specific and infrequent conditions, such as rare environmental stimuli or in locally distinct microhabitats.

### Comparison of growth rates among species and between haploid and diploid phases under various conditions

Several eukaryotic organisms have been identified in acidic environments (pH < 4.0), such as acid mine drainage (AMD) and geothermal hot springs (Amaral-Zettler, 2012). The low pH in these environments facilitates the solubilization of metals into water. Consequently, acidophilic organisms generally possess mechanisms to tolerate not only low pH but also high concentrations of metals, including toxic heavy metals.

To compare temperature preferences among the four genera of Cyanidiophyceae and between their respective diploid and haploid phases, cells of both phases of each strain were cultured across a temperature gradient ranging from 25°C to 45°C (Fig. 4F). Each culture was maintained at a specific pH: pH 1.0 for *Galdieria* haploid cells, pH 0.75 for *Cyanidium* haploid cells, and pH 2.0 for the remaining strains. In *G. partita*, diploid cells exhibited optimal growth at 30–42°C (Fig. 4F; here and below, the conditions for which the growth rate did not differ significantly from the peak according to Tukey’s multiple comparison test are defined as optimal), and haploids at 32–42°C, with both showing the highest growth rate at 37°C (Fig. 4F). *Cd. caldarium* showed optimal growth at 25–37°C for diploids (peak at 35°C) and haploids at 25–35°C (peak at 32°C) (Fig. 4F). *Cc. yangmingshanensis* diploid cells exhibited optimal growth at 32–45°C, and haploids at 27–42°C, with both peaking at 40°C (Fig. 4F). In *Cz. merolae,* diploid cells (strain MS1) exhibited optimal growth at 32–45°C, while haploid strains MS1 Ha3 and 10D exhibited optimal growth at 37–45°C and 32–42°C, respectively, with maximum growth again at 40°C for both phases (Fig. 4F).

To evaluate pH preferences, diploid and haploid cells of each strain were cultured across a pH gradient ranging from 0.1 to 3.0 (Fig. 4F). The temperature was set at 37°C for *Cyanidium* and at 42°C for the other strains. In *G. partita*, diploid cells exhibited optimal growth across pH 0.5–3.0, peaking at pH 1.0, whereas haploid cells preferred pH 0.75–1.0 with peak growth at pH 1.0 (Fig. 4F). In *Cd. caldarium*, diploid cells exhibited optimal growth at pH 0.5–3.0 and peaked at pH 0.75, while haploid cells grew best at pH 0.25–1.5 with a peak at pH 0.5 (Fig. 4F). In *Cc. yangmingshanensis*, both diploid and haploid cells exhibited optimal growth across pH 0.75–3.0 and showed maximum growth at pH 1.0. In *Cz. merolae*, diploid cells (strain MS1) exhibited optimal growth from pH 0.5 to 3.0, while haploid strains (MS1 Ha3 and 10D) exhibited optimal growth from pH 0.75 to 2.0, with peak growth at pH 1.0 for both phases (Fig. 4F).

These results indicate that in all four genera, diploid cells exhibit a broader range of tolerance to both temperature and pH compared to haploid cells. However, the optimal conditions for growth are largely similar between the haploid and diploid phases. Notably, both the diploid and haploid phases of *Cd. caldarium* NIES-551 exhibited a lower preferred temperature range compared to the other genera. Whether this trait represents a synapomorphy of the genus *Cyanidium* remains to be determined, and further investigation of additional strains is required. However, since its sister genus, *Cyanidiofrigus pintoensis* (Cyanidiales in Fig. 1C), inhabits meso-acidophilic environments (Huang et al., 2024), it is plausible that *Cyanidium* has adapted to more moderate conditions during its evolutionary divergence from other thermo-acidophilic members of Cyanidiophyceae.

To compare metal tolerance, diploid and haploid cells of each strain were cultured in MA medium supplemented with different concentrations of arsenate (As(V)) and nickel (Ni) ranging from 0 to 50 mM (Supplementary Fig. S4D). Given that metal toxicity is highly dependent on pH (Peterson et al., 1984), all media used in this assay were adjusted to pH 0.75. As a result, both diploid and haploid cells of *G. partita* exhibited higher tolerance to As(V) than other cyanidiophycean algae (Supplementary Fig. S4D), consistent with a previous study of *Galdieria yellowstonensis* (*G*. *yellowstonensis*) 108.79 E11 (Cho et al., 2023). This result is likely due to the presence of the *arsB* and *arsC* genes of HGT origin, which are encoded only in the *Galdieria* genome (Cho et al., 2023). Regarding Ni tolerance, both diploid and haploid cells of *Cd. caldarium* exhibited greater Ni tolerance than other cyanidiophycean algae (Supplementary Fig. S4H). A recent study showed that *Cz. merolae* 10D avoids Ni accumulation inside the cells in the presence of a high concentration of Ni (Marchetto et al., 2024). Although the molecular mechanism of Ni tolerance is still unclear, it is suggested that each member of Cyanidiophyceae has evolved its genome to adapt to different environments. For tolerance to both As(V) and Ni, there was little difference between the haploid and diploid phases (Supplementary Fig. S4H), unlike the larger ranges of temperature and pH tolerated by diploids compared with haploids (Fig. 4F), suggesting that the presence or absence of a cell wall does not affect heavy metal tolerance.

### Development of procedures for genetic manipulation

Among Cyanidiophyceae, procedures for genetic manipulation were first developed in *Cz. merolae* 10D, where linear DNA constructs are introduced into cell-wall-less cells via polyethylene glycol (PEG)-mediated transformation (Ohnuma et al., 2008), and arbitrary chromosomal loci can be precisely modified through highly efficient homologous recombination (Imamura et al., 2009; Fujiwara et al., 2013). Recently, we successfully adapted this method to *Galdieria* haploid cells, which also lacks a cell wall (Hirooka et al., 2022).

To further facilitate comparative and functional genomic studies in Cyanidiophyceae, we subsequently extended this genetic modification procedure to the cell-wall-less haploid cells of *Cc. yangmingshanensis* and *Cd. caldarium* (Fig. 5A). First, we constructed a plasmid encoding a *Venus* (a fluorescent protein) expression cassette and a chloramphenicol resistance gene (chloramphenicol acetyltransferase; *CAT*) as a selectable marker, flanked on both sides by sequences identical to a chromosomal intergenic region to allow targeted integration (Fig. 5B). Linear DNA, amplified by polymerase chain reaction (PCR), was introduced into pre-cultured cells of *Cc. yangmingshanensis* NIES-4659 Ha1 using a PEG-mediated transformation method developed for *Cz. merolae* 10D (Ohnuma et al., 2008; Fujiwara and Ohnuma, 2018), with minor modifications. The cells were then recovered for three days in iron-rich MA (IMA) liquid medium (an inorganic medium) at pH 1.2 under continuous light without any antibiotic, and subsequently transferred to the same medium supplemented with chloramphenicol for selection. After incubation for three weeks, chloramphenicol-resistant cells were observed (Fig. 5C). These cells were serially diluted into chloramphenicol-free IMA liquid medium or plating on cornstarch slurry spots over gellan gum–solidified IMA plates. After ∼2 weeks of incubation, clonal colonies appeared. PCR analysis of genomic DNA from a chloramphenicol-resistant clone of *Cc. yangmingshanensis* confirmed correct integration of the *Venus* expression cassette and *CAT* marker into the intergenic region (Fig. 5D). Stable expression of *Venus* was validated by immunoblotting and fluorescence microscopy (Fig. 5, E and F).

**Figure 5.**
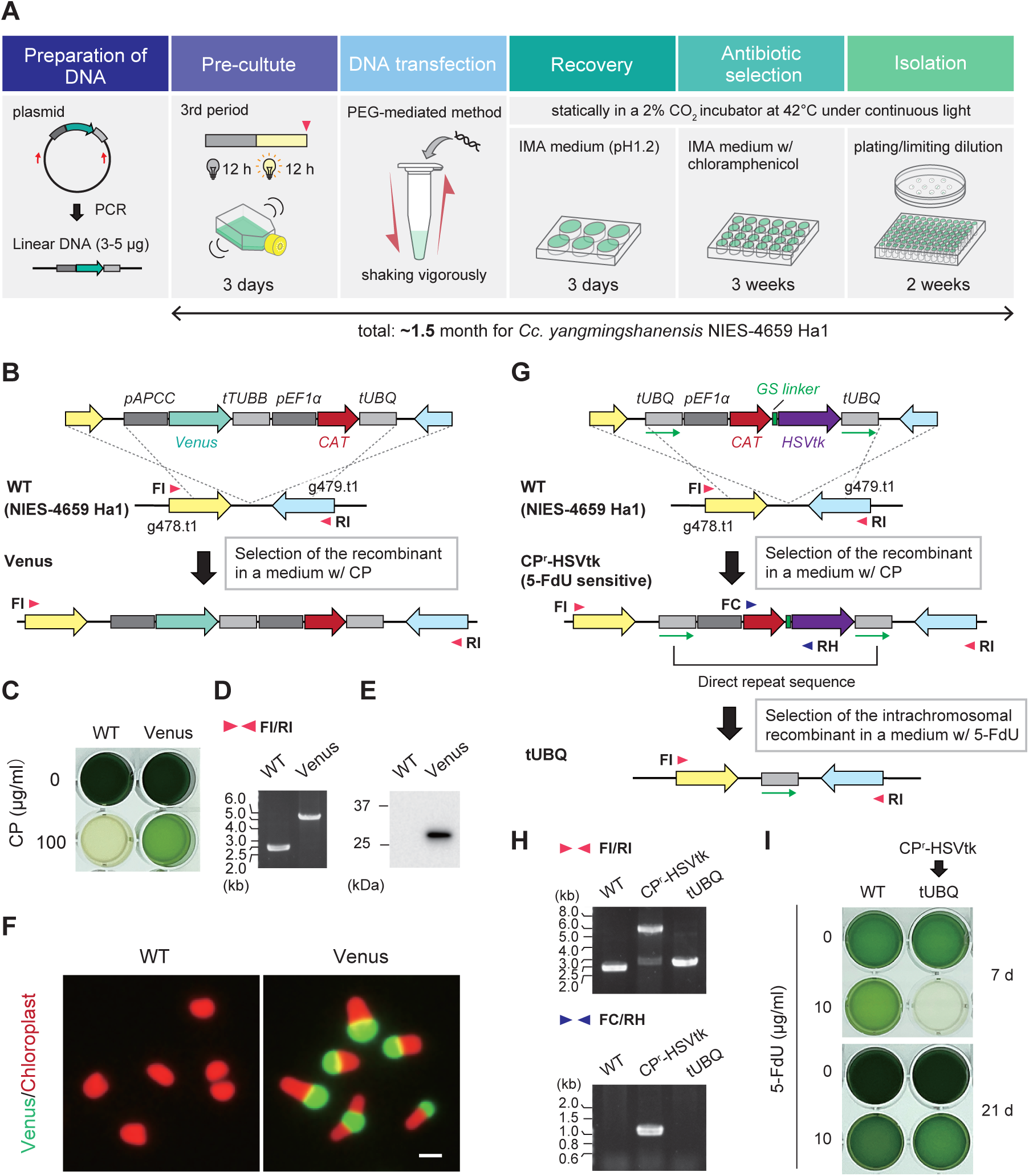
Genetic manipulation of *Cc. yangmingshanensis* NIES-4659 haploid clone Ha1. **A)** Workflow of the transformation protocol used in *Cc. yangmingshanensis* to obtain transformants. **B)** Schematic diagram showing insertion of the *Venus* expression cassette and the *CAT* selectable marker into a chromosomal intergenic region between the *g478.t1* and *g479.t1* loci by homologous recombination in the *n* clone. **C)** Transformed cells (Venus) were selected in IMA medium at pH 1.2 with chloramphenicol (CP) under continuous light for 3 weeks. **D)** Targeted integration of the transgenes into the chromosomal intergenic region was confirmed by PCR using primers FI and RI indicated by arrowheads in **B**. **E and F)** Expression of Venus protein was confirmed by immunoblotting with an anti–Green Fluorescent Protein (GFP) antibody **(E)** and by fluorescence microscopy **(F)** (green, Venus fluorescence; red, chloroplast fluorescence). The wild type (WT) served as a negative control. Scale bar: 2 µm. **G)** Schematic diagram showing targeted integration and subsequent removal of the *CAT* selectable marker. The *CAT* selectable marker and the *HSVtk* suicide marker, linked by a glycine–serine (GS) linker, were flanked by two directly repeated *tUBQ* sequences (green arrows). This construct was integrated into the intergenic region between the *g478.t1* and *g479.t1* loci of the wild-type haploid clone by homologous recombination. After selection of the transformant (CP^r^-HSVtk) in the presence of CP, the selectable marker was excised through intrachromosomal homologous recombination between the two *tUBQ* copies, followed by selection with 5-FdU, which is converted into a toxic product (5-FdUMP) by HSVtk. **H)** Confirmation of recombination events by PCR using the primers FI, RI, FC, and RH, indicated by arrowheads in **G**. **I)** The CP^r^-HSVtk cells were cultured for 21 days in the presence or absence of 5-FdU. The wild-type haploid clone served as a negative control. The procedure and results of genetic manipulation of *Cd. caldarium* NIES-551 haploid clone Ha5 are shown in Supplementary Fig. S5.

A transformant of *Cd. caldarium* NIES-551 Ha5 was obtained using the same procedure, except that the IMA medium was adjusted to pH 0.75 and the culture was maintained at 37°C. Additionally, because the *CAT* selectable marker was ineffective in this strain, it was replaced with a blasticidin S resistance cassette (blasticidin S deaminase; *BSD*) (Supplementary Fig. S5A), which had previously been shown to function in *Cz. merolae* 10D (Fujiwara et al., 2021) and *G. partita* NBRC 102759 haploid clones (Hirooka et al., 2022; Yamashita et al., 2025). As a result, a blasticidin S-resistant transformant expressing *Venus*, with integration of the *Venus* expression cassette and *BSD* marker into the intergenic region, was successfully obtained (Supplementary Fig. S5, B–D).

Recently, we developed a procedure in *G. partita* to remove the selectable marker (*BSD*) from a transformant using a combination of the herpes simplex virus thymidine kinase (*HSVtk*) suicide marker, which converts ganciclovir into a toxic compound (Smith et al., 1982), and intrachromosomal recombination (Hirooka et al., 2022). Using this technique and reusing the *BSD* selectable marker, two or more chromosomal loci can be modified in a single strain (Hirooka et al., 2022). To expand this technique to other members of Cyanidiophyceae, we first generated a *Cc. yangmingshanensis* transformant expressing a *CAT–HSVtk* fusion protein, in which HSVtk is fused to the C-terminus of CAT via a glycine-serine linker (Fig. 5, G and H). When the *CAT–HSVtk* transformant was cultured in MA medium supplemented with various concentrations of ganciclovir, its growth was slightly inhibited but not completely suppressed—contrary to expectations and unlike the case in *G. partita* (Supplementary Fig. S5E). Since HSVtk also converts 5-fluoro-2’-deoxyuridine (5-FdU) into the highly toxic compound 5-fluoro-2’-deoxyuridylate (5-FdUMP), which inhibits thymidylate synthase and leads to cell death (Supplementary Fig. S5F) (Yagil and Rosner, 1971), we tested whether 5-FdU could inhibit the growth of CAT–HSVtk-expressing *Cc. yangmingshanensis* cells. When both wild-type and *CAT–HSVtk* transformant cells were cultured under continuous light in media supplemented with different concentrations of 5-FdU (Supplementary Fig. S5E), most transformant cells died within 7 days, while wild-type cells survived at 10 µg/mL 5-FdU (Supplementary Fig. S5E). Upon continued cultivation, cells that had lost the *CAT–HSVtk* cassette via intrachromosomal homologous recombination between the direct repeat sequences (two copies of the *UBQ* terminator) flanking the *CAT– HSVtk* cassette (Supplementary Fig. S5F) appeared after 21 days (Fig. 5I). The expected recombination event was confirmed by PCR (Fig. 5H). Furthermore, we successfully applied this selectable marker deletion strategy to *Cz. merolae* 10D (Supplementary Fig. S5, G–J).

In summary, genetic manipulation at targeted chromosomal loci by PEG-mediated transformation of cell-wall-less haploids has now been developed for all axenically cultivable genera of Cyanidiophyceae—*Cyanidioschyzon*, *Cyanidiococcus*, *Cyanidium*, and *Galdieria*— and the method for selectable marker removal enables marker recycling for the editing of two or more loci.

### Diploid formation by isogamous mating of haploids and organelle genome inheritance

In *G. partita*, haploid cells not only undergo isogamous mating between distinct mating types to form heterozygous diploids, but also occasionally undergo self-diploidization by endoreduplication, resulting in homozygous diploids (Hirooka et al., 2022). Self-diploidization was also observed in *Cc yangmingshanensis* NIES-4659 Ha1, where cell-walled diploid cells occasionally emerged in the haploid clonal culture. By isolating and propagating the emerged diploid cell, a homozygous diploid culture derived from NIES-4659 Ha1 was obtained. However, this ability of self-diploidization disappeared after prolonged subculturing of haploids, unlike in *G. partita*. The resulting homozygous diploids, lacking interallelic polymorphisms, were suitable for direct comparison with haploids, as described below.

To investigate whether diploids arise through mating between haploids in cyanidiophycean genera other than *Galdieria*, uracil-auxotrophic, drug-resistant haploid strains were generated. This was achieved by replacing the chromosomal *URA5.3* (UMP synthase) locus with *Venus–CAT* in *Cc. yangmingshanensis* NIES-4659 Ha1 and *Cz. merolae* 10D, or with *Venus–BSD* in *Cd. caldarium* NIES-551 Ha5 (Fig. 6A, B; Supplementary Fig. S6A, B). These Δ*URA5.3* strains express *Venus* (Supplementary Fig. S6C) and exhibit uracil auxotrophy along with resistance to chloramphenicol or blasticidin S, whereas the wild-type strains are uracil prototrophic and drug-sensitive (Fig. 6C; Supplementary Fig. S6B).

**Figure 6.**
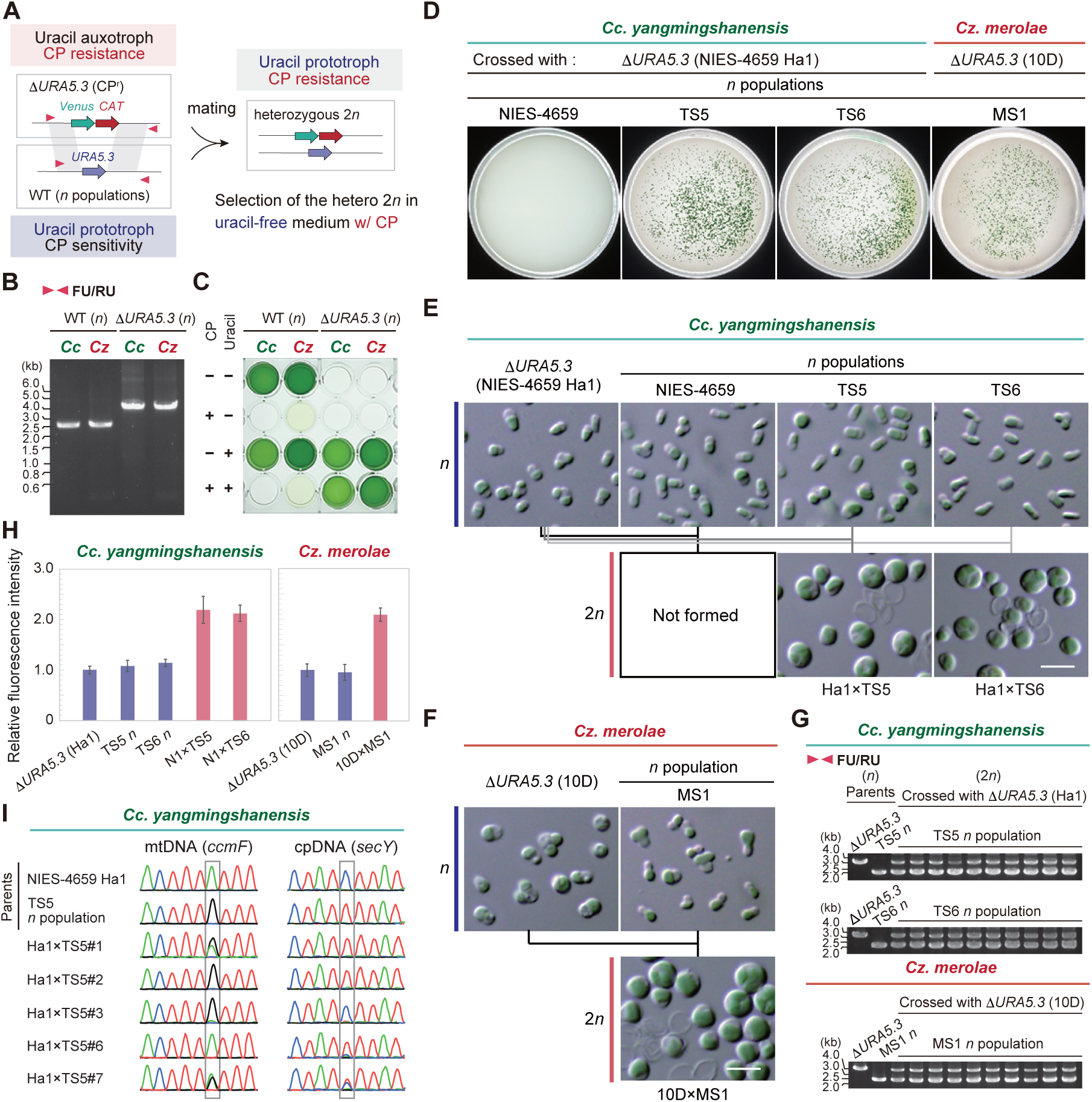
Diploid generation by haploid mating and organellar DNA inheritance in *Cc. yangmingshanensis* and *Cz. merolae*. **A)** Schematic illustration of the strategy for selecting heterozygous 2*n* cells possessing both the *CAT* selectable marker and the *URA5.3* gene, following mating of the Δ*URA5.3* (CP^r^) *n* clone with a wild-type *n* population derived from a 2*n* clone in uracil-free, CP-supplemented medium. **B)** Replacement of the chromosomal *URA5.3* locus with the *Venus* expression cassette and the *CAT* selectable marker in the generated Δ*URA5.3 n* clone was confirmed by PCR using primers FU and RU (arrowheads in **A**). The WT *n* clone served as a control. Cc, *Cc. yangmingshanensis*; Cz, *Cz. merolae.* **C)** Uracil auxotrophy and CP resistance of the Δ*URA5.3 n* clones were confirmed by cultivation for 7 days in the presence or absence of uracil and CP. WT *n* clones served as a control. Cc, *Cc. yangmingshanensis*; Cz, *Cz. merolae.* **D)** Δ*URA5.3* (CP^r^) *n* clones of *Cc. yangmingshanensis* and *Cz. merolae* were crossed with their respective wild-type *n* populations. Heterozygous 2*n* cells generated by mating were selected on HATF Immobilon nitrocellulose membranes (85 mm) placed over gellan gum– solidified medium at pH 2.0 with CP. The phylogenetic relationships among strains NIES-4659, TS5, TS6, 10D, and MS1 are shown in Fig. 4E. **E)** DIC micrographs of *Cc. yangmingshanensis*: Δ*URA5.3* (NIES-4659 Ha1) *n* clone, *n* populations derived from the original 2*n* clones NIES-4659, TS5, and TS6, and hybrid 2*n* clones Ha1×TS5 and Ha1×TS6. Scale bar: 5 µm. **F)** DIC micrographs of *Cz. merolae*: Δ*URA5.3* (10D) *n* clone, *n* population derived from the original 2*n* clone MS1, and the hybrid 2*n* clone 10D×MS1. Scale bar: 5 µm. **G)** Heterozygosity of the hybrid 2*n* clones confirmed by PCR using primers FU and RU. **H)** Nuclear DNA content compared by measuring fluorescence intensity of DAPI-stained nuclei. The mean fluorescence intensity of Δ*URA5.3 n* cells was defined as 1.0. Data are means ± standard deviations (SD) from 20 independent cells. **I)** Sanger sequencing chromatograms of mitochondrial (*ccmF*) and chloroplast (*secY*) DNA regions from parental *n* clones and their 2*n* progenies (colonies #1, #2, #3, #6, and #7). Gray boxes indicate nucleotide variants between the parental *n* clones. Green = adenine; red = thymine; black = guanine; blue = cytosine.

Therefore, diploids resulting from mating between Δ*URA5.3* haploids and wild-type haploids would possess both the drug-resistance gene (*CAT* or *BSD*) and *URA5.3*, and are thus expected to exhibit both drug resistance and uracil prototrophy (Fig. 6A). A similar strategy was previously employed in *G. partita* mating experiments (Hirooka et al., 2022).

For mating, each Δ*URA5.3* haploid clone was co-cultured with the corresponding wild-type haploid population—derived from heterozygous diploid strains—that should contain both mating types, if such sexuality exists, in MA medium supplemented with uracil, under a 12-h light/12-h dark cycle for 7 days (or 10 days for *Cd. caldarium*). After mating, the mixed cultures were transferred to selective conditions lacking uracil and containing the relevant drug. For *Cc. yangmingshanensis* and *Cz. merolae*, the cultures were plated on nitrocellulose membranes over gellan gum-solidified MA medium with chloramphenicol. For *Cd. caldarium*, the culture was incubated in liquid MA medium with blasticidin S.

As a result, no colonies appeared when Δ*URA5.3* (*Cc. yangmingshanensis* NIES-4659 Ha1) was crossed with the wild-type haploid population derived from the heterozygous diploid NIES-4659 (the reason remains unclear). However, crosses with other wild-type haploid populations (derived from heterozygous diploid TS5 or TS6; Fig. 4E; Supplementary Fig. 4C) did yield colonies (Fig. 6D). Similarly, for *Cz. merolae*, crosses between Δ*URA5.3* (10D) and wild-type haploid population derived from the MS1 diploid also produced colonies (Fig. 6D). For *Cd. caldarium*, crosses between Δ*URA5.3* (NIES-551 Ha5) and wild-type haploid populations derived from the diploids NIES-551 and SAG 16.91 yielded growing cells in the liquid medium as well (Supplementary Fig. S6D). The cells in these colonies or in the liquid culture showed diploid morphology (Fig. 6E and F; Supplementary Fig. S6E), genetic heterozygosity (Fig. 6G; Supplementary Fig. S6F), approximately twice the nuclear DNA content of haploid cells (Fig. 6H; Supplementary Fig. S6G), and expression of *Venus* (Supplementary Fig. S6H). These results indicate that *Cyanidium*, *Cyanidiococcus*, and *Cyanidioschyzon* also undergo isogamous sexual reproduction, and that *Cz. merolae* 10D is a haploid clone capable of sexual reproduction.

Notably, when Δ*URA5.3* haploid clones of *Cd. caldarium* and *Cz. merolae* were co-cultivated with their corresponding wild-type haploid clones (*Cd. caldarium* NIES-551 Ha5 and *Cz. merolae* 10D), no drug-resistant and uracil-prototrophic colonies or cells were observed (Supplementary Figs. S6D and S7A), suggesting that homothallic mating—i.e., mating type switching in a subset of cells within a population of the same mating type (Vanwinkle-Swift and Hahn, 1986)—does not occur in these species. The same was observed for *G. partita* (Hirooka et al., 2022). This experiment could not be conducted for *Cc. yangmingshanensis* because the Δ*URA5.3* clone of NIES-4659 Ha1 failed to generate diploids when crossed with the wild-type haploid population derived from the NIES-4659 diploid as above. In addition, when *Cz. merolae* 10D Δ*URA5.3* was co-cultivated with various haploid clones of *Cz. merolae* MS1 (Ha2, Ha3, Ha5, Ha6, Ha15), diploid colonies were produced in crosses with Ha2, Ha3, and Ha15 (i.e., drug-resistant and uracil-prototrophic), but not with Ha5 or Ha6 (Supplementary Fig. S7, A–C). These results suggest the presence of two mating types: Ha2, Ha3, and Ha15 versus 10D, Ha5 and Ha6.

Unlike nuclear DNA, organellar DNA—mitochondrial (mtDNA) and chloroplast (cpDNA)—exists in multiple copies within a single organelle and cell, and is uniparentally inherited in the majority of eukaryotes, although biparental inheritance has also been reported in several lineages (Birky, 1995). In angiosperms, approximately 20% of species exhibit biparental inheritance of cpDNA (Corriveau and Coleman, 1988; Zhang et al., 2003). While uniparental inheritance of cpDNA has been observed in the multicellular red alga *Pyropia yezoensis* (formerly *Porphyra yezoensis*) (Choi et al., 2008), the patterns of organellar DNA inheritance in unicellular red algae remain unclear. In *Cz. merolae* 10D, it has been reported that a single cell—containing a single mitochondrion and a single chloroplast—harbors approximately 80 copies each of mtDNA and cpDNA (Kuroiwa et al., 2021).

To clarify how organellar DNA is inherited in cyanidiophycean red algae, we performed crosses using haploid strain combinations that exhibit polymorphisms in both mtDNA and cpDNA: *Cc. yangmingshanensis* NIES-4659 Ha1 (Δ*URA5.3*) and TS5, *Cz. merolae* 10D (Δ*URA5.3*) and MS1, and *G. partita* NBRC 102759 N1 (Δ*URA1*; uracil auxotrophic and blasticidin S-resistant) (Hirooka et al., 2022) and *G. partita* NJ-5 (Supplementary Fig. S7D). We then analyzed the sequences of mtDNA and cpDNA regions containing nucleotide variations between the parental strains in multiple diploid progeny colonies derived from these crosses (Fig. 6I; Supplementary Fig. S8). For *Cd. caldarium*, we were unable to prepare two strains with polymorphisms in both mtDNA and cpDNA.

As a result, except for mtDNA in *G. partita*, the majority of diploid colonies exhibited mtDNA and cpDNA sequences derived from only one of the parental strains, selected in an apparently random manner. In some cases, both mtDNA and cpDNA originated from the same parent, while in others, they were derived from different parents, indicating that mtDNA and cpDNA are inherited independently. In contrast, some diploid colonies retained mtDNA (e.g., colonies #1, #4, #7, #8, #9, and #10 of *Cc. yangmingshanensis* Ha1 × TS5; colony #9 of *Cz. merolae* 10D × MS1) and cpDNA (e.g., colonies #7 and #8 of *Cc. yangmingshanensis* Ha1 × TS5; colony #9 of *Cz. merolae* 10D × MS1; colony #2, #4, #5, #7, #8, #9, and #10 of *G. partita* NBRC 102759 N1 × NJ-5) from both parents (Fig. 6I; Supplementary Fig. S8). In the yeast *S. cerevisiae*, it is known that mtDNA is biparentally inherited, but one parental type is rapidly lost, resulting in homoplasmy (Westermann, 2014). Based on this, we investigated whether only one parental organellar genome is eventually retained in each cell of diploid colony #7 of *Cc. yangmingshanensis* Ha1 × TS5, colony #9 of *Cz. merolae* 10D × MS1, and colony #2 of *G. partita* NBRC 102759 N1 × NJ-5, all of which initially inherited mtDNA and/or cpDNA from both parents (Supplementary Fig. S8). To test this, the diploid progenies were subcultured for 1 to 3 months, after which the cells were spread onto gellan gum-solidified medium. Single colonies were isolated and their mtDNA and cpDNA sequences were analyzed. As a result, all clones retained either mtDNA or cpDNA from only one parent, and the parental origin varied randomly among the colonies (Supplementary Fig. S8). In addition, in the above investigations, we analyzed the behavior of two loci located at the most distant positions on the circular genome of each organelle: *ccmF* and *nad6* for mtDNA, and *secY* and *psbD* for cpDNA. In all cases, the paired loci showed identical inheritance patterns (Supplementary Fig. S8), indicating that no recombination occurred between organellar genomes from the two parents.

These results suggest that, except for mtDNA in *G. partita*, organellar DNA in Cyanidiophyceae is initially inherited biparentally when a diploid is formed by mating between haploids. However, during diploid proliferation, organellar genomes derived from one parent (randomly determined) are gradually lost, resulting in homoplasmy. In contrast to these results, mtDNA in *G. partita* diploids was consistently inherited from a specific parent (in our experiments, from *G. partita* NBRC 102759 N1) (Supplementary Fig. S8A). This suggests that in Cyanidiophyceae, only *Galdieria* possesses a mechanism—similar to that found in most other eukaryotes—that ensures the uniparental inheritance of mtDNA from a specific mating type.

### Differences in transcriptome and histone modification between the diploid and haploid phases

To gain insights into the genetic basis of the phenotypic differences between diploid and haploid cells in Cyanidiophyceae, we first compared the transcriptomes of haploid *Cc. yangmingshanensis* NIES-4659 Ha1 and its derived homozygous diploid cells, which were spontaneously generated by self-diploidization as described above (Fig. 7A; Supplementary Data Set 7). In addition, to complement the transcriptome data, we conducted a comparative proteome analysis (Supplementary Data Set 7). The results showed that among the 4,884 nucleus-encoded genes, 159 genes (3.3%) were predominantly expressed in the diploid (FDR < 0.01, log_2_CPM > 2, log_2_FC(2*n/n*) > 2), with 139 of these supported by the proteome analysis, while 91 genes (2.8%) were predominantly expressed in the haploid (FDR < 0.01, log_2_CPM > 2, log_2_FC(2*n/n*) < −2), with 74 supported by proteome analysis (Supplementary Data Set 7). We also prepared comparative transcriptome datasets between diploid and haploid phases for *Cz. merolae*, *Cd. caldarium*, and *G. partita* (Supplementary Fig. S9; Supplementary Data Sets 8-10). Among them, similar expression patterns to those observed in *Cc. yangmingshanensis* were found in *Cd. caldarium* and *Cz. merolae*, which share 93.1% and 99.4% of their genes, respectively, with *Cc. yangmingshanensis* (Fig. 3A), as described below.

**Figure 7.**
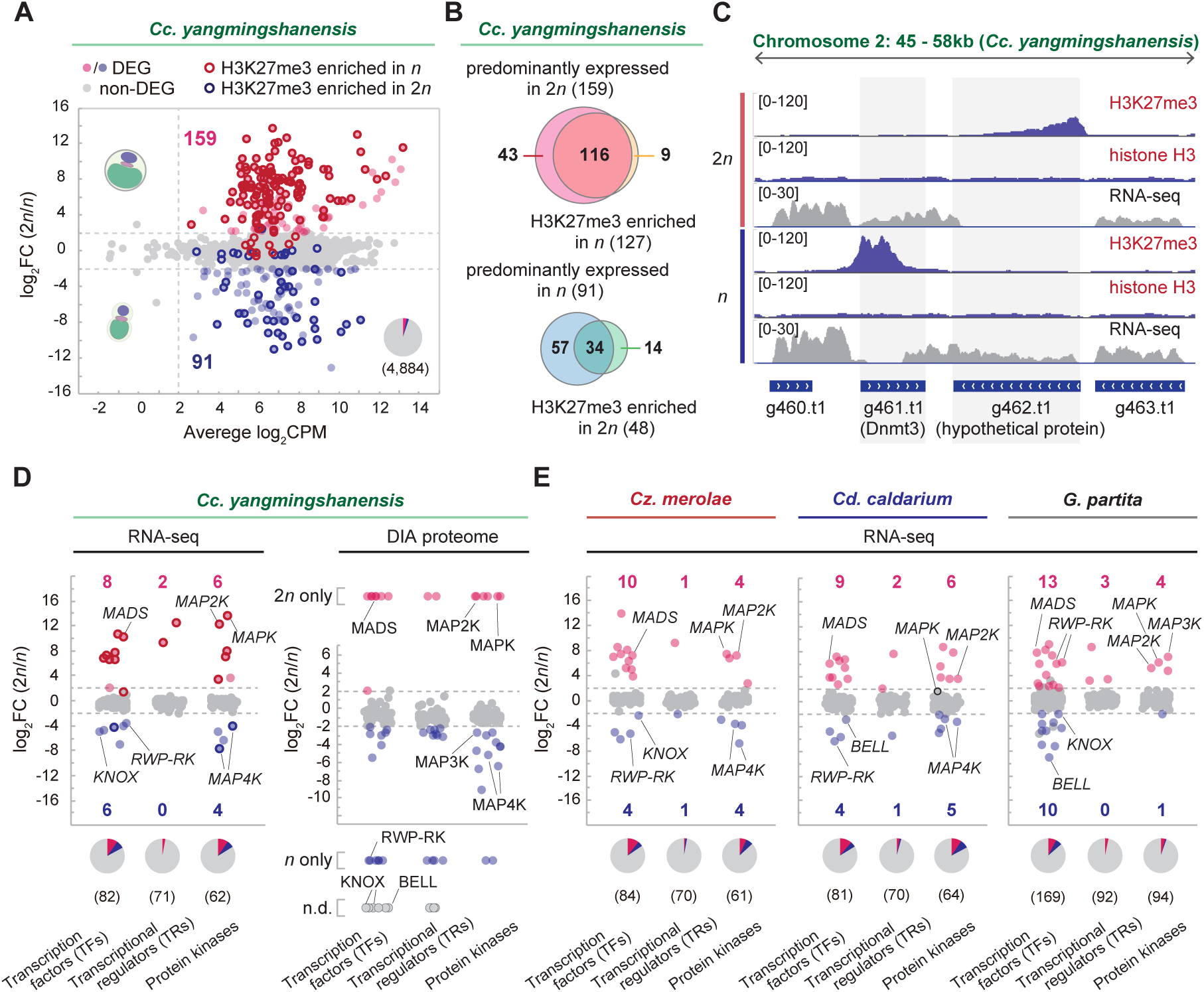
Comparison of transcriptomes and genomic distribution of H3K27me3 between diploid and haploid phases in Cyanidiophyceae. **A)** MA plot showing 159 genes upregulated in 2*n* (homozygous 2*n* derived from NIES-4659 Ha1) and 91 genes upregulated in *n* (NIES-4659 Ha1) among 4,884 nucleus-encoded genes (DEGs; FDR < 0.01, log CPM > 2, and |log FC| > 2 in three independent cultures). **B)** Venn diagrams showing the overlap between genes predominantly expressed in 2*n* cells and genes with elevated H3K27me3 levels in *n* cells (upper), and between genes predominantly expressed in *n* cells and genes with elevated H3K27me3 levels in 2*n* cells (lower). **C)** Patterns of read count levels from ChIP-seq of H3K27me3 and histone H3, and RNA-seq, in chromosome Ccyan0002: 45–58 kb in 2*n* and *n* cells. Normalized coverage (per million mapped reads) is shown. Numbers in brackets indicate value ranges. Gene models are shown as blue bars at the bottom. **D)** Ratios of mRNA and corresponding protein abundance between 2*n* and *n* cells for genes encoding transcription factors, transcriptional regulators, and protein kinases in *Cc. yangmingshanensis*. Numbers in parentheses below the pie chart indicate the number of genes in each category. See also Supplementary Data Set 7 for details. **E)** Ratios of mRNA between 2*n* and n cells for genes encoding transcription factors, transcriptional regulators, and protein kinases in *Cz. merolae, Cd. caldarium,* and *G. partita*. Numbers in parentheses below the pie chart indicate the number of genes in each category. See also Supplementary Data Sets 8-10 for details.

In multicellular organisms, the histone modification H3K27me3 silences the expression of specific sets of genes, leading to phenotypic outcomes such as cellular differentiation during development (Margueron and Reinberg, 2011). Recent studies have shown that H3K27me3 also occurs in some unicellular eukaryotes. In the diatom *Phaeodactylum tricornutum* (*P. tricornutum*), for example, genes involved in the establishment and maintenance of cell morphology are marked with H3K27me3, and changes in this modification are associated with morphological changes (Zhao et al., 2021). In *Cz. merolae* 10D (which turned out to be a haploid clone, as described above), Gene Ontology (GO) enrichment analysis revealed that H3K27me3-targeted genes are significantly enriched for terms related to development (Mikulski et al., 2017).

Based on these findings, to examine whether H3K27me3 is involved in phenotypic changes between the haploid and diploid phases in Cyanidiophyceae, we compared the genome-wide distribution of H3K27me3 between haploid *Cc. yangmingshanensis* NIES-4659 Ha1 and its derived homozygous diploid using chromatin immunoprecipitation sequencing (ChIP-seq) (Fig. 7A; Supplementary Data Set 7). The results showed that 127 genes are marked with H3K27me3 in haploid cells, 116 of which are predominantly expressed in diploid cells. In contrast, 48 genes are marked with H3K27me3 in diploid cells, 34 of which are predominantly expressed in haploid cells (Fig. 7B and C). Thus, in *Cc. yangmingshanensis*, the patterns of genes marked with H3K27me3 in haploid and diploid phases largely correlate with those whose expression is suppressed in the corresponding phases, suggesting that phenotypic changes during the life cycle are epigenetically regulated.

### Transcription factors and signaling pathways differentially expressed in haploid and diploid cells

The mitogen-activated protein kinase (MAPK) cascades regulate a wide variety of stimulus-responsive cellular processes, including proliferation, differentiation, and stress responses in eukaryotes (Plotnikov et al., 2011). The signals transmitted through these cascades enter the nucleus, where they modulate the activity of transcription factors (TFs) and chromatin remodeling proteins (Plotnikov et al., 2011). In each of the nuclear genomes of *Cc. yangmingshanensis*, *Cz. merolae*, and *Cd. caldarium*, three MAPKs, one MAPK kinase (MAP2K), one MAPK kinase kinase (MAP3K), and four MAPK kinase kinase kinases (MAP4Ks) were identified by iTAK (Supplementary Data Sets 7-9). Among them, one MAPK and one MAP2K were predominantly expressed in diploid cells, whereas two MAP4Ks were predominantly expressed in haploid cells in *Cc. yangmingshanensis*, *Cz. merolae*, and *Cd. caldarium* (Fig. 7D and E; Supplementary Fig. S9 A-C). The only minor exception was that, in *Cd. caldarium*, one *MAPK* gene showed slightly higher expression in haploid cells (log_2_FC(2*n*/*n*) = 1.76, below the threshold for differential expression defined in this study) (Fig. 7E). In *Cc. yangmingshanensis*, proteome analyses also showed similar expression patterns of MAPK cascade components (Fig. 7D). In *Galdieria*, three MAPKs, two MAP2Ks, one MAP3K, and five MAP4Ks were identified, of which one MAPK, one MAP2K, and one MAP3K were predominantly expressed in diploid cells (Fig. 7E). Thus, changes in MAPK cascades are likely involved in regulation of life cycle in Cyanidiophyceae.

Of the 82 transcription factors (TFs) encoded in the nuclear genome of *Cc. yangmingshanensis* (84 in *Cz. merolae*, 80 in *Cd. caldarium* and 169 in *G. partita*), eight genes—including one encoding a MEF2-type MADS-box TF—were predominantly expressed in diploid cells, while six genes—including those encoding KNOX and RWP-RK TFs—were predominantly expressed in haploid cells (Fig. 7D; Supplementary Data Set 7). A similar expression pattern of TF genes was also observed in *Cd. caldarium*, *Cz. merolae* and *G. partita* (Fig. 7E; Supplementary Data Sets 8-10). Among these TF genes, seven of the eight diploid-dominant TFs were marked with H3K27me3 in haploid cells, whereas only one of the six haploid-dominant TFs was marked with H3K27me3 in diploid cells in *Cc. yangmingshanensis* (Fig. 7D; Supplementary Data Set 7). Notably, the gene encoding DNA (cytosine-5)-methyltransferase 3 (Dnmt3)—the only predicted enzyme responsible for DNA methylation, which is associated with gene silencing in other eukaryotes (Lyko, 2018)—was predominantly expressed in diploid cells in *Cc. yangmingshanensis*, *Cz. merolae* and *Cd. caldarium* (Fig. 7C; Supplementary Data Sets 7-9). These observations raise the possibility that distinct gene-silencing mechanisms are involved in haploid-to-diploid and diploid-to-haploid transitions. In contrast, the *Galdieria* genome encodes three *Dnmt3* genes, which are expressed at similar levels in both diploid and haploid cells (Supplementary Data Set 10), suggesting that the gene-silencing mechanism associated with DNA methylation may differ from that in other cyanidiophycean algae.

The MADS-box family of eukaryotic transcription factors is involved in various cellular processes, including cell differentiation, cell wall reinforcement, and arginine metabolism in yeast and animals (Messenguy and Dubois, 2003). In angiosperms, the MIKC-type MADS-box family has undergone substantial expansion and regulates diverse developmental processes, including flower formation (Smaczniak et al., 2012). MIKC-type MADS-box genes are conserved in streptophyte algae and land plants (together classified as Streptophyta in Fig 1A) and are thought to have evolved from ancestral MEF2-type MADS-box genes—found in both green and red algae—through the acquisition of additional domains, such as the I and K domains (Thangavel and Nayar, 2018). However, little is known about the function of MEF2-type MADS-box genes in unicellular green and red algae, although a previous study suggested that in the unicellular green alga *Coccomyxa subellipsoidea*, the gene is involved in development and stress tolerance (Nayar and Thangavel, 2021). We previously found that in *Galdieria*, the single MEF2-type MADS-box gene encoded in the genome is predominantly expressed in the diploid phase, and its deletion inhibits the haploid-to-diploid transition, suggesting that it plays a role in diploid cell formation (Fig. 7E) (Hirooka et al., 2022). Similarly, in *Cc. yangmingshanensis*, *Cz. merolae* and *Cd. caldarium*, the single MEF2-type MADS-box gene encoded in each genome was also predominantly expressed in diploid cells (Fig. 7D and E). In *Cc. yangmingshanensi*s, proteome analyses also showed the exclusive expression of MADS-box protein in the diploid phase (Fig. 7D). These results suggest a conserved function of this gene across Cyanidiophyceae. The function of such MEF2-type MADS-box genes in diploid formation may reflect their origin in Archaeplastida, although future analyses, such as the identification of their target genes, could provide further insights.

KNOX and BELL-related proteins, both of which are TALE homeodomain (HD) transcription factors, regulate sexual life cycle transitions in some lineages of Viridiplantae (green algae and land plants) (Lee et al., 2008; Sakakibara et al., 2013; Horst et al., 2016; Dierschke et al., 2021; Hisanaga et al., 2021). Similarly, the TALE-HD TFs OUROBOROS and SAMSARA promote the transition from the gametophyte (haploid) to the sporophyte (diploid) generation after fertilization in the multicellular brown alga *Ectocarpus* (Coelho et al., 2011; Arun et al., 2019). In the unicellular green alga *Chlamydomonas reinhardtii*, the KNOX protein GSM1 and the BELL-related protein GSP1 and the are exclusively expressed in minus and plus mating-type gametes, respectively (Lee et al., 2008). These proteins heterodimerize, trigger nuclear and organellar fusion between mating types (Kariyawasam et al., 2019), and activate diploid gene expression after mating (Lee et al., 2008). Furthermore, in multicellular red algae, the expression level of the *KNOX* gene varies with life cycle transitions in *Pyropia yezoensis* (Mikami et al., 2019), and a distinct pair of TALE-HD TFs is encoded on the sex-determining regions (SDRs) of the female and male, respectively, in *Bostrychia moritziana* (Petroll et al., 2025a). In *Cc. yangmingshanensis*, *Cz. merolae, Cd. caldarium* and *G. partita*, two *KNOX* genes and a single *BELL*-related gene are encoded in their nuclear genomes (Joo et al., 2018) (Supplementary Data sets 7-10). We previously showed that, in *Galdieria*, one of the two *KNOX* genes and the single *BELL*-related gene were predominantly expressed in haploids of both mating types, and deletion of either transcription factor inhibited the haploid-to-diploid transition (Hirooka et al., 2022). In contrast, in *Cc. yangmingshanensis*, the one *KNOX* gene—but not the *BELL*-related gene—was predominantly expressed in haploid cells (Fig. 7D). However, proteome analysis failed to detect BELL-related proteins, and moreover, KNOX proteins were also not detected (Fig. 7D; Supplementary Data Set 7). We also found that, in *Cz. merolae*, the *KNOX* gene was predominantly expressed in haploid cells of both mating types (10D and MS1 Ha3), while the *BELL*-related gene was not (Fig. 7E; Supplementary Data Set 8). Conversely, in *Cd. caldarium*, the *BELL*-related—but not the *KNOX* gene—was predominantly expressed in haploid cells (Supplementary Data Set 9). Considering these results, the haploid cultures of these three species used in this study may represent populations of vegetative haploid cells.

Upon some form of stimulation, they may transform into as-yet-unknown gamete forms in which BELL-related and KNOX proteins are expressed as a result of a substantial increase in the mRNA levels of these genes, compared to those observed in this study.

RWP-RK transcription factors, found only in Archaeplastida, are involved in responses to nitrogen availability and sexual reproduction-associated processes (e.g., gamete sex determination, gametogenesis, and gametophyte development) in Viridiplantae (Chardin et al., 2014). Only a single RWP-RK TF gene is encoded in each genome of *Cc. yangmingshanensis*, *Cz. merolae* and *Cd. caldarium*, and it is predominantly expressed in haploid cells (Fig. 7D and E). In *Cc. yangmingshanensi*s, proteome analyses also showed the exclusive expression of RWP-RK protein in the haploid phase (Fig. 7D). In contrast, in *G. partita*, among the eleven RWP-RK TF genes encoded in the nuclear genome, two were predominantly expressed in diploid cells (Fig. 7E; Supplementary Data Set 10). In *P. yezoensis*, it was reported that RWP-RK TF genes were upregulated both in gametophytes treated with the ethylene precursor 1-aminocyclopropane-1-carboxylic acid (ACC)—which promotes gamete formation—and in mature gametophytes (Uji et al., 2016). Although the precise roles of RWP-RK TFs in red algae remain to be elucidated, these observations suggest their possible involvement in sexual reproduction-associated processes, highlighting their deep evolutionary origin within Archaeplastida.

### Differentially expressed genes underlying differences in cellular architecture between haploid and diploid cells

Cell walls in photosynthetic eukaryotes display a considerable degree of diversity in their compositions and molecular architectures. Red algae typically possess composite cell walls that are primarily composed of cellulose microfibrils and also contain unique polysaccharides called sulfated galactans, including the economically important carrageenan and agar (Borg et al., 2023). However, Cyanidiophyceae have lost genes encoding cellulose synthase (CesA) and cellulose synthase-like (Csl) enzymes (Yin et al., 2009), and are thus thought to possess a unique cell wall. Notably, their ability to survive several months of darkness suggests that the cell wall is likely impermeable to protons, possibly as an adaptation to highly acidic environments (Gross, 2000). In addition, previous study showed that in *Cd. caldarium* (the strain number was not reported, and thus its phylogenetic position remains unclear; it could potentially be assigned to *Cc. yangmingshanensis* or *Cz. merolae*), the cell wall is largely proteinaceous (50–55%), unlike other red algae (Bailey and Staehelin, 1968).

In algae and plants, genes encoding secretory proteins and glycosyltransferases contribute to the formation of the extracellular matrix and cell wall. In *Cc. yangmingshanensis*, *Cz. merolae, Cd. caldarium* and *G. partita*, approximately 35–50% of the genes encoding secretory proteins are differentially expressed between cell-walled diploid and cell wall-less haploid cells (Fig. 8A and B). We found that, in *Cc. yangmingshanensis*, *Cz. merolae* and *Cd. caldarium*, genes encoding secretory proteins such as fasciclin (FAS1) domain proteins and lysyl oxidase (LOX) were predominantly expressed in diploid cells (Fig. 8A and B). In *Cc. yangmingshanensi*s, proteome analyses also showed the predominant expression of these proteins in the diploid phase (Fig. 8A). FAS1 domain proteins are membrane-anchored glycoproteins involved in building cell wall structures in plants (Silva et al., 2020), and in *G. partita*, they are also expressed exclusively in diploid cells (Fig. 8B) (Hirooka et al., 2022). LOX plays a critical role in the formation and repair of the extracellular matrix by oxidizing lysine residues in fibrous proteins such as elastin and collagen in animals (Kagan and Li, 2003), and it has been identified as an extracellular protein in *Cc. yangmingshanensis* RK-1 (renamed from *Cd. caldarium* RK-1 by Park et al., 2023) (Tomo et al., 2018). LOX domain-containing genes are widely distributed in animals and other eukaryotes, as well as bacteria and archaea (Grau-Bove et al., 2015). However, within Archaeplastida, they have been identified only in *Porphyra* and in cyanidiophycean algae (except for *Galdieria*), suggesting the acquisition of these genes through HGT from other organisms.

**Figure 8.**
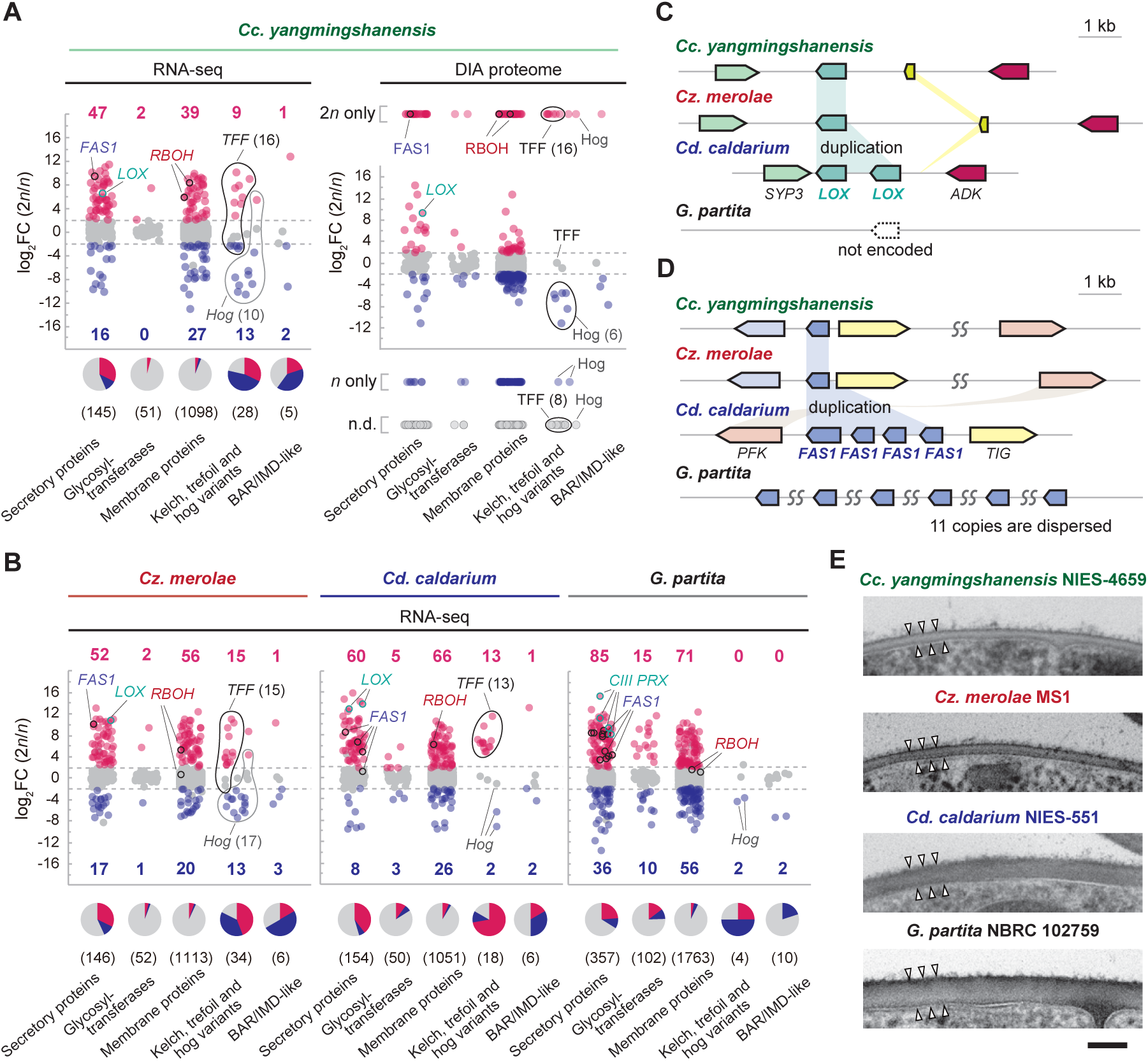
Comparison of transcriptomes and copy numbers of cell wall-related genes in Cyanidiophyceae. **A)** Ratios of mRNA and corresponding protein abundance between 2*n* and *n* cells for genes encoding secretory proteins, glycosyltransferases, and other functional categories related to cell wall formation/regulation in *Cc. yangmingshanensis*. Numbers in parentheses below the pie chart indicate the number of genes in each category. See also Supplementary Data Set 7 for details. **B)** Ratios of mRNA abundance between 2*n* and *n* cells for genes encoding secretory proteins, glycosyltransferases, and other functional categories related to cell wall formation/regulation in *Cz. merolae, Cd. caldarium,* and *G. partita*. Numbers in parentheses below the pie chart indicate the number of genes in each category. See also Supplementary Data Sets 8-10 for details. **C and D)** Variation in the copy numbers of the cell wall–related genes *LOX* **(C)** and *FAS1* **(D)** in the nuclear genomes of cyanidiophyceaen algae. **E)** Transmission electron micrographs of the cell walls of diploid cells in cyanidiophycean algae. Scale bar: 100 µm.

We also found that the nuclear genomes of *Cd. caldarium* NIES-551 Ha5 and the previously sequenced *Cd. caldarium* 063 E5 (Cho et al., 2023) encode two copies of *LOX* and four copies of *FAS1* genes, all of which are arranged in tandem repeats, in contrast to the genomes of *Cz. merolae* 10D (Matsuzaki et al., 2004) and *Cc. yangmingshanensis* 8.1.23 F7 (Cho et al., 2023) and NIES-4659 Ha1, which each possess a single copy of these genes (Fig. 8C and D). In *G. partita*, eleven copies of *FAS1* genes are dispersed throughout the genome (Fig. 8D). These differences, particularly in the copy number of *FAS1* genes, might be related to the variation in cell-wall thickness observed among *Cc. yangmingshanensis* NIES-4659, *Cz. merolae* MS1, *Cd. caldarium* NIES-551, and *G. partita* NBRC 102759 (Fig. 8E).

Regarding glycosyltransferase genes, in *Cc. yangmingshanensis*, only two were predominantly expressed in diploid cells (two in *Cz. merolae* and five in *Cd. caldarium*), whereas in *G. partita*, 13 were predominantly expressed in diploid cells (Fig. 8A and B). Thus, *Cyanidiococcus* (as well as *Cyanidioschyzon* and *Cyanidium*) likely have a simpler cell wall polysaccharide composition than *Galdieria*.

In plants, reactive oxygen species (ROS), such as superoxide (O₂⁻) and hydrogen peroxide (H₂O₂), play important regulatory roles in cell differentiation and development (Mhamdi and Van Breusegem, 2018). Some enzymes, such as respiratory burst oxidase homologs (RBOHs) and class III peroxidases (CIII PRXs), play predominant roles in apoplastic ROS regulation, contributing to cell wall loosening and hardening in plants (Cosio and Dunand, 2009). The genome of *Cc. yangmingshanensis* encodes two *RBOH* genes (two in *Cz. merolae* and one in *Cd. caldarium*), both of which are predominantly expressed in diploid cells of *Cc. yangmingshanensis* (Fig. 8A), a result supported by proteome analyses (Fig. 8A). In *Cz. merolae*, one of the two *RBOH* genes, and in *Cd. caldarium*, the single *RBOH* gene is predominantly expressed in diploid cells (Fig. 8B). In *P. yezoensis*, RBOH is suggested to be involved in archeospore (asexual reproductive spore) formation, which is associated with changes in cell wall composition (Gui et al., 2022). Although two *RBOH* genes are encoded in the *G. partita* genome, they are expressed at similar levels in both diploid and haploid cells (Supplementary Data Set 10). In contrast, the *G. partita* genome encodes four *CIII PRX* genes, all of which are predominantly expressed in diploid cells (Fig. 8B) (Hirooka et al., 2022). No *CIII PRX* genes are found in the genomes of *Cyanidiococcus*, *Cyanidioschyzon*, or *Cyanidium*, nor in other red algae. The presence of *RBOHs* and *CIII PRXs*—enzymes typically found in plants—in Cyanidiophyceae (despite lineage-specific losses), and their expression specifically in cell-walled diploid cells, highlight the deep evolutionary origin of apoplastic ROS regulation by these proteins within Archaeplastida.

The majority of the 18–34 genes encoding proteins with combinations of Hedgehog (Hh), trefoil factor (TFF), and kelch (K)-like domains (e.g., TFF, Hog, K+TFF, K+Hog) are encoded in the subtelomeric regions of several chromosomes in *Cd. caldarium*, *Cc. yangmingshanensis*, and *Cz. merolae* (Nozaki et al., 2007; Cho et al., 2023). In the *G. partita* genome, two K-like domain–containing genes and two Hh-like domain–containing genes are encoded, with the *Hh*-like genes located in subtelomeric regions, whereas *TFF*-like genes are absent. However, the proteins previously annotated as Hh in Cyanidiophyceae retain the carboxy-terminal autocatalytic domain HhC, which contains a Hint module, but lack the amino-terminal signaling domain HhN typically found in bona fide Hh proteins. Thus, these proteins encoded in cyanidiophycean genomes have been reclassified as non-Hh Hog proteins (Burglin, 2008). While bona fide Hh is restricted to metazoans, non-Hh Hog proteins—of unknown function but potentially involved in processes such as cell division, membrane organization, stress response, or extracellular matrix construction through lipid modification of proteins—are widespread across eukaryotes (Burglin, 2008). Of the 10 *Hog* genes (K+Hog and Hog) encoded in the *Cc. yangmingshanensis* genome (17 in *Cz. merolae*, three in *Cd. caldarium* and two in *G. partita*), one was predominantly expressed in diploid cells, while eight were predominantly expressed in haploid cells, a result supported by proteome analyses (Fig. 8A). A similar expression pattern of *Hog* genes was also observed in *Cz. merolae* (Fig. 8B). In *Cd. caldarium* and *G. partita*, two of the three and both *Hog* genes, respectively, were predominantly expressed in haploid cells (Fig. 8B). TFFs are small, protease-resistant proteins that are co-secreted with mucin molecules in the stomach, localize to the gastric mucous layer, and inhibit acidification of surface epithelial cells in mammals (Aihara et al., 2017). Of the 16 *TFF*-like genes (K+TFF and TFF) encoded in the *Cc. yangmingshanensis* genome (15 in *Cz. merolae* and 13 in *Cd. caldarium*), seven were predominantly expressed in diploid cells, while five were predominantly expressed in haploid cells (Supplementary Data Set 7).

However, proteome analyses of *Cc. yangmingshanensi*s showed that many of these proteins are predominantly expressed in the diploid phase (Fig. 8A). Furthermore, in *Cz. merolae* and *Cd. caldarium*, 13 *TFF*-like genes in each species were predominantly expressed in diploid cells (Fig. 8B; Supplementary Data Sets 8 and 9). Thus, these proteins are likely to function primarily in the diploid phase. The evolutionary origin of *TFF*-like genes remains unclear; however, given that *TFF*s are found only in vertebrates outside of Cyanidiophyceae, it is possible that they arose through convergent evolution to protect cells from low environmental pH in Cyanidiophyceae.

Eisosomes are trough-shaped invaginations of the cell membrane and are found across various eukaryotic lineages (Lee et al., 2015). Recent studies have identified eisosomes as integral components of major stress response pathways in the yeast *S. cerevisiae* (Lanze et al., 2020), although their roles in other organisms remain unclear. Among microalgae, eisosomes have been observed only in cell-walled species, including cell-walled *Cz. merolae* YNP 1A and *G. sulphuraria* CCMEE 5587.1 (Lee et al., 2015). Consistent with this observation, eisosomes were detected in cell-walled diploid cells, but not in cell wall-less haploid cells, of *Cd. caldarium*, *Cc. yangmingshanensis*, and *Cz. merolae* (Fig. 2C; Supplementary Fig. S1).

In *S. cerevisiae*, Pil1 and Lsp1, which are membrane-sculpting Bin/amphiphysin/Rvs (BAR) domain proteins, have been identified as core components of eisosomes (Walther et al., 2006; Olivera-Couto et al., 2011). Of the five *BAR* genes in the *Cc. yangmingshanensis* genome (six in *Cz. merolae* and *Cd. caldarium*), one gene was predominantly expressed in diploid cells, whereas two others were predominantly expressed in haploid cells (Fig. 8A), a result supported by proteome analyses (Fig. 8A). A similar expression pattern of *BAR* genes was also observed in *Cz. merolae* and *Cd. caldarium* (Fig. 8B). Thus, the *BAR* genes predominantly expressed in the diploid phase may represent a candidate involved in eisosome formation. In contrast, in *G. partita*, two of the ten *BAR* genes were predominantly expressed in haploid cells, whereas the others showed similar expression levels in both diploid and haploid cells (Fig. 8B), suggesting that these genes may be related to both cell wall invaginations in the diploid phase and/or in other membrane invaginations in the haploid phase.

### Differentially expressed genes underlying differences in the mechanisms of cell division between haploid and diploid cells

Actin is involved in various cellular processes, including cellular motility and cytokinesis in eukaryotes (Blanchoin et al., 2014). Notably, previous studies showed that cytokinesis in the cell-walled *Cc. yangmingshanensis* RK-1 (as demonstrated in this study to be diploid) is accompanied by actin ring formation at the division site, whereas in the cell wall-less *Cz. merolae* 10D (demonstrated above to be haploid), cytokinesis occurs without actin ring formation (Suzuki et al., 1995; Takahashi et al., 1998). In *Cz. merolae* 10D, the endosomal sorting complex required for transport III (ESCRT-III), which is involved in cytokinesis in archaea and animal cells, was shown to mediate cytokinesis (Yagisawa et al., 2020). These findings suggest that distinct cytokinesis mechanisms operate in the cell-walled diploid and cell wall-less haploid cells of Cyanidiophyceae.

Consistent with these observations, in *Cc. yangmingshanensis, Cz. merolae* and *Cd. caldarium*, a single-copy actin gene and many other genes encoding actin-associated proteins are predominantly expressed in diploid cells (Fig. 9A and B). Proteome analyses of *Cc. yangmingshanensis* also showed the predominant expression of these proteins in the diploid phase (Fig. 9A). Labeling of F-actin with phalloidin revealed the formation of actin rings at the division plane in diploid cells, but not in haploid cells (Fig. 9C; Supplementary Fig. S10A). In agreement with this, haploid cells—but not diploid cells—were able to proliferate in the presence of the actin polymerization inhibitors cytochalasin B and latrunculin B (Fig. 9D; Supplementary Fig. S10B), indicating that diploid cytokinesis, but not haploid cytokinesis, depends on F-actin formation at the division site. In *G. partita*, which possesses four actin genes, *ACT3* (orthologous to the *Cc. yangmingshanensis* actin gene) and *ACT2* are required for cytokinesis in diploid cells and for cell motility in haploid cells, respectively (Hirooka et al., 2022).

**Figure 9.**
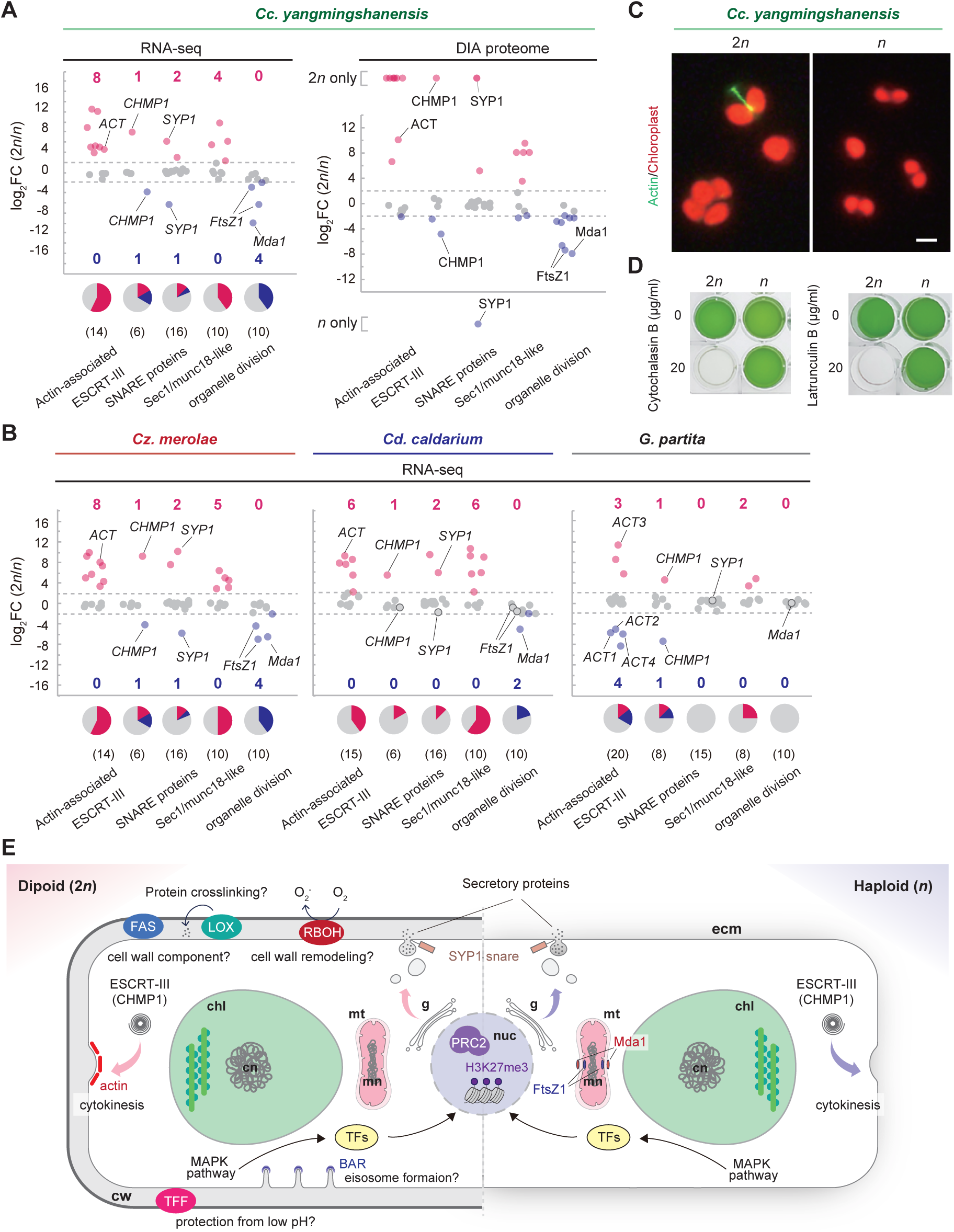
Comparison of transcriptomes of cell division-related genes in Cyanidiophyceae. **A)** Ratios of mRNA and corresponding protein abundance between 2*n* and *n* cells for genes encoding actin, actin-associated proteins, ESCRT-III components, SNARE proteins, and other functional categories related to cell division in *Cc. yangmingshanensis*. Numbers in parentheses below the pie chart indicate the number of genes in each category. See also Supplementary Data Set 7 for details. **B)** Ratios of mRNA abundance between 2*n* and *n* cells for genes encoding actin, actin-associated proteins, ESCRT-III components, SNARE proteins, and other functional categories related to cell division in *Cz. merolae, Cd. caldarium,* and *G. partita*. Numbers in parentheses below the pie chart indicate the number of genes in each category. See also Supplementary Data Sets 8-10 for details. **C)** Actin filaments visualized using Alexa Fluor™ 488 phalloidin in 2*n* and *n* cells (green, phalloidin fluorescence indicating actin filaments; red, chloroplast fluorescence). Scale bar: 2 µm. **D)** Comparison of sensitivity to actin polymerization inhibitors (cytochalasin B or latrunculin B) between the 2*n* and *n* cells of *Cc. yangmingshanensis*. Photographs show 2*n* and *n* cells after 7 days of cultivation in the presence or absence of cytochalasin B or latrunculin B. **E)** Schematic representation of the biological features of 2*n* and *n* cells. *chl*, chloroplast; *cn*, chloroplast nucleoid; *cm*, cell membrane; *cw*, daughter cell wall; *ecm*, extracellular matrix; *g*, golgi apparatus; *mt*, mitochondrion; *mn*, mitochondrial nucleoid; *nuc*, nucleus.

Among the ESCRT-III components, two paralogous *CHMP1* genes encoded in the genomes of *Cc. yangmingshanensis*, *Cz. merolae*, *Cd. caldarium*, and *G. partita* exhibited phase-specific expression: one was predominantly expressed in diploid cells, and the other in haploid cells (Fig. 9A and B). The only minor exception was that, in *Cd. caldarium*, one *CHMP1* gene was also predominantly expressed in diploid cells, whereas the other showed slightly higher expression in haploid cells (log_2_FC(2*n*/*n*) = −0.71, below the threshold for differential expression defined in this study) (Fig. 9B). In *Cc. yangmingshanensi*s, proteome analyses also showed these phase-specific expression patterns of CHMP1 proteins (Fig. 9A). A previous study showed that the *CHMP1* gene expressed in haploid cells localizes at the division site during cytokinesis in *Cz. merolae* 10D (Yagisawa et al., 2020). Although the localization and function of the CHMP1 paralog expressed in diploid cells remain unclear, it is likely that distinct ESCRT-III complexes are involved in cytokinesis in the cell-walled diploid and cell wall-less haploid cells.

Notably, the genomes of *Cc. yangmingshanensis*, *Cz. merolae*, and *Cd. caldarium* do not encode any myosin genes, suggesting that the actin ring involved in diploid cytokinesis functions in the absence of myosin via a mechanism that remains to be elucidated. In *Galdieria*, a single myosin-family gene is encoded in the genome; however, we previously showed that the corresponding protein is not required for cytokinesis (Hirooka et al., 2022).

The soluble N-ethylmaleimide-sensitive factor attachment protein receptor (SNARE) proteins mediate membrane fusion between organelles and between vesicles and the plasma membrane, and their assembly is primarily regulated by Sec1/Munc18-like (SM) proteins (Sudhof and Rothman, 2009). Of the 16 genes encoding SNARE proteins in each genome of *Cc. yangmingshanensis*, *Cz. merolae,* and *Cd. caldarium*, one of the two paralogous *SYP1* genes was predominantly expressed in diploid cells and the other in haploid cells (Fig. 9A and The only minor exception was that, in *Cd. caldarium*, one *SYP1* gene was also predominantly expressed in diploid cells, whereas the other showed slightly higher expression in haploid cells (log_2_FC(2*n*/*n*) = −1.73, below the threshold for differential expression defined in this study) (Fig. 9B). Additionally, of the ten SM genes encoded in each genome of *Cc. yangmingshanensis, Cz. merolae,* and *Cd. caldarium*, four were predominantly expressed in diploid cells (five in *Cz. merolae* and six in *Cd. caldarium*) (Fig. 9A and B). In *Cc. yangmingshanensi*s, proteome analyses confirmed that the two SYP1 proteins were exclusively expressed in the haploid and diploid phases, respectively, whereas five of the SM proteins were predominantly expressed in diploid cells (Fig. 9A). In *Galdieria,* the genome encodes a single *SYP1* gene, which is expressed at similar levels in both diploid and haploid cells, and eight *SM* (SEC) genes, two of which are predominantly expressed in diploid cells (Fig. 9B).

Members of the SYP1 family generally localize to the plasma membrane (PM) and mediate membrane fusion between the PM and secretory vesicles. In *Arabidopsis thaliana*, most *SYP1* genes exhibit tissue-specific expression, contributing to the complexity of plant development through membrane trafficking (Enami et al., 2009). KNOLLE (SYP111), a member of the SYP1 family, interacts with the membrane-associated SM protein KEULE (SEC11) at the cell plate and plays a key role in cytokinesis in *A. thaliana* (Waizenegger et al., 2000). In the liverwort *Marchantia polymorpha*, SYP12A, a member of the SYP1 family, is required for cell plate formation during cytokinesis, functioning similarly to KNOLLE (Kanazawa et al., 2020).

Based on these findings, in *Cyanidiococcus, Cyanidioschyzon,* and *Cyanidium*, the phase-specific expression of the two *SYP1* paralogs may be associated with differences in the secretory pathway, which in turn may lead to the formation of distinct extracellular matrices between diploid and haploid cells. Additionally, one or more SM proteins that are highly expressed in diploid cells may interact with SYP1 and contribute to the construction of a nascent cell wall during cytokinesis. In *Galdieria*, although only a single *SYP1* gene is present and expressed in both diploid and haploid cells, the difference in the expression patterns of *SM* genes likely results in a difference in extracellular matrix formation between the diploid and haploid phases.

Finally, while it is unclear how this impact manifests as phenotypic differences, we found that two types of *FtsZ1* genes (each a single-copy gene) and the *Mda1* gene (also a single-copy gene), but not the dynamin-related gene (*DRP3*), which are responsible for mitochondrial division (Nishida et al., 2003; Nishida et al., 2007), are predominantly expressed in the haploid phase in *Cc. yangmingshanensis*, *Cz. merolae*, and *Cd. caldarium* (Fig. 9A and B). In *Cc. yangmingshanensis*, proteome analyses also showed the predominant expression of these proteins in the haploid phase (Fig. 9A). These results suggest that the haploid and diploid phases may use different mechanisms for mitochondrial division, though further analysis is needed. Regarding *Galdieria*, which possesses a net-like mitochondrion unlike other cyanidiophycean genera with a single disc-shaped mitochondrion, it has lost the mitochondrial *FtsZ* gene during evolution, and the *Mda1* as well as the *DRP3* gene exhibited no differences in expression between the haploid and diploid phases (Fig. 9B; Supplementary Data Set 10).

In summary, multiple candidate genes involved in the phenotypic differences between diploid and haploid phases in cyanidiophycean algae were identified (Fig. 9E). In particular, several genes, including those encoding the MADS-box transcription factor, MAPK cascade components, actin, and the ESCRT-III subunit CHMP1, exhibited similar phase-specific expression patterns across Cyanidiophyceae. These results suggest that the functions of these genes were retained from the last common ancestor of Cyanidiophyceae. The functions of these proteins in Cyanidiophyceae may reflect their ancestral features in Archaeplastida, which will be elucidated by further functional analyses through genetic modifications in Cyanidiophyceae.

## Discussion

### Revisiting Cyanidiophyceae following the discovery of their sexual life cycle

Sexual reproduction had not been observed in unicellular red algae, despite their long history of research. Recently, we discovered a sexual life cycle in *Galdieria*, the earliest-branching genus in the unicellular red algal class Cyanidiophyceae (Hirooka et al., 2022). In this study, we further elucidate that three additional genera of Cyanidiophyceae—*Cyanidium*, *Cyanidiococcus*, and *Cyanidioschyzon*—also have a sexual life cycle, alternating between the cell-walled diploid and the cell wall-less haploid (Fig. 10).

**Figure 10.**
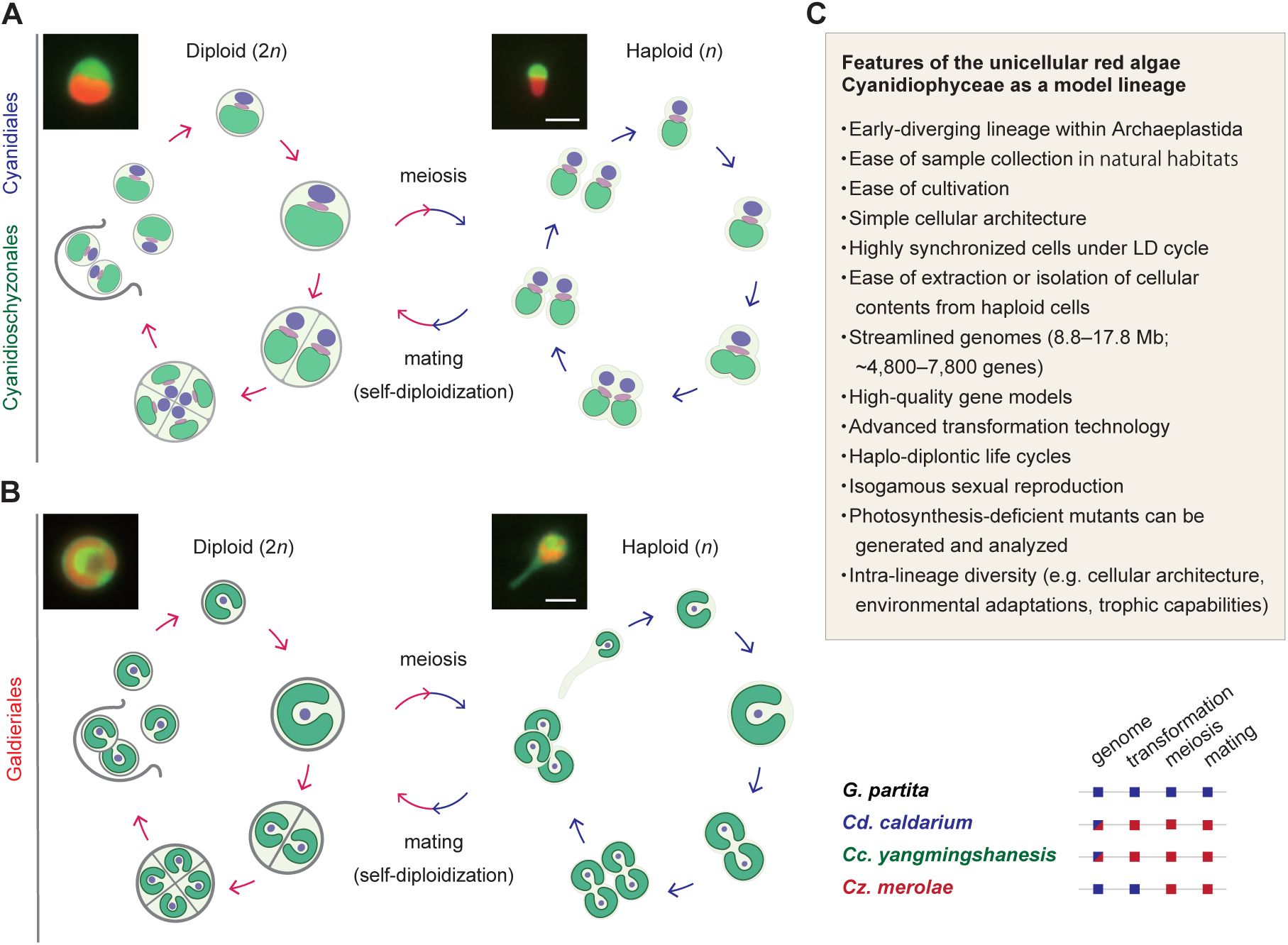
The sexual life cycle and model-lineage characteristics of the unicellular red algae Cyanidiophyceae. **A and B)** Haplo-diplontic life cycle of *Cyanidium* (Cyanidiales), *Cyanidiococcus* and *Cyanidioschyzon* (both Cyanidioschyzonales) **(A)**, and *Galdieria* (Galdieriales) **(B)**, showing the proliferation modes 2*n* and *n* cells. These cyanidiophycean algae can undergo sexual reproduction under laboratory conditions, enabling the generation of stable diploid transformants through mating between haploid transformants. The upper panels show transformants expressing the fluorescent protein Venus in the cytoplasm. Scale bar: 2 µm. **C)** Summary of features and experimental techniques available for Cyanidiophyceae. In this study, we established that all four axenically cultivable genera of Cyanidiophyceae now have access to high-quality genomic resources, genetic manipulation methods, and protocols for inducing meiosis (to generate *n* cells) and mating (to generate heterozygous 2*n*). Blue boxes indicate presence based on previous studies, and red boxes indicate presence based on this study.

Thus far, inconsistencies between morphological and molecular phylogenetic classifications have caused systematic problems in the classification of Cyanidiophyceae. The discovery of the sexual life cycle has revealed that the traditional classification that only *Cyanidioschyzon* among Cyanidiophyceae lacks a cell wall and proliferates by binary fission, whereas *Cyanidiococcus* and *Cyanidium* possess a cell wall and proliferate by forming four daughter cells within the mother cell (Fig. 1, B and C), has turned out to be incorrect. As shown in this study, the diploid forms, as well as the haploid forms, of these three genera are morphologically indistinguishable from one another under light microscopy (Fig. 2B).

Instead, based on current knowledge, the major differences among the three genera, classified by molecular phylogenetic analysis (Park et al., 2023), include large variations in genome size (Fig. 1B) (Cho et al., 2023) and partial differences in gene content (Fig. 3A). The thicker cell wall of *Cyanidium*, compared to the other two genera (Fig. 2C; Supplementary Fig. S1), as well as its lower optimal temperature and pH (Fig. 4F), may serve as distinguishing characteristics for *Cyanidium* in relation to *Cyanidiococcus* and *Cyanidioschyzon*, although further investigation of additional strains is needed.

Furthermore, it has become clear that *Cz. merolae* 10D, the most extensively studied cell wall-less strain among Cyanidiophyceae (Kuroiwa et al., 2018; Miyagishima and Tanaka, 2021), should be recognized as a haploid clone. With this clarification, the long-standing mystery from previous genome analyses of *Cz. merolae* 10D—namely, the presence of numerous genes that show no detectable mRNA expression despite being conserved across other eukaryotic lineages (such as genes encoding actin and its regulatory proteins) (Matsuzaki et al., 2004)—can now be reasonably explained. This study reveals that many of these genes are, in fact, specifically expressed in the diploid phase (Fig. 7A; Supplementary Fig. S9).

In this study, our field sampling in Japanese sulfuric hot springs revealed that only the cell-walled diploid forms of *Cyanidiococcus* and *Galdieria* were present in the blue-green algal mats attached to submerged rocks, and in endolithic habitats and mud deposits above the water surface (Fig. 4A; Supplementary Fig. S4A–F). Furthermore, we reanalyzed recently published diurnal metatranscriptomic (RNA-seq) data from Yellowstone National Park, USA—specifically, cyanidiophycean mats in a stream dominated by *Cz. merolae* sequences and terrestrial mats on mud, where *G. yellowstonensis* sequences were second in abundance after *Cz. merolae* (Stephens et al., 2024). Even in these datasets, diploid-specific genes (identified in this and previous studies) were detected (Supplementary Fig. S11A), whereas haploid-specific gene sets showed almost no detectable expression (Supplementary Fig. S11B). Supporting these results, microscopic observations of the *Cz. merolae* mat in the stream revealed only cell-walled round cells (Bhattacharya et al., 2025).

The cell wall-less *Cz. merolae* strain was originally isolated from hot spring samples collected in Naples, Italy, after propagation of algae in the samples in an inorganic, nutrient-rich medium at pH 1.5 (De Luca et al., 1978). The *Cz. merolae* 10D was also re-isolated from a similar sample (Toda et al., 1995). Thus, these strains were most likely originally cell-walled diploids in nature, which underwent meiosis and produced cell wall-less haploids during cultivation in nutrient-rich media prior to the isolation of haploid clones.

Thus, to date, there have been no confirmed observations of the haploid phase of Cyanidiophyceae in natural habitats. When, where, and under what environmental conditions haploid cells are formed remain open questions for future investigation, although in this study they were generated at low frequency in an inorganic, nutrient-rich medium with a lower pH (1.0 or 0.75) than the standard cultivation conditions for Cyanidiophyceae (∼pH 2.0).

### Utility of Cyanidiophyceae as a model lineage for studying eukaryotic evolution, photosynthesis, life cycles, and phenotypic diversification

In this and our recent (Hirooka et al., 2022) study, taking advantage of the lack of a cell wall in the haploid phase of Cyanidiophyceae, PEG-mediated DNA introduction—originally developed for genetic modification of the cell wall-less strain *Cz. merolae* 10D—has now been successfully applied to all four axenically cultivable genera of Cyanidiophyceae.

Together with genomic and transcriptomic data, this genetic manipulation technology provides a powerful platform for studying the evolution and minimal requirements for the construction and function of photosynthetic eukaryotic cells, as well as the adaptation and colonization of eukaryotes into new environments and other biological processes, based on the following features.

- Cyanidiophyceae possess the simplest cellular and genomic structures among known photosynthetic eukaryotes. These cells contain only a minimal set of membrane-bound organelles (Kuroiwa et al., 2018), and their nuclear genomes encode ∼4,800–7,800 genes. Their cell cycles can be tightly synchronized with the diel cycle, even in natural habitats (Suzuki et al., 1994; Fujiwara et al., 2009; Jong et al., 2021; Stephens et al., 2024). Additionally, haploid cells of Cyanidiophyceae lack a rigid cell wall, which facilitates the extraction and isolation of cellular contents, including intact organelles, as demonstrated in *Cz. merolae* 10D (Miyagishima et al., 1999; Yagisawa et al., 2009). The simple genome facilitates comprehensive omics such as transcriptomics and proteomics. For example, DIA proteome analyses of *Cc. yangmingshanensis* detected more than 90% of the predicted proteins, providing near-complete coverage of the expressed proteome (Supplementary Data Set 7). Taken together, these features enable studies on cell cycle and organelle biology, including inter-organelle studies.
- A variety of genetic modification techniques can be applied to the streamlined genomes of Cyanidiophyceae. Because most proteins are encoded by single-copy genes (except for *Galdieria*; Fig. 3C), phenotypic analyses are relatively straightforward. Marker recycling enables editing of multiple loci (Fig. 5, G-I; Supplementary Fig. S5 G-J), and haploid clones can be converted into diploids as needed (Fig. 10A and B). Strategies developed for *Cz. merolae* 10D—such as gene knockouts, epitope tagging, inducible expression, inducible knockdown, and CRISPR/Cas9-based editing—are in principle applicable to all strains. Currently, generating transformants of *Cc. yangmingshanensis* and *Cd. caldarium* takes over a month; however, this process could be shortened by referring to a recently optimized protocol that enables transformant preparation in *Cz. merolae* 10D within two weeks (Villegas-Valencia et al., 2025). Thus, while further optimization is needed, these technologies are expected to facilitate functional genomic analyses across Cyanidiophyceae.
- Both diploid and haploid cells undergo stable asexual proliferation, representing a haplo-diplontic life cycle. This feature allows straightforward preparation of samples and direct comparison between the two phases. In addition, a recent study reported that 50% of protists distributed across the entire eukaryotic evolutionary tree possess a haplo-diplontic life cycle, suggesting that this may be an ancestral feature of eukaryotes (Rizos et al., 2024). Thus, studying the life cycles of Cyanidiophyceae may provide insights into the characteristics and evolution of the original eukaryotic life cycle.
- Photosynthesis-deficient mutants can be generated and analyzed. The red algal photosynthetic apparatus exhibits intermediate characteristics between those of cyanobacteria and Viridiplantae (Allen et al., 2011) and contains lineage-specific components of unknown function (some also present in organisms with secondary red plastids) (Sturm et al., 2013; Lee et al., 2019). However, molecular genetic studies of photosynthesis-related genes have not been feasible in red algae, limiting their functional analyses. We recently succeeded in generating knockout mutants of genes involved in photosynthetic pigment biosynthesis in *G. partita* (Hirooka et al., 2022). This was possible because *Galdieria* can uniquely grow photoautotrophically, mixotrophically, and heterotrophically, unlike strictly photoautotrophic genera such as *Cyanidium*, *Cyanidiococcus*, and *Cyanidioschyzon* (Gross and Schnarrenberger, 1995; Barbier et al., 2005). Notably, a transgenic *Cz. merolae* expressing a *Galdieria* plasma membrane sugar transporter showed heterotrophic growth in glucose-supplemented medium (Fujiwara et al., 2019). Thus, these features make Cyanidiophyceae a valuable model for uncovering unknown mechanisms of photosynthesis.
- Cyanidiophyceae exhibit intra-lineage diversity in environmental adaptations, trophic capabilities, cellular architectures, and organelle inheritance. After diverging from the common ancestor of other red algae around 1.5 billion years ago, Cyanidiophyceae diversified (Yoon et al., 2004; Cho et al., 2023; Park et al., 2023; Huang et al., 2024). Today, many strains exist, each adapted to distinct, isolated hot, acidic springs—or even to distinct microenvironments within the same location. For example, *G. phlegrea* is adapted to endolithic layers (Qiu et al., 2013). Further, as shown in this study, even among thermo-acidophilic strains, optimal growth conditions and metal tolerance vary (Cho et al., 2023) (Fig. 4F; Supplementary Fig. S4D). Some species have secondarily adapted to mesophilic environments (e.g., the genus *Cyanidiofrigus*; (Huang et al., 2024)), or even to neutral environments (e.g., three genera: *Gronococcus*, *Cavernulicola*, and *Sciadococcus*; (Park et al., 2023)), although their axenic cultures have not yet been established and genomic information is still lacking. Remarkably, 18S rDNA sequences related to Cyanidiophyceae have been detected in global ocean samples (*Tara* Oceans project; https://www.gbif.org/species/334), raising the possibility that additional, yet-undiscovered diversity may exist within Cyanidiophyceae. Thus, comparative analyses of the currently cultivable cyanidiophycean strains exhibiting diverse environmental adaptations, together with the future establishment of axenic cultures of cyanidiophycean algae inhabiting mesophilic environments, are expected to facilitate studies of adaptive evolution in Cyanidiophyceae. In addition, one of the major differences within Cyanidiophyceae is trophic capability: *Galdieria* is capable of heterotrophic growth, and in some strains, the chloroplast (plastid) can reversibly lose the photosynthetic apparatus and thylakoid membrane, whereas other genera are obligate photoautotrophs, possibly having lost this capacity altogether (Gross and Schnarrenberger, 1995; Barbier et al., 2005; Yamashita et al., 2025). Furthermore, Cyanidiophyceae exhibit diversity in other traits, such as the morphology of mitochondria and vacuoles (Fig. 1C) (Albertano et al., 2000), and patterns of mtDNA inheritance (Fig. 6I; Supplementary Fig. S8). Comparative genomic analyses have suggested that gene losses, horizontal gene transfers (HGTs)—including those encoding heavy metal detoxification enzymes and heat shock proteins—and de novo gene emergence, all occurring in a lineage-or strain-specific manner, likely underlie the phenotypic diversification observed across Cyanidiophyceae (Van Etten et al., 2023). To further clarify the genetic bases of phenotypic diversification, functional approaches—such as gene knockout or gene transplantation (for which this study developed a method applicable to all culturable strains), exemplified by the expression of a *Galdieria* sugar transporter in *Cyanidioschyzon* in a previous study (Fujiwara et al., 2019) —will be particularly useful. Collectively, these features establish Cyanidiophyceae as a model lineage for elucidating phenotypic diversification in eukaryotes.

### The Cyanidiophyceae as a basis for understanding the origin of complex multicellular sexual life cycles and associated phenomena in Archaeplastida

Sexual reproduction has been observed in multicellular red algae, certain lineages of both multicellular and unicellular green algae, and land plants. The sexual life cycle of Cyanidiophyceae is the only one known among unicellular red algae and represents the earliest-diverging example found within Archaeplastida to date (Fig. 1A). Thus, it would provide crucial insights into the evolution of multicellular sexual life cycles with cellular differentiation, which independently arose in red algae and Viridiplantae (Fig. 1A), as well as into the broader evolution of such systems in eukaryotes.

In this study, we found that 159 (3.3%) and 91 (1.9%) protein-coding genes in the nuclear genome (4,884 protein-coding genes) are predominantly expressed in the diploid and haploid phases, respectively, in *Cc. yangmingshanensis* (Fig. 7A). Some of these genes are reasonably correlated with phenotypic differences between the two phases, such as the presence or absence of a cell wall and the requirement for actin in cell division (Fig.9E).

Furthermore, the set of differentially expressed genes includes members of transcription factor families that are known in plants to be involved in the formation of reproductive structures and other developmental processes, and the number of such genes is considerably smaller compared to plants (Fig. 7D). Similar results were also observed in *Cz. merolae*, *Cd. caldarium,* and *G. partita* (Fig. 7 E). In addition, most of these haploid-or diploid-predominant genes are marked with H3K27me3 in the opposite phase (Fig. 7 B and C).

H3K27me3 is a histone modification catalyzed by the Polycomb Repressive Complex 2 (PRC2), which mediates gene silencing during differentiation and development in multicellular organisms such as animals and plants. In contrast, the PRC2 complex is absent in the yeasts *Saccharomyces cerevisiae* and *Schizosaccharomyces pombe*, which previously led to the assumption that PRC2 and H3K27me3 may have evolved alongside the emergence of multicellularity to regulate cellular differentiation. However, components of PRC2 and its associated mark (H3K27me1/-me2/-me3) have been later identified in several lineages of unicellular eukaryotes (Shaver et al., 2010; Mikulski et al., 2017; Frapporti et al., 2019; Zhao et al., 2021; Petroll et al., 2025b), suggesting that yeasts secondarily lost PRC2. To date, the function of H3K27me3 has been investigated in only a few unicellular eukaryotes. In the diatom *P. tricornutum* (a diploid clone) (Filloramo et al., 2021) and in *Cz merolae* 10D (a haploid clone), H3K27me3 predominantly marks transposable elements (TEs) and repeat sequences, but is also found on certain protein-coding genes (Hisanaga et al., 2023). In *P. tricornutum*, genes involved in cell morphological changes were found to be marked by H3K27me3, and knockout of the *enhancer of zeste* (*E(z)*) gene, which encodes the catalytic component of the PRC2 complex, abolished normal cell morphology, suggesting that H3K27me3 regulates cell differentiation in this unicellular organism (Zhao et al., 2021).

However, unlike in *P. tricornutum*, knockout of *E(z)* gene in *Cz. merolae* 10D resulted in the expression of TEs, but no obvious changes in cellular morphology were observed (Hisanaga et al., 2023). Thus, in addition to H3K27me3, other histone modifications and/or the expression of specific transcription factors may also be required for the phenotypic differences between the haploid and diploid phases.

KNOX and BELL-related transcription factors are expressed in gametes and are involved in the haploid-to-diploid transition in *Chlamydomonas reinhardtii* and *Marchantia polymorpha* (Lee et al., 2008; Hisanaga et al., 2021), and more recently have also been shown to be required for this transition in *Galdieria* (Hirooka et al., 2022), which may reflect their ancestral roles in the Archaeplastida. However, in this study, we were not able to detect BELL-related or KNOX proteins in either haploid or diploid cells of *Cc. yangmingshanensis* by proteome analysis, although KNOX mRNA was predominantly detected in the haploid phase of *Cyanidioschyzon* and *Cyanidiococcus* (Fig. 7D and E). These results raise the possibility that the haploid cells examined in this study are likely in a vegetative state and may transform into as-yet-unknown gametes expressing KNOX and BELL proteins upon a certain stimulus. In this regard, among the *Galdieria* haploid cell population, cells before cell division phase formed tail-like membrane protrusions in which actin and microtubules were organized (tadpole-shaped cells), and these cells migrated across surfaces at approximately 25 µm/min (Hirooka et al., 2022). In contrast, the haploid cells of *Cz. merolae* 10D moved much more slowly, at only around 4 µm/min (Ohnuma et al., 2011). On the other hand, a recent study showed that increasing the salt concentration of the *Cz. merolae* culture medium induces, within 10 min, the formation of protrusions from part of the cell membrane containing actin filaments on the cytoplasmic side (termed “tentacles”), which in turn appear to accelerate cell motility (Maschmann et al., 2020). Thus, if vegetative haploids can be induced to form gamete through such a process, it will be important in future studies to distinguish and analyze these states separately.

Regarding organelle DNA inheritance upon diploid formation through mating between two haploids, we found that, except for mtDNA in *Galdieria*, both mtDNA and cpDNA are initially inherited from both mating partners, resulting in a heteroplasmic state. As the diploid cells proliferate, they transition to a homoplasmic state in which only one parental DNA type remains, and which type is retained appears to be determined by chance (Fig. 6I; Supplementary Fig. S8). This mode of inheritance is observed in the mitochondrial DNA of budding yeast and is thought to result from the random replication and segregation of the multicopy mtDNA into daughter cells during the proliferation of diploid cells following the mating of haploids (Birky, 1995; Roussou et al., 2024). Although it remains unclear whether similar mechanisms operate in Cyanidiophyceae, the results suggest that in isogamous Cyanidiophyceae, two distinct mechanisms may exist: one that leads to a homoplasmic state containing organelle DNA from either mating partner in a random manner, and another that imposes a bias to ensure that only organelle DNA derived from a specific type of parent (mating type) is retained. The latter mechanism appears to be specific to the inheritance of mtDNA in *Galdieria* in Cyanidiophyceae. Whether this mechanism for mtDNA in *Galdieria* operates before or during mating, or during the early stages of diploid proliferation, requires further investigation.

Further elucidation of the mechanisms underlying sexual reproduction in Cyanidiophyceae—including those that regulate phenotypic transitions between the haploid and diploid phases, as well as those governing organelle DNA inheritance—will require observation of both mating and meiosis, which we were unable to achieve in this study due to their substantially low frequency in our cultivation conditions. Thus, further studies are needed to identify conditions that increase the frequency of mating and meiosis.

## Materials and methods

### Algal strains and culture conditions

Many cyanidiophycean algal strains have been maintained in stock centers, such as the Microbial Culture Collection at the National Institute of Environmental Studies (NIES), the Sammlung von Algenkulturen Culture Collection of Algae (SAG), the Culture Collection of Cryophilic Algae (CCCryo), and the Algal Collection University Federico II (ACUF).

However, some cyanidiophycean strains are not clonal cultures and are contaminated with other strains. For example, *G. sulphuraria* 3377 was isolated from a *Cz. merolae* culture as a green colony that arose on glucose-supplemented plates in the dark (Lopez-Portillo Masson et al., 2025). Likewise, *Cz. merolae* Soos was isolated from a *G. phlegrea* Soos culture as a strict photoautotroph that outgrew the *Galdieria* strain under photoautotrophic conditions (Rossoni et al., 2019). Due to contamination issues in some strains, caution is required when using cyanidiophycean algal cultures. Therefore, in this study, single clones of all strains obtained from stock centers were reisolated prior to analyses. Thus, in this study, single clones of all strains obtained from stock centers were reisolated prior to analyses.

*Cc. yangmingshanensis* N3110 (NIES-4659) was isolated from a sample collected in an acidic hot spring in Hakone, Kanagawa Prefecture, Japan, and is deposited in the NIES Collection. The other *Cyanidiococcus* strains (heterozygous diploids; CcyaKS-1, CcyaKS-5, CcyaKS-11, TS5, TS6, TH2, TH3, and TH12) were isolated from samples collected in acidic hot springs in Kusatsu, Gunma Prefecture (Sunada et al., 2025), or Tsukahara, Oita Prefecture, Japan, and have been stocked at the Symbiosis and Cell Evolution Laboratory, National Institute of Genetics (NIG). *Cz. merolae* MS1 (heterozygous diploid) was isolated from a sample collected in an acidic hot spring in Yellowstone National Park, Wyoming, USA, and is stocked at the Arizona Center for Algae Technology and Innovation, Arizona State University, and at the Symbiosis and Cell Evolution Laboratory, NIG. *Cd. caldarium* NIES-551 (heterozygous diploid) and *Cz. merolae* 10D (NIES-3377; haploid) were obtained from the NIES Collection. *Cd. caldarium* SAG-16.91 (heterozygous diploid) was obtained from the SAG. *G. partita* NBRC102759 (heterozygous diploid) was obtained from the Biological Resource Center, NITE (NBRC). The haploid strain *G. partita* NBRC102759 N1 (Hirooka et al. 2022), derived from *G. partita* NBRC102759, is stocked at the NIG.

These strains were maintained in 20 mL of MA medium at pH 2.0 (for *Cd. caldarium* NIES-551, *Cc. yangmingshanensis* NIES-4659, other *Cyanidiococcus* strains, *Cz. merolae* 10D, *Cz. merolae* MS1, and *G. partita* NBRC102759) or at pH 1.0 (for *G. partita* NBRC102759 N1) in 25-cm² tissue culture flasks (TPP Techno Plastic Products, Trasadingen, Switzerland), statically in a 2% CO_2_ incubator at 37°C for *Cd. caldarium* NIES-551 and at 42°C for the other strains under continuous light (30 μmol photons m^−2^ s^−1^).

To generate haploid cells from the diploid clones (*Cd. caldarium* NIES-551, *Cc. yangmingshanensis* NIES-4659, and *Cz. merolae* MS1), cells in MA medium at pH 2.0 were transferred to 1 mL of MA medium at pH 1.0 (or pH 0.75 for *Cd. caldarium*) in a 24-well plate (1 mL/well). The cells were cultivated statically in a 2% CO_2_ incubator at 37°C for *Cd. caldarium* and at 42°C for the other strains under continuous light (30 μmol photons m^−2^ s^−1^) for 1–3 weeks. In the resulting culture, cell-wall-less haploid cells were generated, mixed with cell-walled diploid cells. A single haploid cell was isolated and transferred into 1 mL of IMA (iron-rich MA) medium at pH 0.75 for *Cd. caldarium* and at pH 1.2 for the other strains. The IMA medium contained 20 mM (NH_4_)_2_SO_4_, 4 mM MgSO_4_·7H_2_O, 4 mM KH_2_PO_4_, 1 mM CaCl_2_, 5 mM FeSO_4_·7H_2_O, and trace elements as described in Minoda et al. (2004). The isolated haploid cell was cultured in one well of a 24-well plate (1 mL/well) statically in a 2% CO_2_ incubator under continuous light to generate a clonal haploid population. The haploid strains are stocked at the Symbiosis and Cell Evolution Laboratory, NIG. Each haploid clone of the respective strains is maintained in 20 mL of MA medium at pH 2.0 (for *Cc. yangmingshanensis* NIES-4659 Ha1 and *Cz. merolae* MS1 Ha3) or at pH 0.5 or 0.75 (for *Cd. caldarium* NIES-551 Ha5) in 25-cm^2^ tissue culture flasks, statically in a 2% CO_2_ incubator at 37°C for *Cd. caldarium* NIES-551 Ha5 and at 42°C for the other strains under continuous light (30 μmol photons m^−2^ s^−1^).

To determine the optimal temperature condition, cells were cultured at various temperatures (ranging from 25°C to 45°C). Growing cells were inoculated into 20 mL of MA at pH 0.75 (for *Cd. caldarium* NIES-551 Ha5), pH 1.0 (for *G. partita* NBRC102759 N1), and pH 2.0 (for the other strains) in 25-cm^2^ tissue culture flasks to give an OD_750_ of 0.2. The cells were cultured on a reciprocal shaker at 120 rpm (NR-2; TAITEC) in an incubator (MIR-154-PJ; PHC) under ambient air and continuous light conditions (50 μmol photons m^−2^ s^−1^) for 7 days. To determine the optimal pH condition, cells were cultured in MA medium at seven different pH values (ranging from pH 0.1 to pH 3.0). The pH of the medium was adjusted using H₂SO₄. Growing cells were harvested by centrifugation at 1,500 × *g* for 5 min, then resuspended in 1 mL of the respective medium in each well of a 24-well plate to give an OD_750_ of 0.2. The cells were then cultured statically in a 2% CO_2_ incubator at 37°C for *Cd. caldarium* and *G. partita*, and at 42°C for the other strains under continuous light (30 μmol photons m^−2^ s^−1^) for 7 days.

To determine arsenate (KH_2_AsO_4_) and nickel (NiSO_4_·6H_2_O) tolerance, cells were cultivated in media containing a series of concentrations of the respective metals (ranging from 0 to 50 mM). Growing cells were harvested by centrifugation at 1,500 × *g* for 5 min, then resuspended in 1 mL of the respective medium at pH 0.75 in each well of a 24-well plate to give an OD_750_ of 0.2. The cells were cultured statically in a 2% CO_2_ incubator at 37°C for *Cd. caldarium* and at 42°C for the other strains under continuous light (30 μmol photons m^−2^ s^−1^) for 7 days. OD_750_ values were measured using a spectrophotometer (BioSpectrometer basic; Eppendorf, Germany). Growth rates based on OD_750_ were determined as described by Hirooka et al. (2014).

To determine the sensitivity of the *CP^r^-HSVtk* strain to ganciclovir and 5-FdU, cells were cultivated in media containing a series of concentrations of ganciclovir (ranging from 0 to 2 mg/mL) and 5-FdU (from 0 to 100 μg/mL). Growing cells were harvested by centrifugation at 1,500 × *g* for 5 min, then resuspended in 1 mL of the respective medium in each well of a 24-well plate to give an OD_750_ of 0.2. The cells were then cultured statically in a 2% CO_2_ incubator at 42°C under continuous light (30 μmol photons m^−2^ s^−1^) for 7 days.

To compare the sensitivity to actin polymerization inhibitors between diploid and haploid cells of cyanidiophycean strains, original diploid clones (*Cc. yangmingshanensis* NIES-4659 and *Cz. merolae* MS1) and haploid clones (*Cc. yangmingshanensis* NIES-4659 Ha1, *Cz. merolae* MS1 Ha3, and *Cz. merolae* 10D), grown in MA medium at pH 2.0, were inoculated into 1 mL of MA medium at pH 2.0 supplemented with 20 µg/mL cytochalasin B or latrunculin B to give an OD_750_ of 0.2. For the *Cd. caldarium* strains (*Cd. caldarium* NIES-551 and *Cd. caldarium* NIES-551 Ha5), cells were grown at 37°C in MA medium at pH 0.75 and inoculated into 1 mL of MA medium at pH 0.75 supplemented with 50 µg/mL cytochalasin B or latrunculin B. Stock solutions of cytochalasin B (10 mg/mL; FUJIFILM Wako Pure Chemical Corp., Osaka Japan) and latrunculin B (5 mg/mL; Sigma-Aldrich) were prepared in dimethyl sulfoxide (DMSO). DMSO was used as a control. The respective cells were cultured statically in one well of a 24-well plate in a 2% CO_2_ incubator at 42°C for 1 week (*Cc. yangmingshanensis* and *Cz. merolae*) or at 37°C for 2 weeks (*Cd. caldarium*), under continuous light (30 μmol photons m^−2^ s^−1^).

### Microscopy

To observe cyanidiophycean cells using differential interference contrast (DIC) and fluorescence microscopy, to compare nuclear DNA content, and to visualize actin filaments, cells were cultured statically in 20 mL of MA medium, adjusted to the optimal pH for each strain, in 25-cm^2^ tissue culture flasks. Cultures were maintained in a 2% CO₂ incubator at the optimal temperature for each strain under continuous light (30 μmol photons m^−2^ s^−1^).

To compare nuclear DNA content, a 0.5 mL sample of cultured cells was fixed with 0.1% glutaraldehyde at room temperature for 10 min. The fixed cells were centrifuged at 1,500 × *g* for 5 min, and the cell pellet was resuspended in 1 mL of 50% (v/v) ethanol and incubated at room temperature for 5 min. The cells were then harvested by centrifugation and resuspended in 0.5 mL of 70% (v/v) ethanol. After centrifugation, the cells were resuspended in 0.5 mL of phosphate-buffered saline (PBS), and DAPI was added to a final concentration of 1 µg mL^−1^.

To visualize actin filaments, 0.1 mL of cultured cells was fixed with 2% paraformaldehyde at room temperature for 10 min, then centrifuged at 1,500 × *g* for 5 min. The fixed cells were washed twice with 0.5 mL of PBS, each followed by centrifugation at 1,500 × *g* for 5 min. The cell pellets were resuspended in 1 mL of ice-cold acetone, incubated at −20°C for 10 min, and centrifuged again at 1,500 × *g* for 5 min. The cells were then resuspended in 0.1 mL of PBS, followed by the addition of 2.5 μL of Alexa Fluor^TM^ 488 phalloidin.

Images of the cells were captured using a BX51 upright microscope (Olympus, Tokyo, Japan) equipped with a DP71 digital camera (Olympus). To detect mVenus/Alexa Fluor^TM^ 488 phalloidin, chloroplast (chlorophyll) fluorescence, and DAPI-stained DNA, the NIBA, WIG, and U-MWU filter sets (Olympus) were used, respectively. The fluorescence intensity of DAPI-stained DNA was measured using ImageJ ver. 1.53a (Schneider et al., 2012).

To observe cyanidiophycean cells using transmission electron microscopy, cells were cultured in MA medium at pH 2.0 and 1.0, respectively, in 25-cm^2^ tissue culture flasks on a rotary shaker (120 rpm) in a 5% CO₂ incubator at 42 °C under continuous light (50 μmol photons m^−2^ s^−1^). Preparation of samples and observations were performed as previously described for haploid and diploid cells of *Cc. yangmingshanensis* (Ichinose and Iwane, 2018; Hirooka et al., 2022), with minor modifications and for those of *Cz. merolae*, *Cd. caldarium* and *G. partita* (Yamashita et al., 2025).

### Phylogenetic analysis

The amino acid and nucleotide sequences were aligned using MAFFT ver. 7.212 (Katoh and Standley, 2013), and ambiguous sites were excluded using Gblocks ver. 0.91b (Castresana, 2000). Substitution models were selected using ModelTest-NG ver. 0.1.7 (Darriba et al., 2020) (LG+G for the amino acid sequences of RbcL in Fig. 2A; HKY+G and GTR+G for the nucleotide sequences of *rbcL* in Fig. 4E and Supplementary Fig. S7D, respectively).

Maximum likelihood (ML) analysis and 1,000 pseudoreplicates of bootstrap analysis were performed using RAxML-NG ver. 1.0.3 (Kozlov et al., 2019). Bayesian analyses were performed using MrBayes 3.2.7 (Ronquist et al., 2012) with the optimal substitution models, 1,000,000 generations of Markov chain Monte Carlo iterations, and a burn-in of 25%.

### DNA extraction and sequencing

For PacBio long-read sequencing, haploid cells (*Cc. yangmingshanensis* NIES-4659 Ha1 and *Cd. caldarium* NIES-551 Ha5) were harvested by centrifugation at 2,000 × *g* for 5 min from 10 mL cultures (OD₇₅₀ ∼5.0). High-molecular-weight genomic DNA was extracted from the harvested cells according to Hirooka *et al*. (2022). Following the manufacturer’s instructions, the genomic DNA was subjected to library construction using the SMRTbell Template Prep Kit (Pacific Biosciences of California, Inc., USA) and sequenced using a PacBio RS II or PacBio Sequel II.

For Illumina short-read sequencing, the cells were harvested by centrifugation at 2,000 × *g* for 5 min from 2-mL cultures (OD_750_ ∼5.0). Genomic DNA was extracted from the harvested cells according to Hirooka et al. (2022). The extracted genomic DNA was separately subjected to library construction using the MiSeq Reagent Kit ver. 3 (Illumina, Inc.) for the 300-bp paired-end library and TruSeq DNA PCR-Free Kit (Illumina, Inc.) for the 150-bp paired-end library. The 300-bp and 150-bp paired-end libraries were sequenced using MiSeq and NovaSeq 6000, respectively. The sequence reads were processed using Cutadapt ver. 3.1 to remove low-quality ends (<QV30), adapter sequences, and reads shorter than 50 bp. The trimmed reads were used for subsequent analyses.

### Genomic analyses

For *de novo* assembly of organelle genomes, Illumina short reads of the haploid clones (*Cc. yangmingshanensis* NIES-4659 Ha1 and *Cd. caldarium* NIES-551 Ha5) were assembled using SPAdes ver. 3.15.0 (Bankevich et al., 2012). Genes of the chloroplast and mitochondrial genomes were annotated using GeSeq on Chlorobox (Tillich et al., 2017), with the previously annotated mitochondrial and chloroplast genomes (Cho et al., 2020; Park et al., 2023) as references. For *de novo* nuclear genome assembly, PacBio long reads were mapped to the chloroplast and mitochondrial genomes using Minimap2 ver. 2.17 (Li, 2018), and the mapped reads were removed by BEDTools (Quinlan and Hall, 2010). The remaining long reads were then assembled *de novo* using Canu ver. 2.0 (Koren et al., 2017). The assembled contigs were error-corrected using Pilon ver. 1.2.3 (Walker et al., 2014), with Illumina reads mapped using Bowtie2 ver. 2.3.4.1 (Langmead and Salzberg, 2012). Nuclear genes were predicted using AUGUSTUS (Stanke et al., 2004) by integrating RNA-seq data from both the diploid and haploid, with additional manual curation. Each translated amino acid sequence was used as a BLASTP query against the nr database (E-value < 1×10⁻³) through Blast2GO (Conesa et al., 2005) to assign provisional annotations (Supplementary Data Set 7). Nuclear genes encoding transcription factors (TFs), transcriptional regulators (TRs), and protein kinases (including MAPK components) were identified using iTAK (Zheng et al., 2016).

However, in *Cd. caldarium*, the *BELL*-related TF gene was not identified by iTAK but was detected through manual curation. Secretory proteins, glycosyltransferases, and transmembrane proteins were identified using SignalP 6.0 (Teufel et al., 2022), dbCAN2 metaserver (Zhang et al., 2018), and the Phobius web server (Kall et al., 2007), respectively. In addition, these sequences were subjected to the SUPERFAMILY database ver. 1.75 (Wilson et al., 2007) to verify the presence of specific domains (Sec1/munc18-like, BAR/IMD, Kelch, trefoil, and hedgehog). SNARE, ESCRT-III, and actin-related proteins were identified by BLAST using annotated sequences as queries, as reported in Sanderfoot (2007); Kusdian et al. (2013); Yagisawa et al. (2020).

For comparative genome analysis, genome datasets downloaded from public databases (Supplementary Data Set 5) were used. To identify orthologous genes, gene clustering analysis was performed using OrthoFinder ver. 2.5.4 (Emms and Kelly, 2019) with default parameters. A BLASTP search for amino acid sequences was performed using DIAMOND ver. 2.1.0 (Buchfink et al., 2015).

To identify SNPs and indels, DNA-seq reads were mapped to the reference genome sequences (*Cc. yangmingshanensis* NIES-4659 Ha1, *Cd. caldarium* NIES-551 Ha5, and *Cz. merolae* 10D) using Bowtie2 ver. 2.3.4.1 (Langmead and Salzberg, 2012). Reads mapped to multiple sites were removed using SAMtools ver. 1.8 (Li et al., 2009). SNPs and indels were called with GATK HaplotypeCaller ver. 3.8 (McKenna et al., 2010), and high-confidence variants were identified using GATK VariantFiltration ver. 3.8 (McKenna et al., 2010) with the parameters DP > 30 and QUAL > 50. SNPs and indels were visualized using the Integrative Genomics Viewer (IGV) ver. 2.8.2 (Robinson et al., 2011).

### Pulsed field gel electrophoresis

Cells of *Cd. caldarium* NIES-551 Ha5, *Cc. yangmingshanensis* NIES-4659 Ha1, and *Cz. merolae* 10D were harvested by centrifugation at 2,000 × *g* for 5 min at room temperature. The resulting cell pellets were resuspended in 50 μL of cell suspension buffer (10 mM Tris-HCl, pH 7.2, 50 mM EDTA, 20 mM NaCl). Next, 50 μL of 2% Certified Low Melt Agarose (Bio-Rad), prepared in 0.5× TBE buffer (44.5 mM Tris, 44.5 mM boric acid, 1 mM EDTA at pH 8.0), was added to each suspension, gently mixed, and poured into disposable plug molds (Bio-Rad) to solidify at room temperature. The solidified agarose gels (plugs) were incubated in *N*-lauroylsarcosine-Na buffer (1% *N*-lauroylsarcosine-Na, 500 mM EDTA, pH 8.0) with 5 mg proteinase K (Sigma-Aldrich) at 50 °C overnight. The plugs were washed with 0.5×TBE buffer before PFGE. For each sample, 2 mm of plugs were loaded into the wells of 1% PFGE certified agarose (Bio-Rad) gels prepared in 0.5×TBE buffer. Electrophoresis was run at 6 V cm^-1^ at 14°C with 120° pulse angle for 15 h with a switch time of 60 s and followed 9 h at a switch time of 90 s using the CHEF-DR III (Bio-Rad) system. Gels were stained with GelRed Nucleic Acid Gel Stain (Biotium), and signals were detected using a ChemiDoc Touch Imaging System (Bio-Rad).

### Preparation of linear DNA for the transformation

The sequences of the primers and all plasmids used in this study are listed in Supplementary Data Set 6. PCR amplification for plasmid construction and purification of PCR products were performed using PrimeSTAR Max DNA Polymerase (Takara Bio Inc., Japan) and the QIAquick PCR Purification Kit (QIAGEN, Venlo, Netherlands), respectively.

For the transformation of *Cc. yangmingshanensis* NIES-4659 Ha1, we constructed the following plasmids. To integrate the *Venus* expression cassette and the chloramphenicol acetyltransferase (*CAT*) selectable marker into the intergenic region between the g478.t1 and g479.t1 loci (IG1) through homologous recombination, plasmid pCcIG1-Venus-CAT was constructed (Fig. 5B). The *Venus* open reading frame (ORF; the nucleotide sequence was codon-optimized for the *Cz. merolae* nuclear genome) and the *CAT* ORF from *Staphylococcus aureus* (UniProtKB/Swiss-Prot ID: P00485; the nucleotide sequence was codon-optimized for the *Cz. merolae* nuclear genome) were commercially synthesized. The *mVenus* ORF was connected with the *Cc. yangmingshanensis APCC* promoter (*pAPCC*; 500 bp upstream of the *APCC* ORF) and the *Cc. yangmingshanensis β-TUBULIN* terminator (*tTUBB*; 250 bp downstream of the *β-TUBULIN* ORF). The *CAT* ORF was connected with the *Cc. yangmingshanensis ELONGATION FACTOR 1α* promoter (*pEF1α*; 500 bp upstream of the *EF1α* ORF) and the *Cc. yangmingshanensis UBIQUITIN* terminator (*tUBQ*; 250 bp downstream of the *UBIQUITIN* ORF).

To generate a *Cc. yangmingshanensis* transformant in which the *CAT* selectable marker and the herpes simplex virus thymidine kinase (*HSVtk*) suicide marker, linked by a Glycine-Serine (GS) linker, are integrated into the IG1 locus, plasmid pCcIG1-tUBQ-CAT-HSVtk-tUBQ was constructed (Fig. 5G).

The *HSVtk* ORF (UniProtKB/Swiss-Prot ID: P13157; the nucleotide sequence was codon-optimized for the *Cz. merolae* nuclear genome) was commercially synthesized. The *CAT-HSVtk* fusion ORF was connected with *pEF1α*. To generate a uracil-auxotrophic (Δ*URA5.3*) chloramphenicol-resistant *Cc. yangmingshanensis* transformant (Fig. 6A), plasmid pCcΔURA5.3-Venus-CAT was constructed (Supplementary Fig. S6A).

For the transformation of *Cd. caldarium* NIES-551 Ha5, we constructed the following plasmids. To integrate the *Venus* expression cassette and the blasticidin S deaminase (*BSD*) selectable marker into the intergenic region between the g620.t1 and g621.t1 loci (IG1) through homologous recombination, plasmid pCdIG1-Venus-BSD was constructed (Supplementary Fig. S5A). The *mVenus* ORF (codon-optimized for the *Cd. caldarium* nuclear genome) and the *BSD* ORF from *Aspergillus terreus* (UniProtKB/Swiss-Prot ID: P0C2P0.1; codon-optimized for the *Cd. caldarium* nuclear genome) were commercially synthesized. The *mVenus* ORF was connected with the *Cd. caldarium APCC* promoter (*pAPCC*; 500 bp upstream of the *APCC* ORF) and the *Cd. caldarium β-TUBULIN* terminator (*tTUBB*; 250 bp downstream of the *β-TUBULIN* ORF). The *BSD* ORF was connected with the *Cd. caldarium ELONGATION FACTOR 1α* promoter (*pEF1α*; 500 bp upstream of the *EF1α* ORF) and the *Cd. caldarium UBIQUITIN* terminator (*tUBQ*; 250 bp downstream of the *UBIQUITIN* ORF). To generate a uracil-auxotrophic (Δ*URA5.3*), blasticidin S-resistant *Cd. caldarium* transformant (Fig. S8), plasmid pCdΔURA5.3-Venus-CAT was constructed (Supplementary Fig. S6A).

For the transformation of *Cz. merolae* 10D, we constructed the following plasmids. To generate a *Cz. merolae* transformant in which the *Venus* expression cassette, the *CAT* selectable marker, and the *HSVtk* suicide marker are integrated into the intergenic region (IG1) upstream of the *URA5.3* gene locus, plasmid pCzURAup-Venus-HSVtk-CAT-URAup was constructed (Supplementary Fig. S5G). The *mVenus* ORF was connected with the *Cz. merolae APCC* promoter (*pAPCC*; 600 bp upstream of the *APCC* ORF) and the *Cz. merolae β-TUBULIN* terminator (*tTUBB*; 197 bp downstream of the *β-TUBULIN* ORF). The *HSVtk* ORF was connected with the *Cz. merolae CAB* promoter (*pCAB*; 589 bp upstream of the *CAB* ORF) and the *Cz. merolae APX* terminator (*tAPX*; 420 bp downstream of the *APX* ORF). The *CAT* ORF, fused to a sequence encoding a chloroplast transit peptide (the N-terminal 60 amino acids of *Cz. merolae APCC*), was connected with the *Cz. merolae CPCC* promoter (*pCPCC*; 500 bp upstream of the *CPCC* ORF) and the *Cz. merolae UBIQUITIN* terminator (*tUBQ*; 275 bp downstream of the *UBIQUITIN* ORF). To generate a uracil-auxotrophic (Δ*URA5.3*), chloramphenicol-resistant *Cz. merolae* transformant, plasmid pCzΔURA5.3-Venus-CAT was constructed (Supplemental Figure S6A). The *Venus* ORF was connected with the *Cz. merolae APCC* promoter (*pAPCC*; 500 bp upstream of the *APCC* ORF) and the *Cz. merolae β-TUBULIN* terminator (*tTUBB*; 250 bp downstream of the *β-TUBULIN* ORF). The *CAT* ORF was connected with the *Cz. merolae ELONGATION FACTOR 1α* promoter (*pEF1α*; 500 bp upstream of the *EF1α* ORF) and the *Cz. merolae UBIQUITIN* terminator (*tUBQ*; 250 bp downstream of the *tUBQ* ORF).

Linear DNA used for transformation was prepared by PCR with the primer set pUC19F and pUC19R (designed based on the pUC19 sequence), using the respective plasmids as templates. PCR amplification was performed using PrimeSTAR Max DNA Polymerase (Takara Bio Inc.). PCR products were purified using the QIAquick PCR Purification Kit (QIAGEN) and eluted with 55 μL of sterile water, of which 45 μL was used for transformation.

### Transformation

PEG-mediated transformation of haploid clones of *Cc. yangmingshanensis* NIES-4659 Ha1, *Cd. caldarium* NIES-551 Ha5, and *Cz. merolae* 10D was performed according to Fujiwara and Ohnuma (2018), with the following modifications.

To prepare cells for transformation, haploid cells of *Cc. yangmingshanensis* NIES-4659 Ha1 and *Cz. merolae* 10D were inoculated into 20 mL of MA medium at pH 2.0, while haploid cells of *Cd. caldarium* NIES-551 Ha5 were inoculated into 20 mL of MA medium at pH 0.75, each in a 25-cm² tissue culture flask, to give an OD_750_ of 0.4. The cells were then cultured at 42 °C (for *Cc. yangmingshanensis* and *Cz. merolae*) or 37 °C (for *Cd. caldarium*) under a 12-h light/12-h dark cycle (50 μmol photons m⁻² s⁻¹) on a reciprocal shaker at 140 rpm (NR-2; TAITEC). At the end of the third light period, when the percentage of S-phase cells reached its peak (Fujiwara et al., 2020; Jong et al., 2021), Tween-20 was added to the culture to a final concentration of 0.001%. The cells were harvested by centrifugation at 2,000 × *g* for 5 min at room temperature. The cells were then resuspended in IMA medium at pH 1.2 (or pH 0.75 for *Cd. caldarium*) to adjust the OD_750_ to 100. To prepare ∼60% (wt/vol) PEG solution, 0.3 g of PEG4000 (Sigma-Aldrich) was dissolved in 225 μL of MA2 medium (Ohnuma et al., 2008) at 95 °C for 5 min and then kept at 42 °C on a heat block until use. Next, 45 μL of DNA solution (3–5 μg of linear DNA), 5 μL of 10× transformation solution (400 mM [NH_4_]_2_SO_4_, 40 mM MgSO_4_, 0.3% H_2_SO_4_), and 62.5 μL of PEG solution were mixed by pipetting in a 1.5-mL tube, yielding a total volume of 112.5 μL. Then, 12.5 μL of cell suspension was added to the 112.5 μL transformation-DNA-PEG mixture (final PEG concentration ∼30% [wt/vol]). The tube was vigorously inverted ten times, and the contents were immediately transferred to 10 mL of IMA medium at pH 1.2 (or pH 0.75 for *Cd. caldarium*) in one well of a 6-well plate (VTC-P6, VIOLAMO). For Δ*URA5.3* transformants, the media were supplemented with 0.5 mg/mL uracil (Sigma-Aldrich). The cells were cultured statically in a 2% CO₂ incubator at 42 °C (or 37 °C for *Cd. caldarium*) under continuous light (30 μmol photons m⁻² s⁻¹) for 3 days. After incubation, the cells were harvested by centrifugation at 2,000 × *g* for 5 min at room temperature. The cell pellet was resuspended in 1 mL of IMA medium at pH 1.2 (or pH 0.75 for *Cd. caldarium*). A 100-μL aliquot of the cell suspension was inoculated into 1 mL of IMA medium at pH 1.2 supplemented with 100 μg/mL CP (or IMA medium at pH 0.75 supplemented with 1 mg/mL BS for *Cd. caldarium*). For Δ*URA5.3* transformants, the media were supplemented with 0.5 mg/mL uracil and 5-FOA (FUJIFILM Wako). The cells were cultured statically in a 2% CO₂ incubator at 42 °C (or 37 °C for *Cd. caldarium*) under continuous light (30 μmol photons m^-2^ s^-1^) for 3 weeks to allow drug-resistant transformants to grow. Single clones were obtained by limiting dilution of the culture in a 96-well plate (92696, TPP Techno Plastic Products). The Δ*URA5.3* clones were maintained in MA medium at pH 2.0 (or pH 0.75 for *Cd. caldarium*) supplemented with 0.5 mg/mL uracil.

To remove the *HSVtk* suicide marker from a chromosome of a transformant through intrachromosomal homologous recombination, the cells in which the *HSVtk* suicide marker was integrated were inoculated into 1 mL of MA medium at pH 2.0 supplemented with 10 µg/mL 5-FdU (or 20 µg/mL for *Cz. merolae*), in one well of a 24-well plate, to give an OD₇₅₀ of 0.2. The cells were then cultured statically in a 2% CO₂ incubator at 42 °C under continuous light (30 μmol photons m⁻² s⁻¹) for 3 weeks to select for cells that had lost the *HSVtk* suicide marker. Single clones were obtained by limiting dilution in a 96-well plate or by plating on cornstarch slurry spots over gellan gum–solidified MA medium.

The occurrence of homologous recombination in each transformant clone was confirmed by PCR using genomic DNA as the template, with the primer sets indicated in Supplementary Data Set 6. PCR amplification was performed using KOD One PCR Master Mix (TOYOBO CO., LTD., Osaka, Japan).

### Immunoblotting

Cyanidiophycean cells were harvested by centrifugation at 2,000 × *g* for 5 min at room temperature. The cell pellets were lysed in sample buffer (2% SDS, 62 mM Tris-HCl, pH 6.8, 100 mM DTT, 10% glycerol, and 0.01% bromophenol blue) and incubated at 95°C for 5 min. The lysate was centrifuged at 15,000 × *g* for 10 min, and the supernatant was collected.

Protein concentration was determined using the XL-Bradford assay (Integrale, Japan). A total of 6 µg of protein was separated by SDS-PAGE and transferred to PVDF membranes (Immobilon; Millipore). Membrane blocking, antibody incubation, and signal detection were performed as described by Fujiwara *et al*. (2024).

### Mating experiments and organelle DNA inheritance analysis

To generate and select a heterozygous diploid clone by mating haploids, cells of a Δ*URA5.3* haploid clone and those of a wild-type haploid clone, or cells from a haploid population derived from a diploid clone, were inoculated into 1 mL of MA medium at pH 2.0 (or pH 0.75 for *Cd. caldarium*), supplemented with 0.5 mg/mL uracil, to give an OD_750_ of 0.1 for each (total OD_750_ = 0.2). The cells were cultured statically in one well of a 24-well plate, enclosed in an AnaeroPack CO_2_ generator (Mitsubishi Gas Chemical), in an incubator at 42 °C (or 37 °C for *Cd. caldarium*) under 12-h light/12-h dark conditions (50 μmol photons m⁻² s⁻¹) for 7 days (or 10 days for *Cd. caldarium*) to allow haploid cell mating.

For *Cc. yangmingshanensis* and *Cz. merolae*, the cells were harvested by centrifugation at 1,500 × *g* for 5 min and resuspended in 1 mL of MA medium at pH 2.0. Then, 100 μL of the cell suspension was spread onto an HATF Immobilon nitrocellulose membrane (85 mm; Merck Millipore) placed over gellan gum–solidified MA medium at pH 2.0 (9-cm Petri dish, 0.5% gellan gum) supplemented with 100 μg/mL CP (or 250 μg/mL for *Cz. merolae*). The cells were cultured in a 2% CO₂ incubator at 42 °C under continuous light (30 μmol photons m⁻² s⁻¹) for 2 weeks to generate colonies of heterozygous diploid cells, which exhibit CP resistance but not uracil auxotrophy (Fig. 6D; Supplementary Fig. S7A). Single colonies were picked and inoculated into 1.0 mL of MA medium at pH 2.0. For *Cd. caldarium*, instead of spreading the harvested cells onto an HATF Immobilon nitrocellulose membrane, the cells were resuspended in 1 mL of MA medium at pH 2.0, and 100 μL of the cell suspension was inoculated into 1 mL of MA medium at pH 2.0 supplemented with 1 mg/mL blasticidin S in each well of a 24-well plate. The cells were cultured in a 2% CO₂ incubator at 37 °C under continuous light (30 μmol photons m⁻² s⁻¹) for ∼1 month. Single clones were obtained by limiting dilution of the culture in a 96-well plate.

The heterozygosity of the resulting diploid clones was confirmed by PCR (primer sequences are listed in Supplementary Data Set 6). The heterozygous diploid clones were maintained in MA medium at pH 2.0.

To examine organelle DNA inheritance, DNA was extracted from parental haploid clones and their diploid progeny using the Kaneka Easy DNA Extraction Kit ver. 2 (Kaneka Corp., Tokyo, Japan) with acid-washed glass beads (425–600 μm; Sigma-Aldrich). Targeted regions of the mitochondrial and chloroplast DNA were amplified by PCR using specific primer sets (sequences are listed in Supplementary Data Set 6) and sequenced by Sanger sequencing.

### RNA extraction and RNA-seq analyses

Cells of *Cc. yangmingshanensis* NIES-4659 Ha1 and the homozygous diploid derived from NIES-4659 Ha1 were inoculated into 50 mL of MA medium at pH 2.0 to give an OD_750_ of 0.2 in 100-mL test tubes (IWAKI, Japan). The cells were then cultured with aeration (500 mL ambient air min^−1^) at 40°C under continuous light (100 μmol photons m^−2^ s^−1^) for 3 days.

Cells of other cyanidiophycean strains (*Cz. merolae* MS1, *Cz. merolae* MS1 Ha3, *Cd. caldarium* NIES-551*, Cd. caldarium* NIES-551 Ha5, *G. partita* NBRC 102759, and *G. partita* NBRC102759 N1) were inoculated into 20 mL of MA medium at pH 2.0 for *Cz. merolae*, pH 0.75 for *Cd. caldarium*, and pH 1.0 for *G. partita* to give an OD_750_ of 0.2 in 25-cm² tissue culture flasks. The cells were cultured on a reciprocal shaker at 120 rpm in an incubator at 42°C under ambient air and continuous light conditions (50 μmol photons m^−2^ s^−1^) for 3 days. The cells were harvested by centrifugation at 2,000 × *g* for 5 min, frozen in liquid nitrogen, and stored at −80°C until use. Total RNA was extracted from the frozen cells according to (Hirooka et al., 2022). Strand-specific cDNA libraries were constructed and sequenced by Novogene Co. Ltd. (Beijing, China) using a NovaSeq 6000 with 150-bp paired-end sequencing. The sequence reads were processed using Cutadapt ver. 3.1 (Martin, 2011) to remove low-quality ends (<QV30), adapter sequences, and reads shorter than 50 bp. The trimmed reads were used for subsequent analyses.

The RNA-seq paired-end reads were mapped to protein-coding genes using Bowtie2 ver. 2.3.4.1 (Langmead and Salzberg, 2012). Strand-specific read counts for each protein-coding gene were calculated using SAMtools ver. 1.8 (Li et al., 2009) with the -f 80 option, BEDtools ver. 2.17.0 bamToBed (Quinlan and Hall, 2010), and R ver. 3.2.3. The count data were then normalized between the diploid and haploid cells, and differentially expressed genes (DEGs) were identified using edgeR ver. 3.12.1 (Robinson et al., 2010), with the following criteria: FDR < 0.01, logCPM > 2, |logFC| > 2. Transcripts per million (TPM) values for each protein-coding gene were calculated according to (Li and Dewey, 2011).

### Chromatin immunoprecipitation and sequencing

Cells of *Cc. yangmingshanensis* NIES-4659 Ha1 and the homozygous diploid derived from NIES-4659 Ha1 were inoculated into 50 mL of MA medium at pH 2.0 to give an OD_750_ of 0.2 in 100-mL test tubes (IWAKI, Japan). The cells were cultured with aeration (500 mL ambient air min^-1^) at 40°C under continuous light (100 μmol photons m^-2^ s^-1^) for 3 days. 50 mL of cultured cells were fixed with 1% formaldehyde at 30°C for 10 min and quenched with 250 mM glycine on ice for 5 min. The fixed cells were harvested by centrifugation at 2,500 × *g* for 5 min and washed twice with ice-cold Tris-buffered saline (TBS). The cell pellets were resuspended in extraction buffer (20 mM Tris-HCl, pH 7.5, 150 mM NaCl, 1 mM EDTA, 1% Triton X-100, 0.1% sodium deoxycholate, 0.1% SDS, and a protease inhibitor cocktail [Complete Mini, EDTA-free; Roche Diagnostics]) to give an OD_750_ of 25 for haploid cells and 100 for diploid cells. The cells were disrupted using 0.5-mm low-alkaline glass beads (YGBLA05; Yasui Kikai, Osaka, Japan) in a Multi-Beads Shocker (Yasui Kikai) at 2,700 rpm for 20 cycles (1 min on, 1 min off) at 0°C. The lysates were transferred to a milliTUBE 1 mL AFA Fiber (Covaris) and sheared using a Covaris S2 Focused-ultrasonicator (Covaris) for 20 min (60 sec × 20) at 4–6°C, with a water level of 15, duty cycle of 5%, intensity of 4, and 200 cycles per burst. The lysates were centrifuged at 15,000 × *g* for 10 min, and the supernatant was collected, frozen in liquid nitrogen, and stored at –80°C until use. A 250-μL aliquot of the chromatin solution was diluted with an equal volume of extraction buffer and incubated with 2 μL of antibody using a Microtube Rotator (MTR-103, AS ONE, Japan) at maximum speed at 4°C overnight. For the antibodies, an anti-H3K27me3 polyclonal antibody (07-449; Merck) and an anti-histone H3 polyclonal antibody (ab1791; Abcam) were used. Antibody– chromatin mixtures were then incubated with 80 μL of Dynabeads Protein G (Invitrogen in Thermo Fisher Scientific, MA, USA), pre-washed twice with 0.5 mL of extraction buffer, using a Microtube Rotator at maximum speed at 4°C for 4 h. The beads were washed once with 1 mL of extraction buffer, five times with 1 mL of LiCl wash buffer (50 mM Tris-HCl, pH 7.5, 0.5 M LiCl, 1 mM EDTA, 1% IGEPAL CA-630, and 0.5% sodium deoxycholate), and once with 1 mL of 50TE buffer (50 mM Tris-HCl, pH 7.5, and 10 mM EDTA). DNA was eluted from the beads by adding 100 μL of 50TE buffer containing 1% SDS and reverse crosslinked at 65°C overnight. The DNA solutions were treated with 5 μL of PureLink RNase A (Invitrogen) for 1 h at 37°C, followed by treatment with 1 μL of Proteinase K (FUJIFILM Wako) for 1 h at 50°C. DNA samples were purified using the Monarch PCR & DNA Cleanup Kit (NEB) and eluted with 20 μL of elution buffer. DNA libraries were constructed and sequenced by Macrogen (Korea) using a NovaSeq 6000 system with 150-bp paired-end sequencing. The sequence reads were processed using Cutadapt ver. 3.1 (Martin, 2011) to remove low-quality ends (<QV30), adapter sequences, and reads shorter than 50 bp. The trimmed reads were used for subsequent analyses as ChIP-seq reads.

ChIP-seq reads were mapped to the reference genome sequence of *Cc. yangmingshanensis* NIES-4659 Ha1 using Bowtie2 ver. 2.3.4.1 (Langmead and Salzberg, 2012). Mapped reads were counted for each protein-coding gene using SAMtools ver. 1.8 (Li et al., 2009) and BEDtools ver. 2.17.0 (Quinlan and Hall, 2010). The count data were normalized between the diploid and haploid cells, and genes that are differentially enriched for H3K27me3 were identified using edgeR ver. 3.12.1 (Robinson et al., 2010), with the following criterion: |logFC| > 2. For visualization, TDF files were generated from BAM files using igvtools (Robinson et al., 2011), and visualized using IGV ver. 2.8.2 (Robinson et al., 2011).

### Proteome analysis

Cells of *Cc. yangmingshanensis* NIES-4659 Ha1 and the homozygous diploid derived from NIES-4659 Ha1 were inoculated into 50 mL of MA medium at pH 2.0 to give an OD₇₅₀ of 0.2 in 100-mL test tubes (IWAKI). The cells were then cultured with aeration (500 mL ambient air min⁻¹) at 40°C under continuous light (100 μmol photons m⁻² s⁻¹) for 3 days. The cells were harvested by centrifugation at 2,000 × *g* for 5 min, frozen in liquid nitrogen, and stored at –80°C until use. Proteins were extracted from the respective frozen cell samples, hydrolyzed to peptides, and separated and analyzed using an UltiMate 3000 RSLCnano LC system (Thermo Fisher Scientific) and a Q Exactive HF-X mass spectrometer (Thermo Fisher Scientific) by Kazusa Genome Technologies Inc. (Kisarazu, Japan). The identification of peptides and quantitative analysis of the obtained MS data (including cross-run normalization) were conducted using DIA-NN 1.8.1 (Demichev et al., 2020), with both precursor and protein FDRs < 0.01.

## Data availability

The nuclear, mitochondrial, and chloroplast genome sequences of *Cc. yangmingshanensis* NIES-4659 Ha1 and *Cd. caldarium* NIES-551 Ha5, as well as the associated raw genomic sequence reads (including those of additional related strains), have been deposited in the DNA Data Bank of Japan (DDBJ)/European Molecular Biology Laboratory (EMBL)/GenBank (BioProject accession nos. PRJDB19695 and PRJDB19696, respectively). The Illumina raw genomic sequence reads of *Cz. merolae* strains (10D, MS1, and others) have been deposited in DDBJ/EMBL/GenBank (BioProject accession no. PRJDB37388). The RNA-sequencing data of cyanidiophycean algae have been deposited in DDBJ/EMBL/GenBank (BioProject accession no. PRJDB36168). The ChIP-sequencing data of *Cc. yangmingshanensis* have been deposited in DDBJ/EMBL/GenBank (BioProject accession no. PRJDB36311). The Illumina raw genomic sequence reads of *Cc. yangmingshanensis* strains (CcyaKS-1, CcyaKS-5, CcyaKS-11, TS5, TS6, TH2, TH3, and TH12) have also been deposited in DDBJ/EMBL/GenBank (BioProject accession no. PRJDB19695). All other data are included in the manuscript and/or the supporting information.

## Supporting information

List of putative gamete fusion and meiotic tool genes in the genomes of cyanidiophycean strains.

Comparison of growth rates under various conditions in cyanidiophycean strains.

Gene models for Cz. merolae 10D (General Transfer Format [GTF]), including newly identified gene models (CMX013C-CMX231C).

OrthoFinder output for four genera of Cyanidiophyceae.

Numbers of protein-coding genes and genome sizes in various eukaryotes.

Nucleotide sequences of primers and synthetic DNAs used in this study.

RNA-seq, DIA proteomic, and ChIP-seq data of nuclear genome-encoded protein-coding genes in Cc. yangmingshanensis haploid clone NIES-4659 Ha1 and the

RNA-seq data of nuclear genome-encoded protein-coding genes in Cz. merolae original diploid clone MS1 and haploid clones MS1 Ha3 and 10D.

RNA-seq data of nuclear genome-encoded protein-coding genes in Cd. caldarium original diploid clone NIES-551 and haploid clone NIES-551 Ha5.

RNA-seq data of nuclear genome-encoded protein-coding genes in G. partita original diploid clone NBRC 102759 and haploid clone NBRC 102759 N1.

## Acknowledgements

We thank Drs. Tsuneyoshi Kuroiwa, Haruko Kuroiwa, Kan Tanaka, Hisayoshi Nozaki, Osami Misumi, Hirofumi Yoshikawa, Yukihiro Kabeya, Kohta Yoshida, Takehiko Kanazawa and Yusuke Kobayshi for kind encouragement and advice on this work; Dr. Takeshi Itabashi for valuable advice on TEM observation; Drs. Koichi Hori, Ryoma Kamikawa and Motomichi Matsuzaki for kind advice on genomic studies; Drs. Yamato Yoshida and Sousuke Imamura for helpful advice on *Cz. merolae* research; Drs. Frédéric Berger and Daniel Schubert for their valuable advice on epigenetic studies. We also thank T. Ichinose, K. Saitoh, N. Shigenobu, S. Oda, K. Hashimoto, R. Ujigawa, and U. Sugimoto for their technical support. This work was supported by Grants-in-Aid for Scientific Research (23K05882 to S.H., and 24H00579 to S.-y.M.) from the Japan Society for the Promotion of Science, and JST-MIRAI Program (grant no. JPMJMI22E1 to S.-y.M.) from the Japan Science and Technology Agency.

## Author contributions

Conceptualization: S.H., T.F. and S.-y.M.; data curation: S.H.; formal analysis: S.H., S.I., and S.Y.; investigation: S.H., T.F., M.S., S.Y., D.T., R. Onuma., Y.K., S.W., Y.H., R. Ohbayashi., M.T., B.Z., R.T., F.Y., P.L., A.H.I and S.-y.M.; methodology: S.H., T.F., S.I., A.H.I and S.-y.M.; project administration: S.H. and S.-y.M.; resources: M.S. and P.L.; visualization: S.H., T.F. and S.-y.M.; writing—original draft: S.H. and S.-y.M; Writing—review & editing: All authors.

---

**Supplementary Figure S1.**
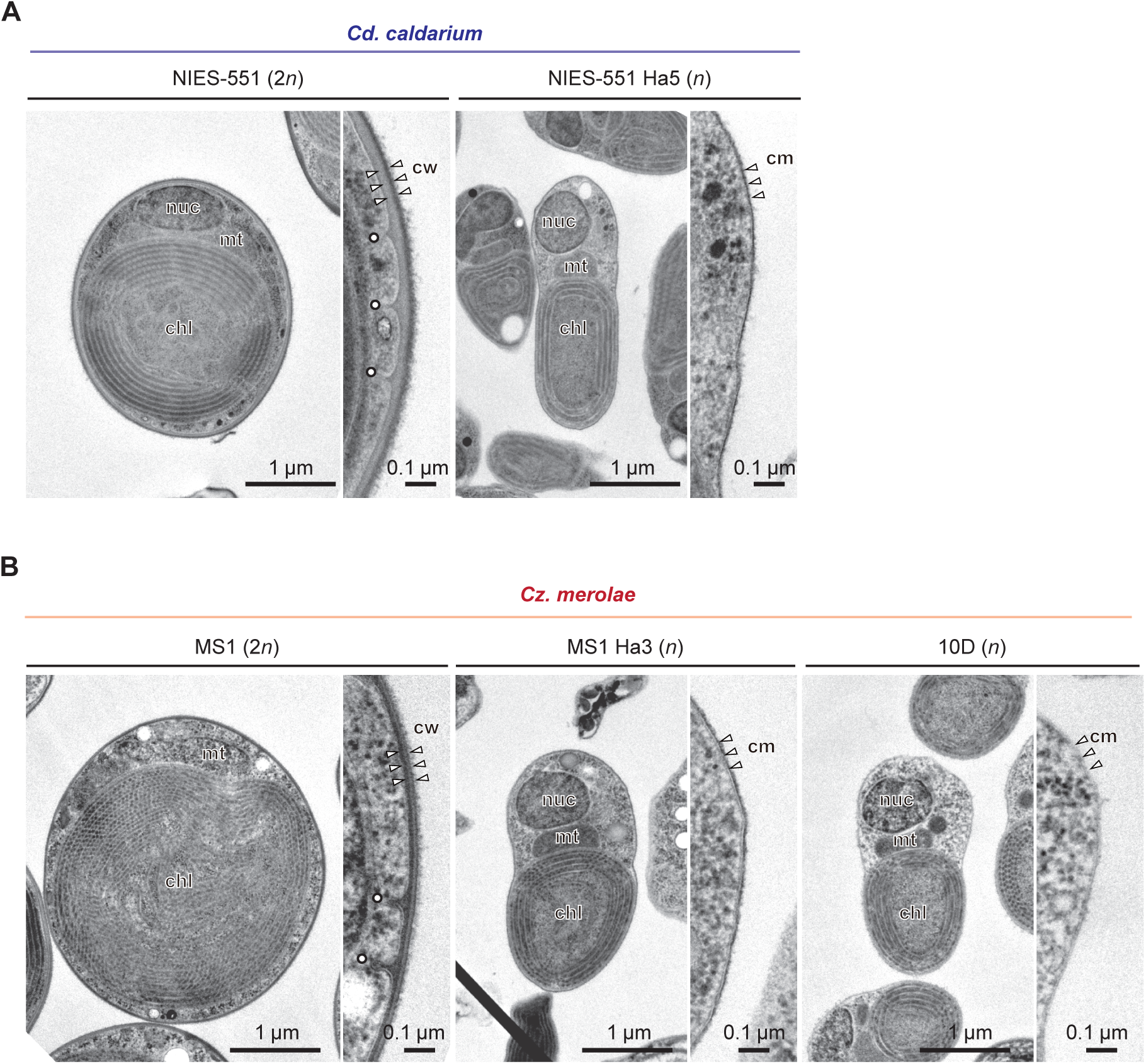
Transmission electron micrographs of *Cd. caldarium* and *Cz. merolae.* **A)** Transmission electron micrographs of the *Cd. caldarium* original 2*n* clone NIES-551 and *n* clone NIES-551 Ha5. **B)** Transmission electron micrographs of the *Cz. merolae* original 2*n* strain MS1 and *n* clones MS1 Ha3 and 10D. *chl*, chloroplast; *cm*, cell membrane; *cw*, daughter cell wall; *mt*, mitochondrion; *nuc*, nucleus; white dots, eisosomes.

**Supplementary Figure S2.**
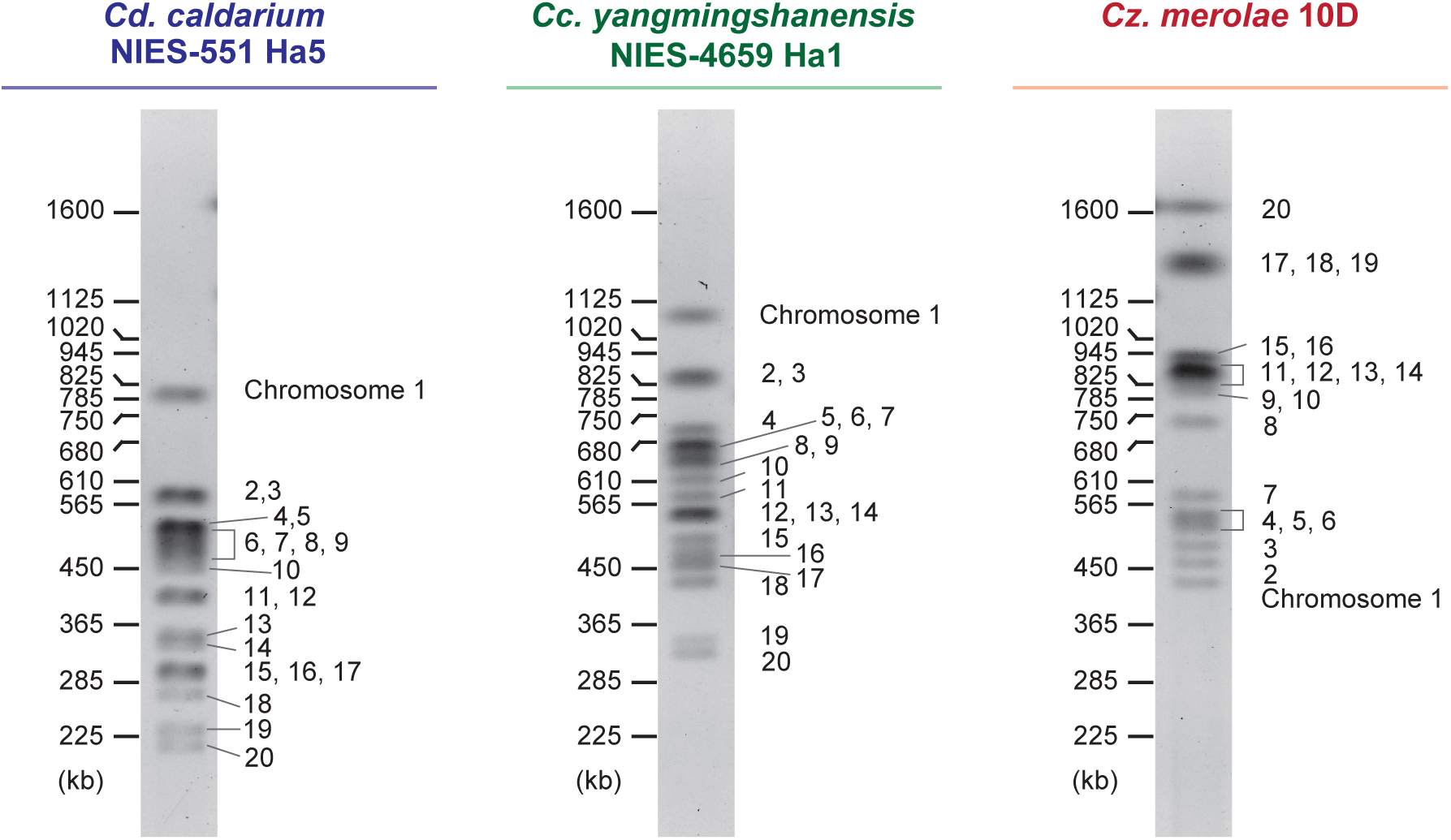
Pulsed-field gel electrophoresis separation of chromosomal DNA from haploid clones of *Cd. caldarium*, *Cc. yangmingshanensis*, and *Cz. merolae*.

**Supplementary Figure S3.**
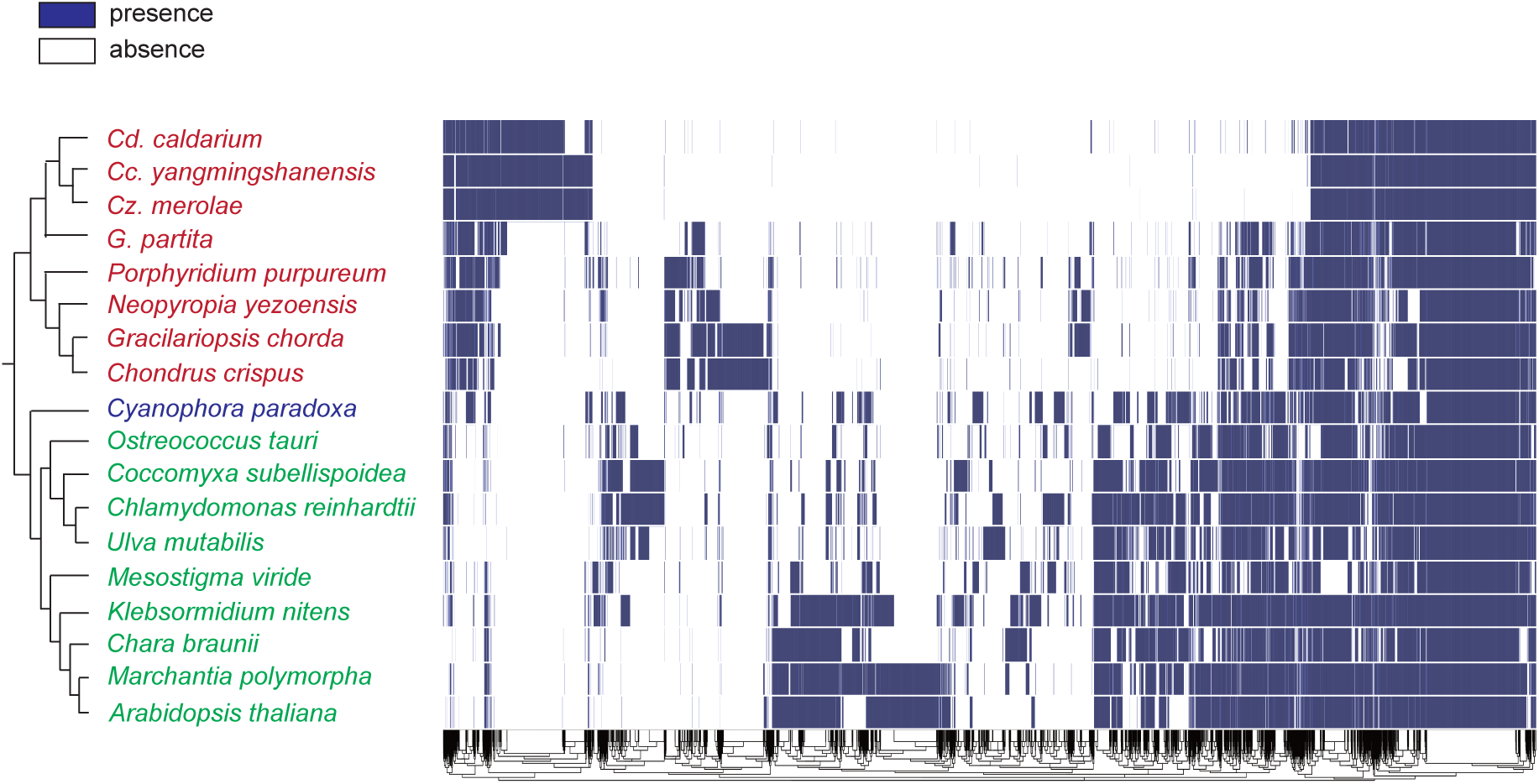
Binary heat map showing the distribution of 12,468 orthologous groups in 18 members of Archaeplastida, including Cyanidiophyceae. The orthologous groups were identified using OrthoFinder (Emms and Kelly, 2019), and species-specific orthologous groups were excluded. The clustering of orthologous groups was performed using the pheatmap R package ver. 1.0.12 (Kolde, 2019) with default parameters. Columns represent orthologous groups, while rows represent species. The dendrogram on the left side of the species names is based on previous studies (Munoz-Gomez et al., 2017; One Thousand Plant Transcriptomes, 2019). The presence of orthologous genes is shown in blue, and the absence is shown in white.

**Supplementary Figure S4.**
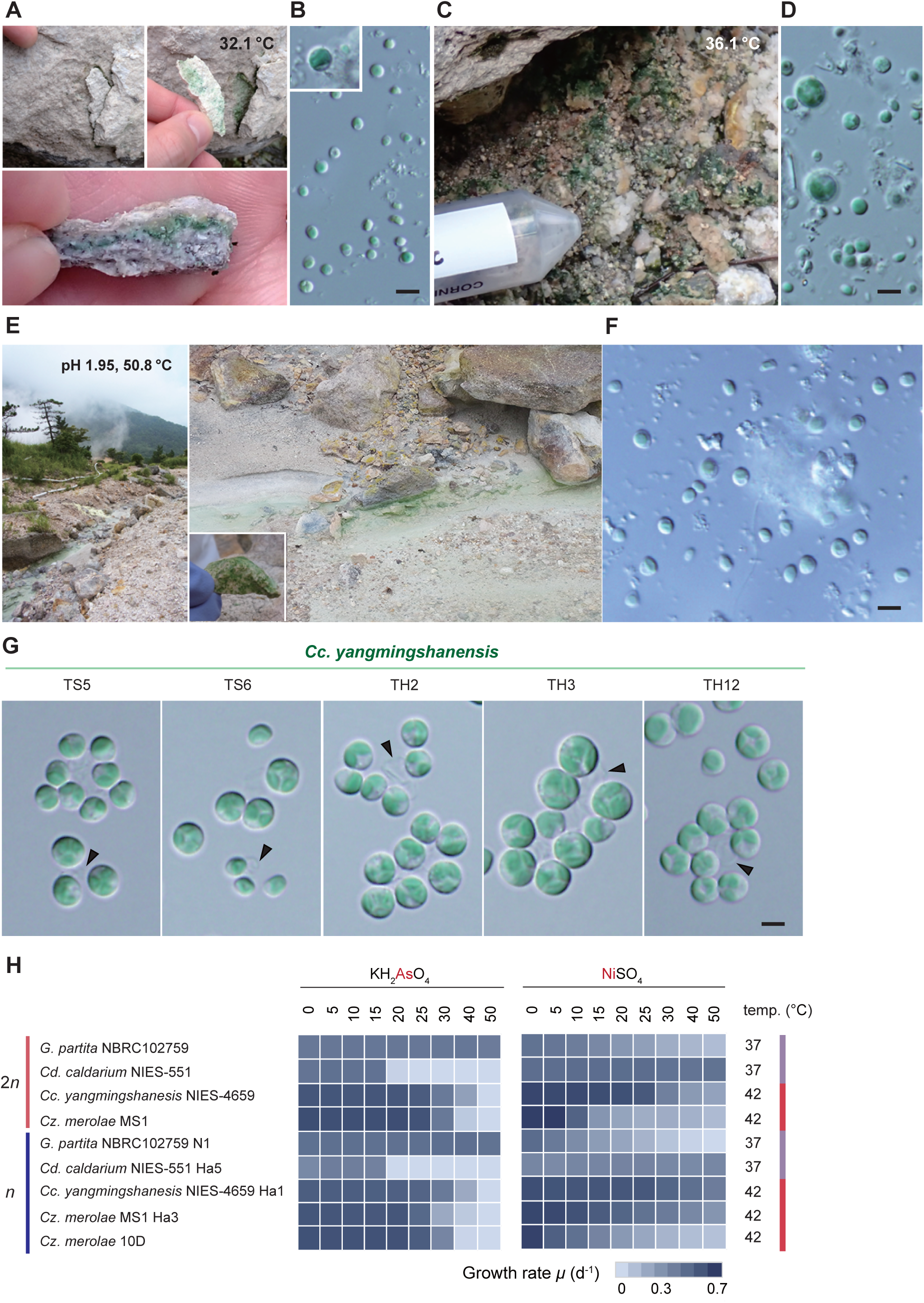
Natural habitat and comparison of metal tolerance of cyanidiophycean algae. **A)** Cyanidiophycean algae inhabiting the endolithic region around an acidic hot spring in Kusatsu, Gunma Prefecture, Japan (only temperature was measured because pH could not be determined). **B)** A DIC micrograph of cell-walled cyanidiophycean algae (smaller algal cells are most probably *Cyanidiococcus* spp.) obtained from an endolithic sample. The inset shows a larger algal cell, rarely observed, that is most probably *Galdieria*. Scale bar: 5 µm. **C)** Cyanidiophycean algae inhabiting the ground above spring water around an acidic hot spring in Kusatsu (only temperature was measured because pH could not be determined). **D)** A DIC micrograph of cell-walled cyanidiophycean algae (smaller algal cells are most probably *Cyanidiococcus* spp. and larger algal cells *Galdieria* spp.) obtained from the ground. Scale bar: 5 µm. **E)** Cyanidiophycean algae inhabiting an acidic hot spring in Tsukahara, Oita Prefecture, Japan, were predominantly found in association with stones in the water. **F)** A DIC micrograph of cell-walled cyanidiophycean algae (all algal cells are most probably *Cyanidiococcus* spp.) obtained from blue-green mats collected in the acidic hot spring. Scale bar: 5 µm. **G)** DIC micrographs of cultured cell-walled *Cc. yangmingshanensis* strains isolated from the acidic hot spring in Tsukahara. The white arrowhead indicates the mother cell wall released upon hatching of daughter cells. Scale bar: 2 µm. **H)** Heatmaps showing the growth rate based on the increase in OD_750_ under the indicated metal concentrations. See also Supplementary Data Set 2 for details.

**Supplementary Figure S5.**
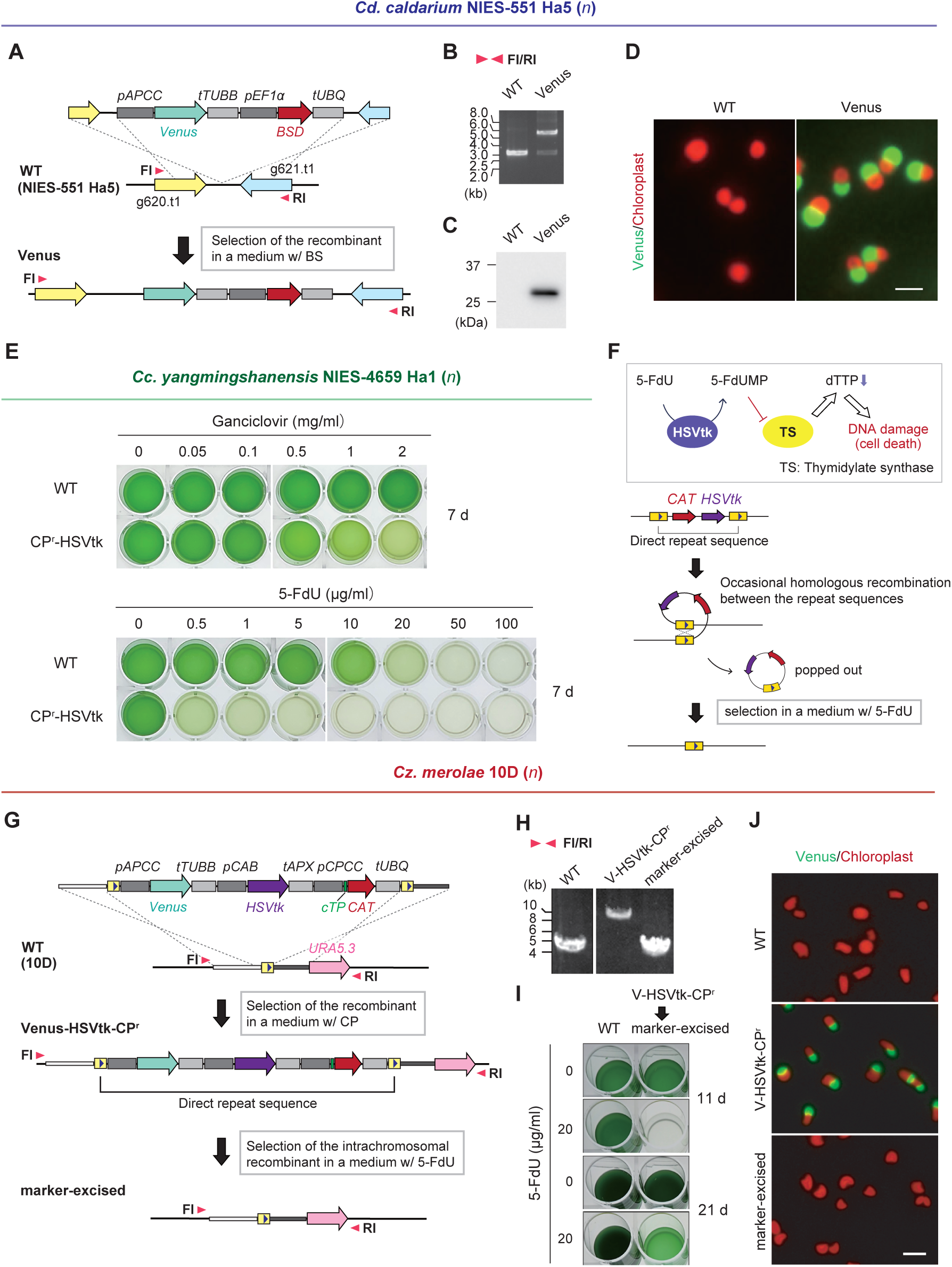
Genetic manipulation and marker removal in the cyanidiophycean algae. **A)** A schematic diagram showing insertion of the *Venus* expression cassette and the *BSD* selectable marker into a chromosomal intergenic region between the *g620.t1* and *g621.t1* loci by homologous recombination in *Cd. caldarium n* clone NIES-551 Ha5. **B)** Targeted integration of the transgenes into the chromosomal intergenic region was confirmed by PCR using primers FI and RI indicated by arrowheads in **A**. **C and D)** Expression of Venus protein was confirmed by immunoblotting with an anti–Green Fluorescent Protein (GFP) antibody **(C)** and by fluorescence microscopy **(D)** (green, Venus fluorescence; red, chloroplast fluorescence). WT served as a negative control. Scale bar: 2 µm. **E)** The CP^r^-HSVtk and WT *n* cells of *Cc. yangmingshanensis* were cultured for 7 days in the presence of the indicated concentrations of ganciclovir (upper) and 5-FdU (lower). **F)** HSVtk converts 5-FdU into the toxic product 5-FdUMP, which inhibits thymidylate synthase. Cells in which the selectable marker was removed through intrachromosomal homologous recombination between the repeat sequences (indicated by a blue arrowhead in the yellow boxes) can be selected in medium containing 5-FdU. **G)** A schematic diagram showing the targeted integration and subsequent removal of the *CAT* selectable marker in *Cz. merolae* 10D. The *Venus* expression cassette, the *HSVtk* suicide marker, and the *CAT* selectable marker were sandwiched between two directly repeated *URA5.3* upstream sequences (indicated by a blue arrow in the yellow boxes). This construct was integrated into an intergenic region (upstream of the *URA5.3* gene locus) of WT *Cz. merolae* 10D by homologous recombination. After selecting the transformant (V-HSVtk-CP^r^) in the presence of CP, the selectable marker was removed through intrachromosomal homologous recombination between the two copies of the repeated sequences, followed by selection with 5-FdU. **H)** The recombination events were confirmed by PCR using the primers FI and RI indicated by arrowheads in **G**. **I)** The V-HSVtk-CP^r^ cells were cultured for 21 days in the presence or absence of 5-FdU. WT *Cz. merolae* 10D served as a negative control. **J)** Expression of Venus was confirmed by fluorescence microscopy (green, Venus fluorescence; red, chloroplast fluorescence). Scale bar: 5 µm.

**Supplementary Figure S6.**
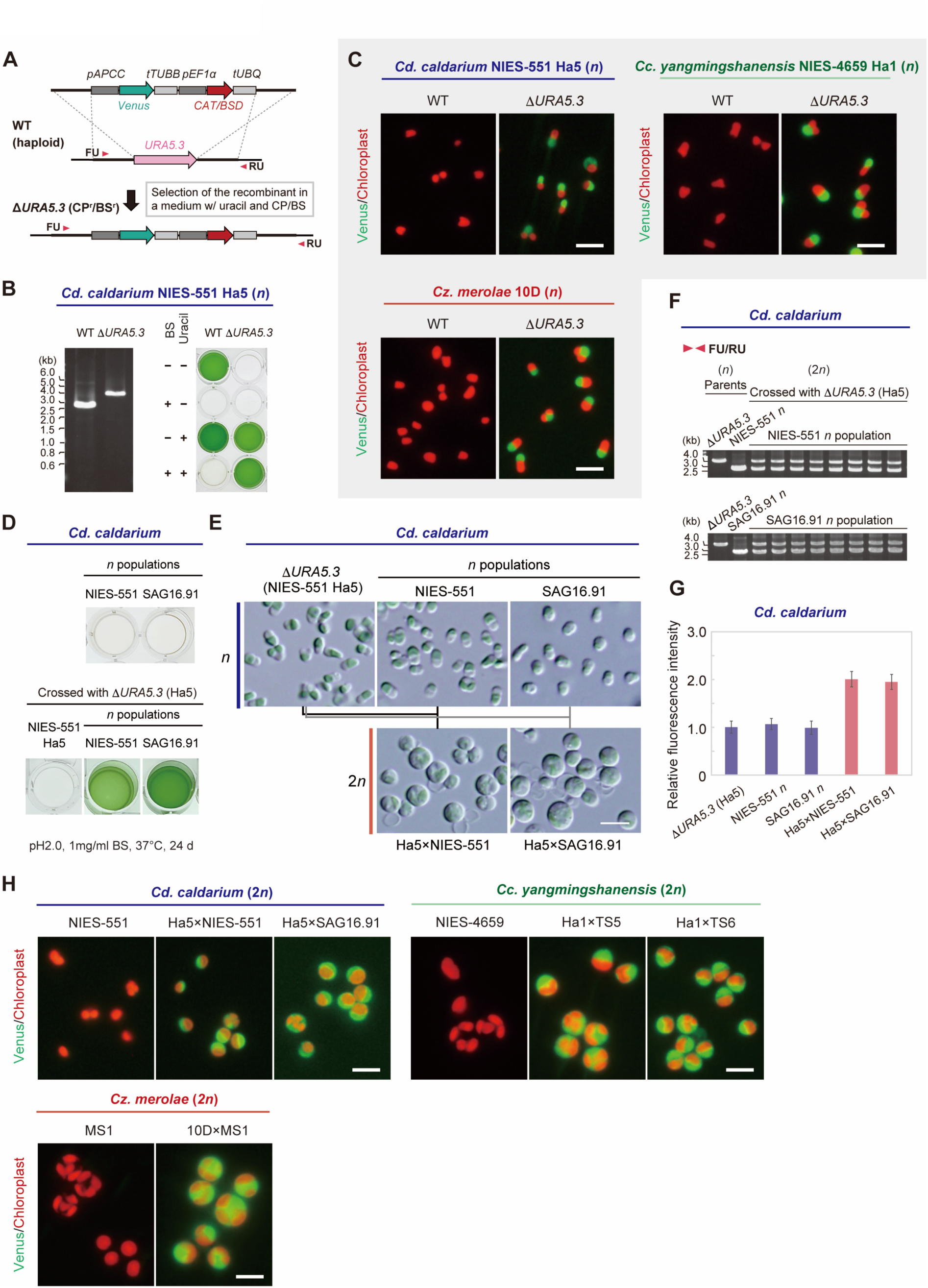
Construction of *URA5.3* knockout haploid strains and confirmation of Venus expression in *URA5.3* knockout haploid strains and hybrid diploid clones. **A)** To select heterozygous 2*n* clones, a uracil-auxotrophic, BS/CP-resistant *n* clone was generated. To knockout *URA5.3* and provide BS and CP resistance to *Cd. caldarium* NIES-551 Ha5 and *Cz. merolae* 10D, respectively, the Venus expression cassette and the *BSD* or *CAT* selectable marker were integrated into the chromosomal *URA5.3* locus by homologous recombination. **B)** Replacement of the chromosomal *URA5.3* locus with the Venus expression cassette and the *BSD* selectable marker in the resultant *Cd. caldarium* Δ*URA5.3 n* clone was confirmed by PCR using primers FU and RU, indicated by the arrowheads in **A**. WT *n* clone served as a control. The uracil auxotrophy and BS resistance of the Δ*URA5.3 n* clone were confirmed by cultivation for 7 days in the presence or absence of uracil and BS. WT *n* clone served as controls. **C)** Expression of Venus in the *URA5.3* knockout *n* clones was confirmed by fluorescence microscopy (green, Venus fluorescence; red, chloroplast fluorescence). Respective WT clones served as negative controls. Scale bar: 5 µm. **D)** *Cd. caldarium* Δ*URA5.3* (BS^r^) *n* clone was crossed with the wild-type NIES-551 clone Ha5 (the parental *n* clone of the Δ*URA5.3* [BS^r^] *n* clone) and with *n* populations derived from NIES-551 2*n* and SAG16.91 2*n*. Heterozygous 2*n* clones generated by mating were selected in MA liquid medium at pH 2.0 with 1 mg/ml BS. Respective WT *n* populations served as negative controls. The phylogenetic positions of *Cd. caldarium* NIES-551 and SAG16.91 are shown in Fig. 4E. **E)** DIC micrographs of *Cd. caldarium* clones: Δ*URA5.3* (NIES-551 Ha5) *n* clone, *n* populations derived from the original 2*n* clones NIES-551 and SAG16.91, and hybrid 2*n* clones Ha5×NIES-551 and Ha5×SAG16.91 from their respective combinations. **F)** Heterozygosity of the hybrid 2*n* clones was confirmed by PCR using primers FU and RU. **G)** Nuclear DNA content was compared by measuring fluorescence intensity of DAPI-stained nuclei. The mean fluorescence intensity of Δ*URA5.3 n* cells was defined as 1.0. Data are means ±SD from 20 independent cells. **H)** Venus expression in the hybrid 2*n* clones was confirmed by fluorescence microscopy (green, Venus fluorescence; red, chloroplast fluorescence). Respective WT 2*n* clones served as negative controls. Scale bar: 5 µm.

**Supplementary Figure S7.**
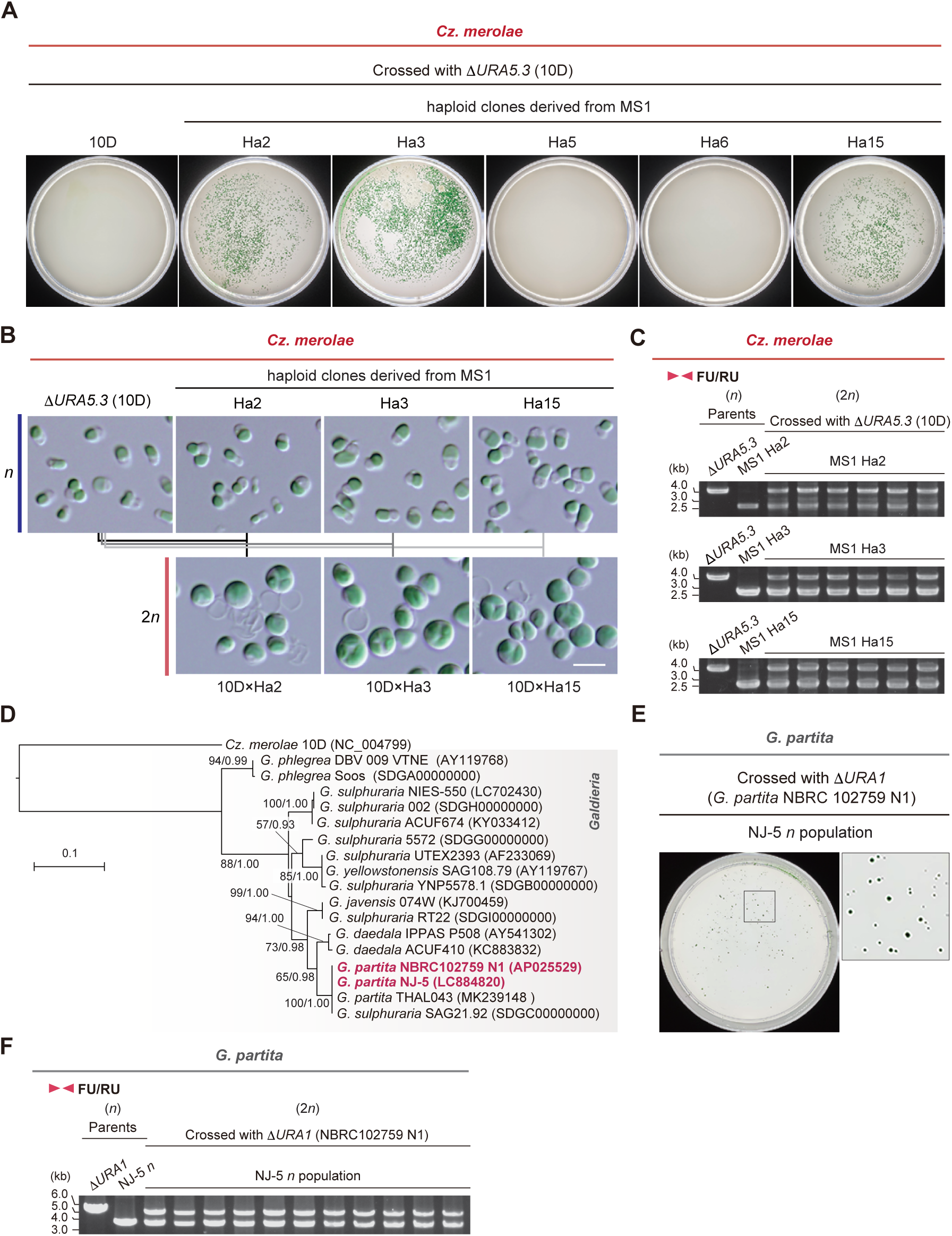
Heterozygous diploid generation by haploid mating in *Cz. merolae* and *G. partita.* **A)** Δ*URA5.3* (CP^r^) *n* clone of *Cz. merolae* 10D was crossed with *n* clones derived from the original 2*n* clone MS1, and heterozygous 2*n* clones generated by their mating were selected on an HATF Immobilon nitrocellulose membrane (85 mm) placed over gellan gum–solidified medium at pH 2.0 with CP. **B)** DIC micrographs of *Cz. merolae* clones: Δ*URA5.3* (10D) and *n* clones derived from the original 2*n* clone MS1, as well as hybrid 2*n* clones 10D×Ha2, 10D×Ha3, and 10D×Ha15 from their respective combinations. Scale bar: 5 µm. **C)** Heterozygosity of the hybrid 2*n* clones was confirmed by PCR using primers FU and RU, as indicated in Supplementary Figure S6A. **D)** Phylogenetic relationship of several *Galdieria* strains: The tree, based on nucleotide sequences of the chloroplast-encoded *rbcL*, was generated using maximum likelihood analysis (RAxML-NG ver. 1.0.3) (Kozlov et al., 2019). *Cz. merolae* 10D was used as an outgroup. BP >50% (left), obtained using ML, and BI >0.95 (right), calculated using Bayesian analysis (MrBayes ver. 3.2.7) (Ronquist et al., 2012), are indicated above the branches. Branch lengths reflect the evolutionary distances, as indicated by the scale bar. The strains used in this study are highlighted in red. **E)** *G. partita* NBRC 102759 N1 Δ*URA1* (BS^r^) *n* clone was crossed with the WT *n* population derived from the original 2*n* clone *G. partita* NJ-5, and heterozygous 2*n* clones were selected on MA gellan gum–solidified medium at pH 2.0 with 100 μg/ml BS. **F)** Heterozygosity of the hybrid 2*n* clones was confirmed by PCR using primers FU and RU.

**Supplementary Figure S8.**
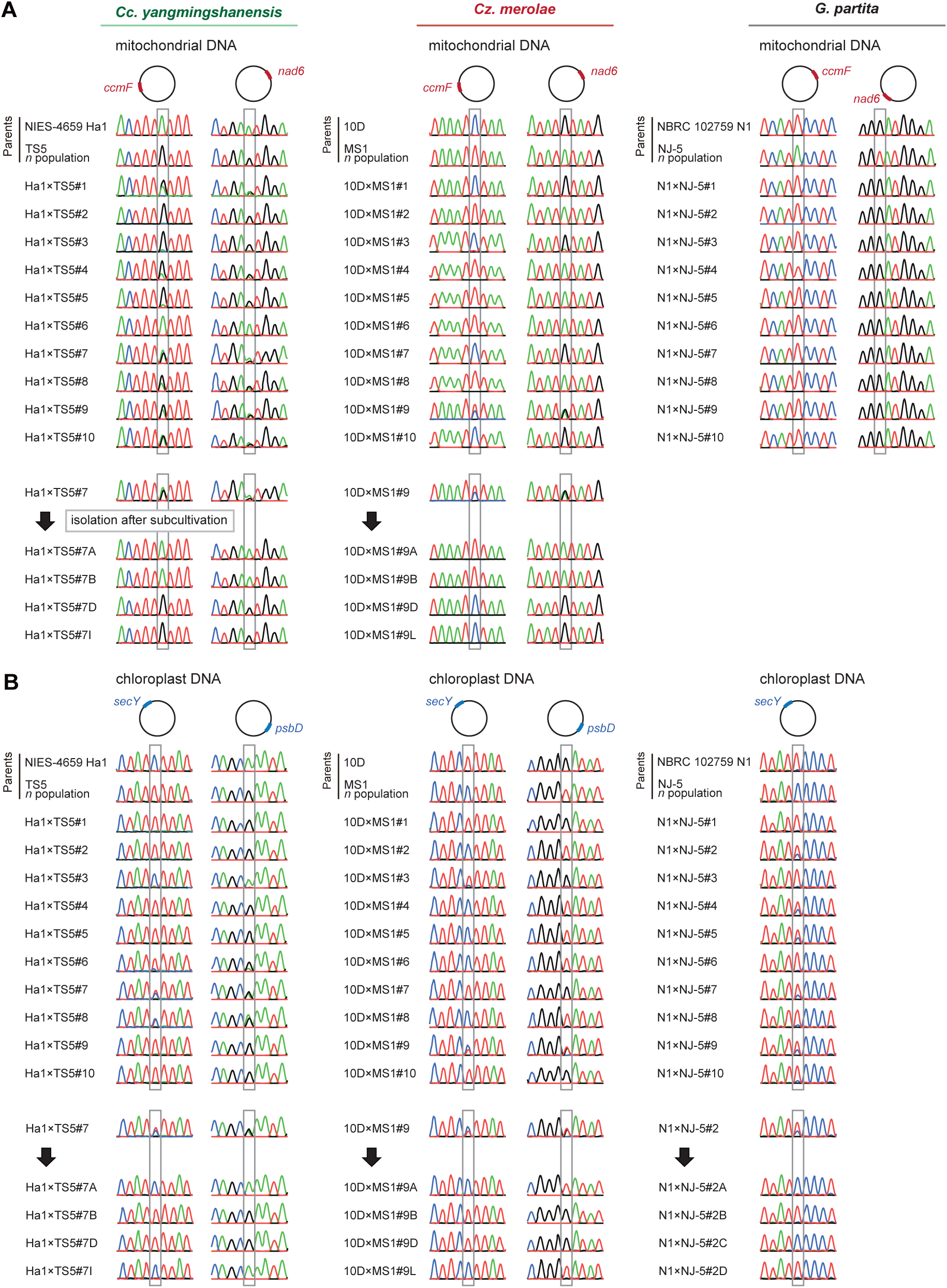
Organelle DNA inheritance in the cyanidiophycean algae. **A and B)** Sanger sequencing chromatograms of mitochondrial (*ccmF* and *nad6* loci) **(A)** and chloroplast (*secY* and *psaD* loci) **(B)** DNA regions obtained from parental *n* clones and their 2*n* progeny. Gray boxes highlight nucleotide variants between the parental *n* clones. Green = adenine, red = thymine, black = guanine, blue = cytosine. After subculturing the 2*n* clones Ha1×TS5#7 (*Cc. yangmingshanensis*), 10D×MS1#9 (*Cz. merolae*), and N1×NJ-5#2 (*G. partita*), which initially inherited mitochondrial and/or chloroplast DNA from both parents, single colonies were again isolated, and the target region of each colony was examined by Sanger sequencing.

**Supplementary Figure S9.**
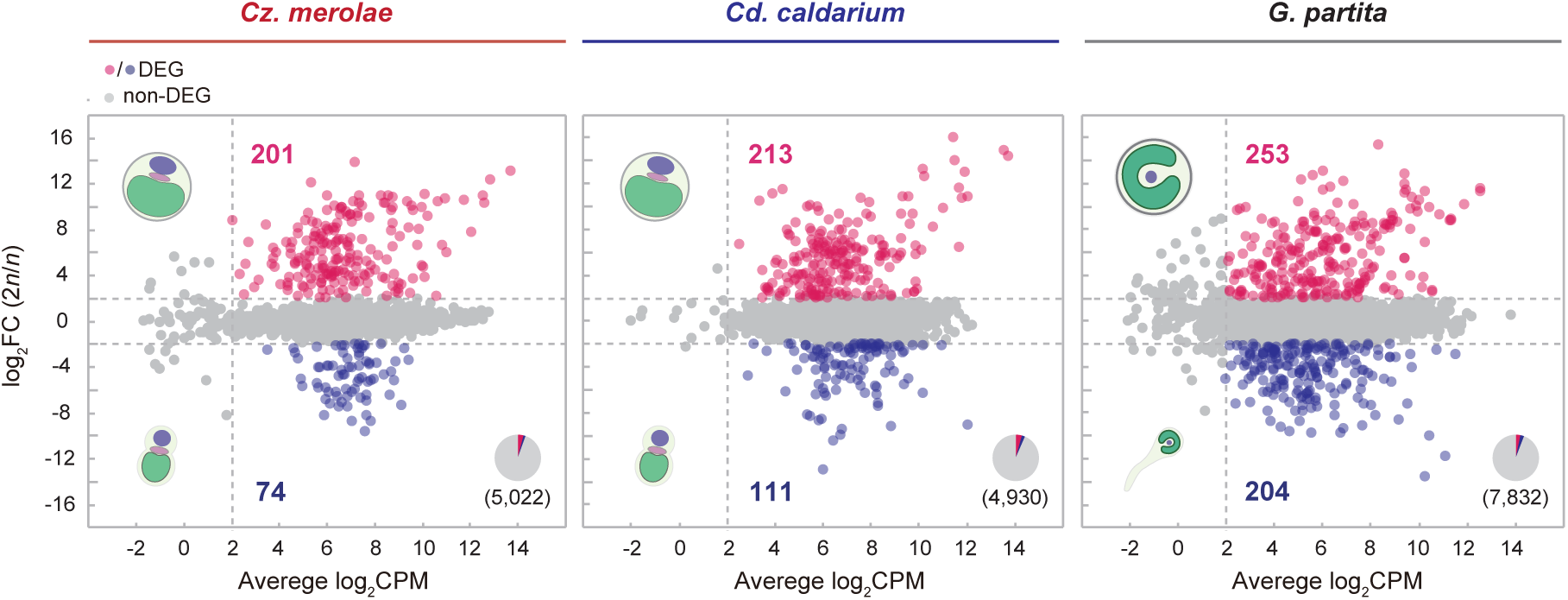
Comparison of transcriptomes between diploid and haploid phases in *Cz. merolae*, *Cd. caldarium*, and *G. partita*. These figures are identical to Figure 7A, except that the results are shown for *Cz. merolae*, *Cd. caldarium*, and *G. partita* rather than *Cc. yangmingshanensis*, and ChIP-seq of H3K27me3 was not conducted.

**Supplementary Figure S10.**
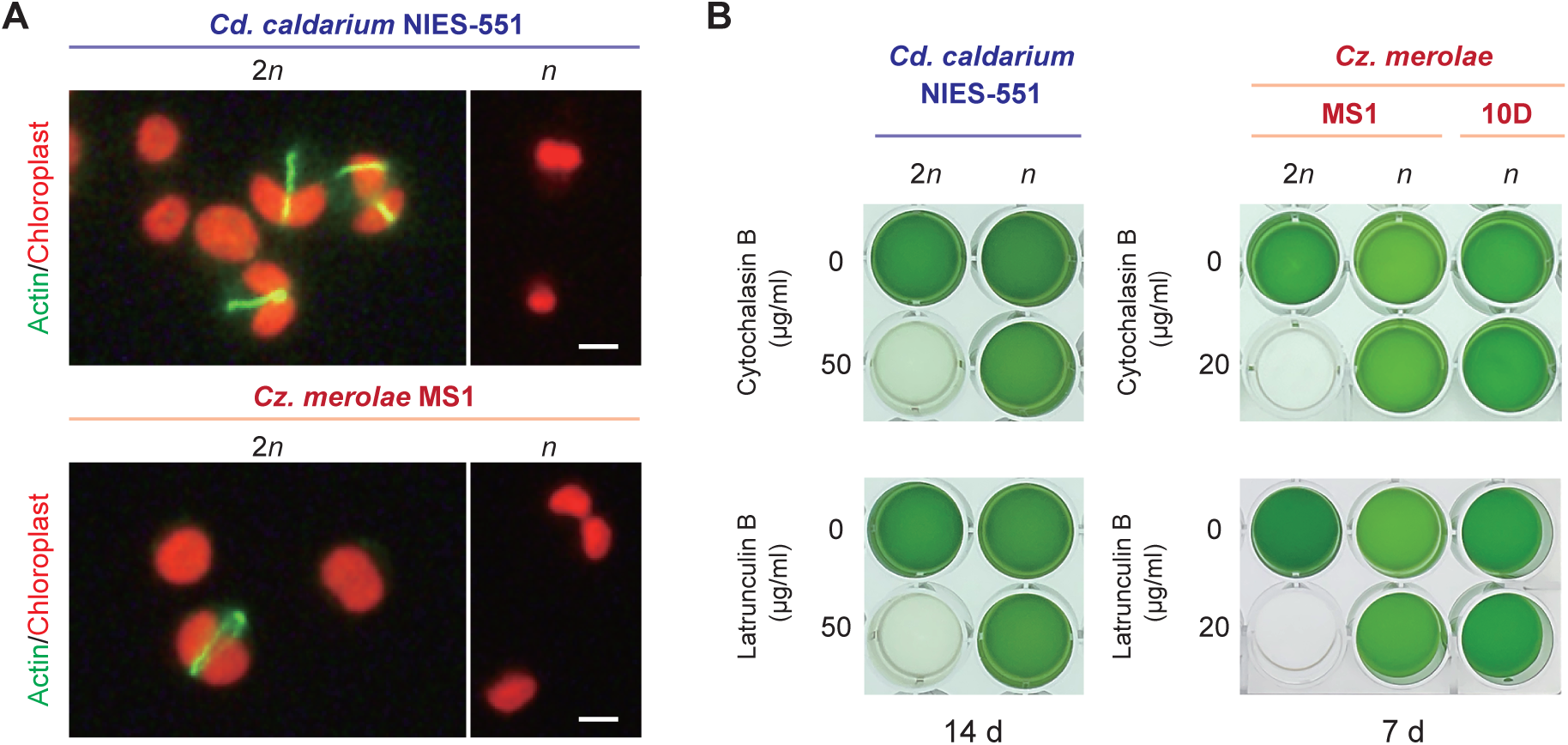
Comparison of sensitivity to actin polymerization inhibitors between diploid and haploid cells of *Cd. caldarium* and *Cz. merolae.* **A)** Actin filaments in 2*n* and *n* cells were visualized with Alexa Fluor^TM^ 488 Phalloidin and observed by fluorescence microscopy; green, Phalloidin fluorescence indicating actin filaments; red, chloroplast fluorescence. Scale bar: 2 µm. **B)** Photographs of the 2*n* and *n* cultures after 14 or 7 days of cultivation in the presence or absence of cytochalasin B or latrunculin B.

**Supplementary Figure S11.**
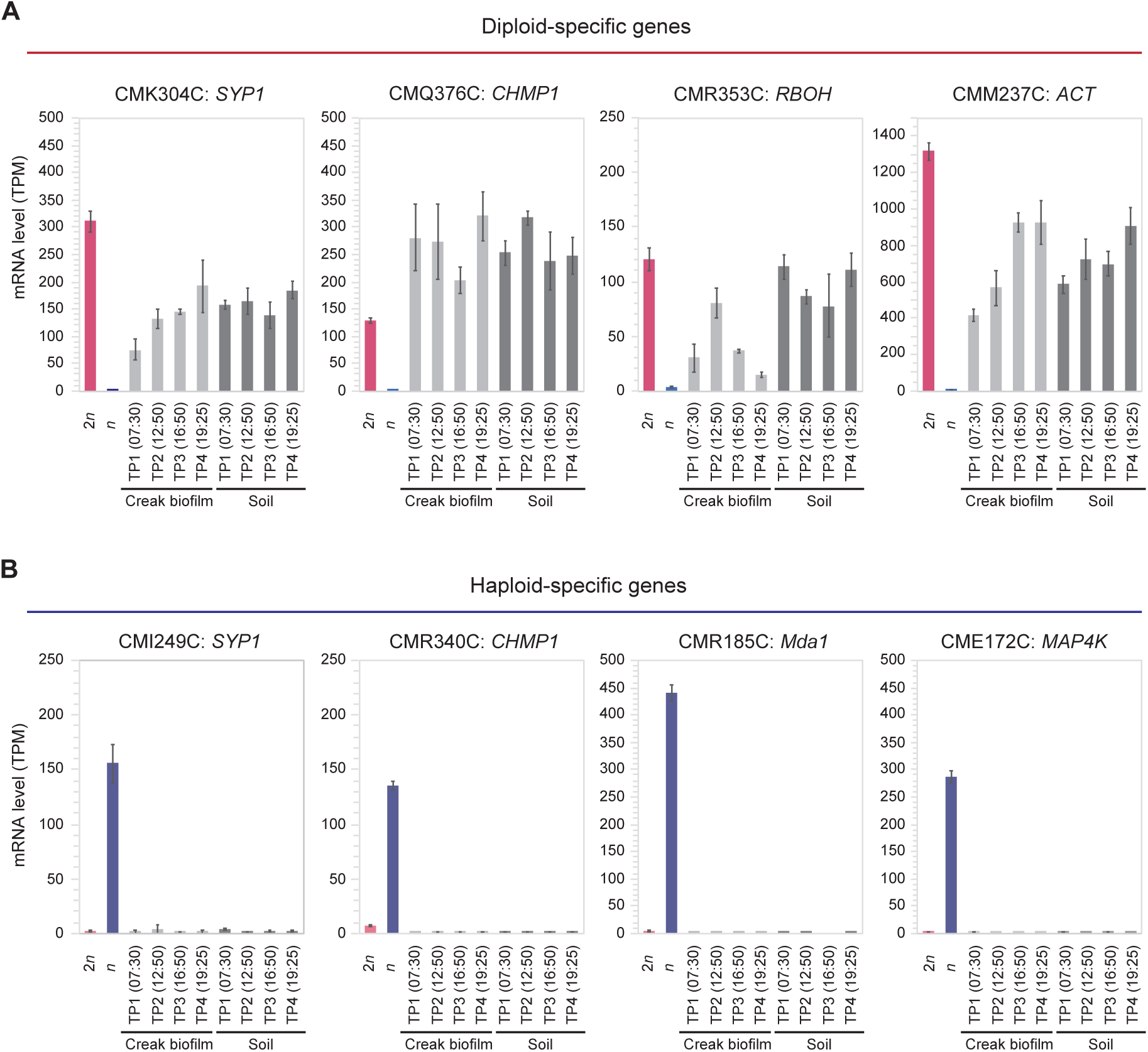
Reanalysis of metatranscriptome data from Yellowstone National Park, USA, showing expression patterns of diploid- and haploid-specific genes in *Cz. merolae*. The raw data were obtained from Stephens *et al*. (2024). **A and B)** mRNA levels (TPM) of diploid-specific **(A)** and haploid-specific genes **(B)** from metatranscriptome data of creek biofilms (aquatic) and soil environments (terrestrial; above the water surface) at four time points (TP1–TP4). TPM values of 2*n* and *n* cells were extracted from Supplementary Data Set 8. Data are means ± standard deviations (SDs) from three replicates.

**Supplementary Dataset S1.** List of putative gamete fusion and meiotic tool genes in the genomes of cyanidiophycean strains.

**Supplementary Dataset S2.** Comparison of growth rates under various conditions in cyanidiophycean strains.

**Supplementary Dataset S3.** Gene models for *Cz. merolae* 10D (General Transfer Format [GTF]), including newly identified gene models (CMX013C–CMX231C).

**Supplementary Dataset S4.** OrthoFinder output for four genera of Cyanidiophyceae.

**Supplementary Dataset S5.** Numbers of protein-coding genes and genome sizes in various eukaryotes.

**Supplementary Dataset S6.** Nucleotide sequences of primers and synthetic DNAs used in this study.

**Supplementary Dataset S7.** RNA-seq, DIA proteomic, and ChIP-seq data of nuclear genome–encoded protein-coding genes in *Cc. yangmingshanensis* haploid clone NIES-4659 Ha1 and the homozygous diploid clone derived from NIES-4659 Ha1.

**Supplementary Dataset S8.** RNA-seq data of nuclear genome–encoded protein-coding genes in *Cz. merolae* original diploid clone MS1 and haploid clones MS1 Ha3 and 10D.

**Supplementary Dataset S9.** RNA-seq data of nuclear genome–encoded protein-coding genes in *Cd. caldarium* original diploid clone NIES-551 and haploid clone NIES-551 Ha5.

**Supplementary Dataset S10.** RNA-seq data of nuclear genome–encoded protein-coding genes in *G. partita* original diploid clone NBRC 102759 and haploid clone NBRC 102759 N1.

